# A versatile and interoperable computational framework for the analysis and modeling of COVID-19 disease mechanisms

**DOI:** 10.1101/2022.12.17.520865

**Authors:** Anna Niarakis, Marek Ostaszewski, Alexander Mazein, Inna Kuperstein, Martina Kutmon, Marc E. Gillespie, Akira Funahashi, Marcio Luis Acencio, Ahmed Hemedan, Michael Aichem, Karsten Klein, Tobias Czauderna, Felicia Burtscher, Takahiro G. Yamada, Yusuke Hiki, Noriko F. Hiroi, Finterly Hu, Nhung Pham, Friederike Ehrhart, Egon L. Willighagen, Alberto Valdeolivas, Aurelien Dugourd, Francesco Messina, Marina Esteban-Medina, Maria Peña-Chilet, Kinza Rian, Sylvain Soliman, Sara Sadat Aghamiri, Bhanwar Lal Puniya, Aurélien Naldi, Tomáš Helikar, Vidisha Singh, Marco Fariñas Fernández, Viviam Bermudez, Eirini Tsirvouli, Arnau Montagud, Vincent Noël, Miguel Ponce de Leon, Dieter Maier, Angela Bauch, Benjamin M. Gyori, John A. Bachman, Augustin Luna, Janet Pinero, Laura I. Furlong, Irina Balaur, Adrien Rougny, Yohan Jarosz, Rupert W. Overall, Robert Phair, Livia Perfetto, Lisa Matthews, Devasahayam Arokia Balaya Rex, Marija Orlic-Milacic, Monraz Gomez Luis Cristobal, Bertrand De Meulder, Jean Marie Ravel, Bijay Jassal, Venkata Satagopam, Guanming Wu, Martin Golebiewski, Piotr Gawron, Laurence Calzone, Jacques S. Beckmann, Chris T. Evelo, Peter D’Eustachio, Falk Schreiber, Julio Saez-Rodriguez, Joaquin Dopazo, Martin Kuiper, Alfonso Valencia, Olaf Wolkenhauer, Hiroaki Kitano, Emmanuel Barillot, Charles Auffray, Rudi Balling, Reinhard Schneider, the COVID-19 Disease Map Community

## Abstract

The COVID-19 Disease Map project is a large-scale community effort uniting 277 scientists from 130 Institutions around the globe. We use high-quality, mechanistic content describing SARS-CoV-2-host interactions and develop interoperable bioinformatic pipelines for novel target identification and drug repurposing. Community-driven and highly interdisciplinary, the project is collaborative and supports community standards, open access, and the FAIR data principles. The coordination of community work allowed for an impressive step forward in building interfaces between Systems Biology tools and platforms. Our framework links key molecules highlighted from broad omics data analysis and computational modeling to dysregulated pathways in a cell-, tissue- or patient-specific manner. We also employ text mining and AI-assisted analysis to identify potential drugs and drug targets and use topological analysis to reveal interesting structural features of the map. The proposed framework is versatile and expandable, offering a significant upgrade in the arsenal used to understand virus-host interactions and other complex pathologies.

## 1. Introduction

The COVID-19 pandemic was and continues to be one of the most significant social and health challenges faced by humankind recently. The scientific community responded to these challenges with incredible resilience, adaptability, and eagerness to contribute. Large-scale community efforts emerged, and scientists from all over the world found new ways to connect and offer their skills to tackle the pandemic from various angles. The COVID-19 Disease Map project is a large-scale community effort to build an open-access, computable repository of COVID-19 molecular mechanisms—the COVID-19 Disease Map (C19DMap). The Map represents molecular and signaling pathways described in a broad range of the COVID-19 scientific literature in over forty diagrams compliant with systems biology standards. The content is based on human biocuration and supported by text mining solutions, such as INDRA (1) and AILANI (https://ailani.ai), and a plethora of tools and platforms for data integration, analysis, and computational modeling (2) (3).

In parallel with the content building, the community has also been developing an ecosystem of analytical and modeling pipelines that we aim to showcase here, extending the application use cases presented in our previous report (2). The pipelines mentioned above are developed either *de novo* or adapted to suit the high-quality mechanistic content of the C19DMap. The workflows aim to identify actionable targets to mitigate or remediate viral infection’s effects. At the same time, the actionable targets can inform on the disease’s possible adverse effects and serve as a basis for drug repurposing. This paper presents our efforts to map key molecules highlighted from broad omics data analysis and computational modeling to dysregulated pathways in a cell-or tissue-or patient-specific manner. We then employ text mining and AI-assisted analysis to identify drugs for the retrieved targets. In parallel, we also use topological analysis to reveal interesting structural features of the map.

## 2. Results

### 2.1 Multi-omic data analysis and mechanistic diagram mapping

#### 2.1.1 Footprint-based analysis and causal network contextualization in a SARS-CoV-2 infected A549 cell line

We used a published transcriptome dataset (4) focusing on A549 cells and combined it with phosphoproteomic data of mock-treated and SARS-CoV-2-infected cells (5). We thus contextualized the perturbed signaling events of the viral infection and inferred a causal network using the Carnival tool (6) with the COSMOS approach (7) based on a prior knowledge network assembled from OmniPath resources (8). The Carnival-inferred network connected the top ten deregulated kinases with the top 30 deregulated transcription factors (TFs; **Fig. S1A**). Among the deregulated proteins in the Carnival-inferred network, we found four kinases (TBK1, IKBKE, TICAM1, MAPK3), four TFs (IRF3, ATF4, ATF6, SMAD1), and one serine protease (MBTPS1) among seven diagrams of the C19DMap (**Fig. S1B**; **Table 1**). The results highlighted the activation of the MAP kinase family in response to SARS-CoV-2 infection, a result supported by several publications (18,19). Our approach also highlighted proteins from the curated TGFb signaling pathway, such as MAPK3 and SMAD1, and the signaling proteins PIK3CA, BRCA1, and RUNX1. In addition, TICAM1, TBK1, IKBKE, and IRF3 are found in the results and the curated pathogen-associated molecular patterns (PAMPs) and Interferon-1 pathways. Lastly, we identified relevant players in the Endoplasmic Reticulum (ER) stress pathway, particularly ATF4, ATF6, and MBTPS1. Potential crosstalk between ER stress and immune pathways was discussed in our previous work (1).

**Table 1.**
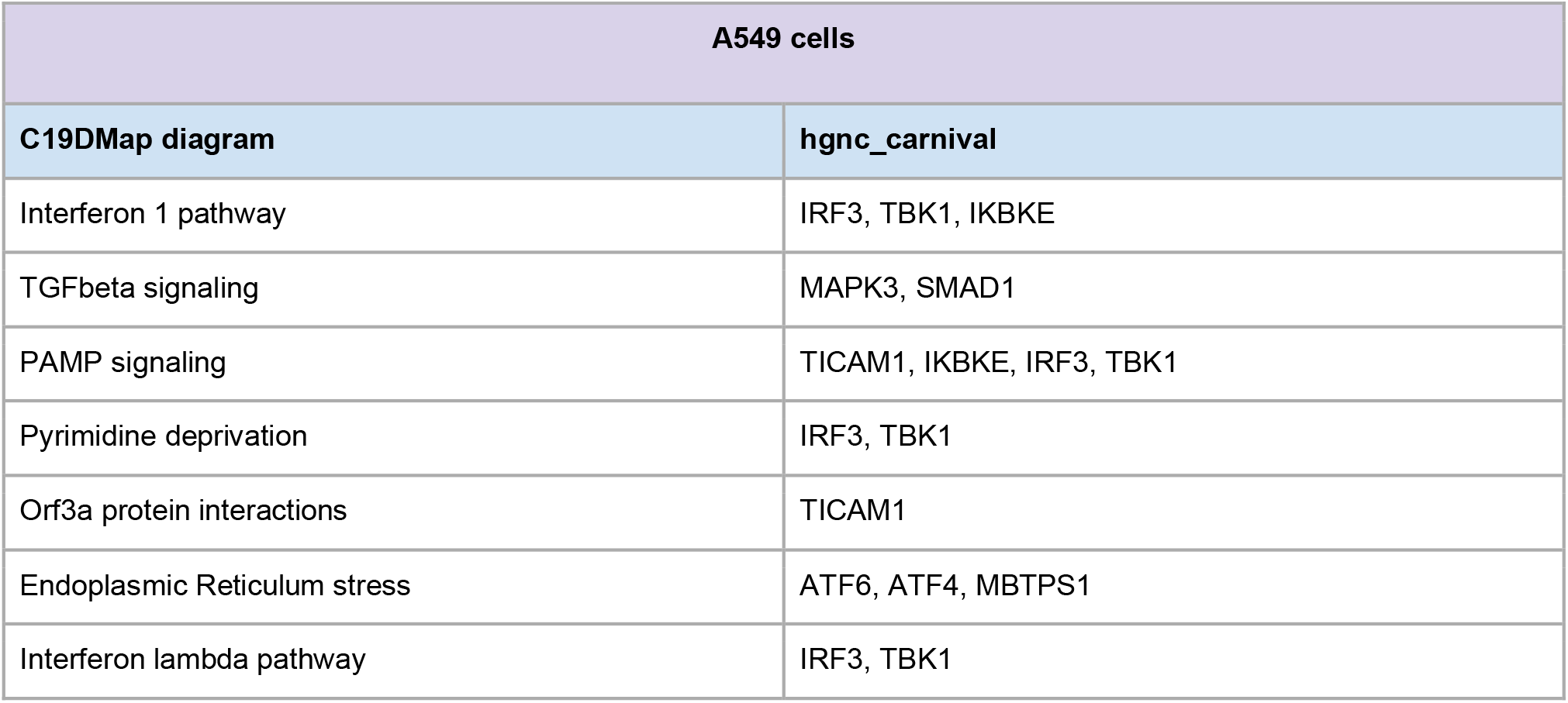
Highlighted C19DMap diagrams, including deregulated kinases and TFs, identified in A549 cells using the footprint analysis.

#### 2.1.2 Transcription factor activity and gene expression analysis in SARS-CoV-2 infected NHBE and A549 cell lines

Next, we expanded our TF study to include RNA-seq data from Normal Human Bronchial Epithelial (NHBE) cells. We used the same datasets (GSE147507) (4, 9) to detect differentially expressed genes (DEGs) between these two cell lines (**Fig. S2A**). TFs that statistically significantly regulate these DEGs were detected by limitless arity multiple testing procedures (LAMP; **Fig. S2B**) (10). Results showed that the number of TFs detected for A549 cells was higher than that for NHBE cells, similar to the results of the DEG analysis. Many TFs detected in both cell types were involved in immune responses. We also performed Gene Ontology (GO) enrichment analysis to determine the functions of the genes regulated by these TFs (**Fig. S2C**). The number of enriched GO terms was greater in A549 cells than in NHBE cells, and almost all NHBE-enriched terms were also enriched in A549 cells. Common terms included those related to the immune system (GO:0002376, immune system process; GO:0002250, adaptive immune response). Several TFs detected in both cell types were also in the Disease Map, while others, such as ESR1 and KLF6, were novel (**Table 2**). These TFs are not yet characterized in the context of COVID-19. Their inclusion in the Disease Map may provide an opportunity to reveal more detailed mechanisms of gene regulation hijacked by the coronavirus infection.

**Table 2:**
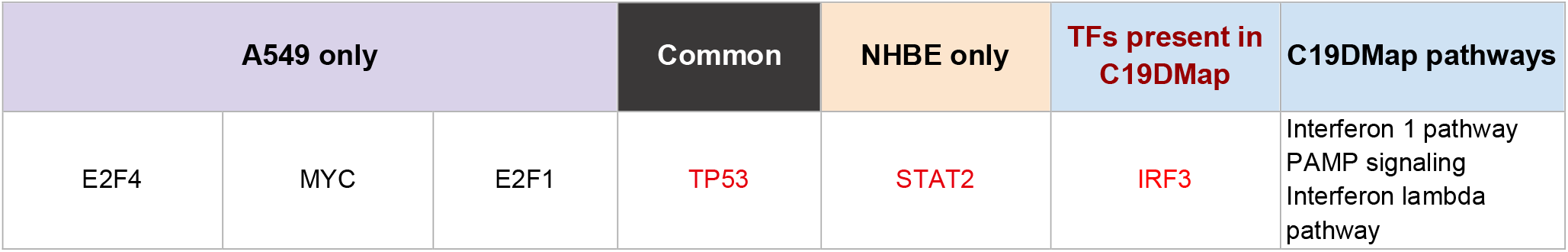
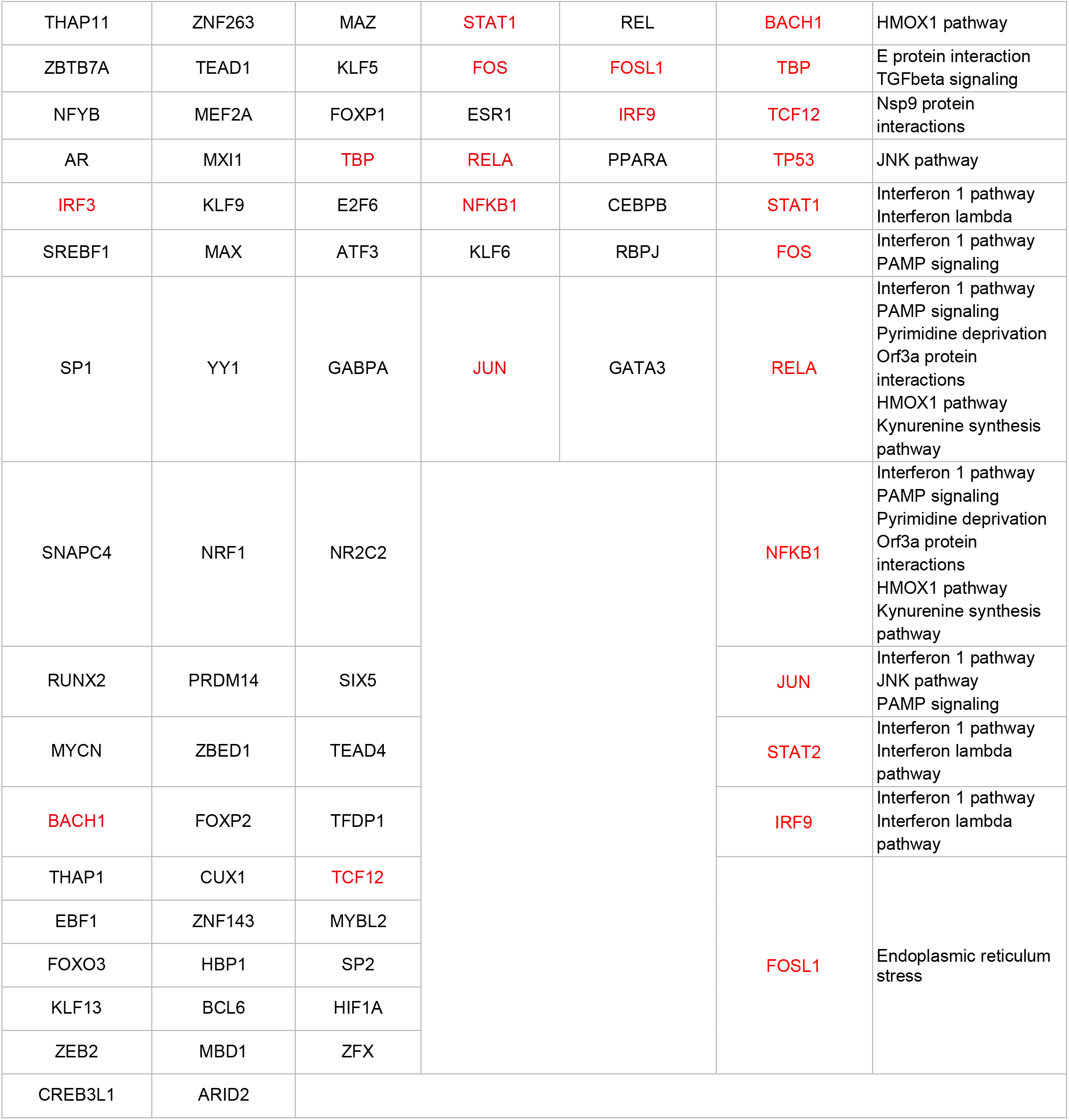
Transcription factors that regulate DEGs before and after coronavirus infection in NHBE and A549 cells; “Common” indicates transcription factors detected in both cell types; transcription factors in red indicate those already present in the C19DMap.

#### 2.1.3 Extended pathway analysis in SARS-CoV-2 infected NHBE and A549 cell lines

Next, we identified altered COVID-19-specific and general molecular pathways in NHBE and A549 infected cells using the same RNA-seq dataset (GSE147507) (4, 9). Then, over-representation analysis was performed on a combined pathway collection from C19DMap (2), WikiPathways (11), and Reactome (12) with 1,840 human pathways containing 12,037 unique genes (**Fig. 2A**). Over-representation analysis revealed 74 altered pathways in NHBE and 101 altered pathways in A549 cells of which 11 pathways were changed in both, including several immune- and metabolism-related pathways (**Fig. 2B**). Interestingly, NHBE cells showed several C19DMap pathways altered after SARS-CoV-2 exposure including interferon and coagulation pathways (**Fig. S3**). However, A549 cells mainly show changes in general processes, of which many have been associated with SARS-CoV-2 infection, including cell cycle, DNA mismatch repair, and cholesterol biosynthesis pathways (**Fig. S4**). A pathway-gene network using the shared DEGs and the C19DMap pathways (23 pathways with 657 unique genes) was created in the following step. In the pathways, 25 genes linked to 19 different pathways were found to be differentially expressed in both cell lines (**Fig. 3A**). Besides the SARS-CoV-2 innate immunity evasion and cell-specific immune response (WP5039; 7 genes) and the interferon lambda pathway (6 genes), also the Type I interferon induction and signaling during SARS-CoV-2 infection (WP4868; 5 genes), the Host-pathogen interaction of human coronaviruses-interferon induction pathway (WP4880; 5 genes) and the Coagulation pathway (4 genes) have several genes that are altered in both cell lines. Central genes in the network are IFIH1 (7 pathways), IL1B (6 pathways), and IRF9 (5 pathways).

**Figure 1.**
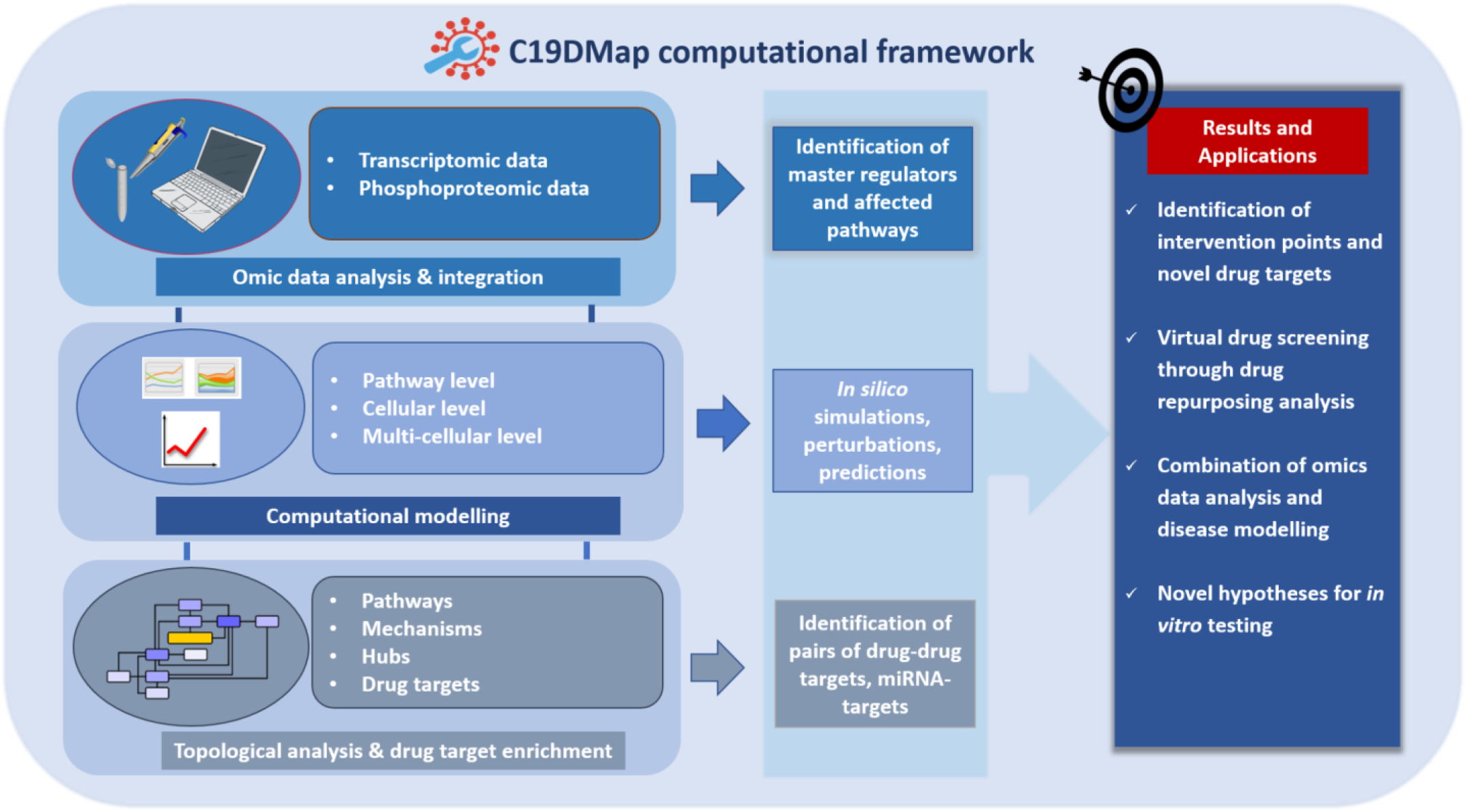
The main workflow of the pipelines developed to analyze the mechanistic content of the C19Dmap. We used it to suggest intervention points, drug repurposing and novel hypotheses for *in vitro* testing.

**Figure 2.**
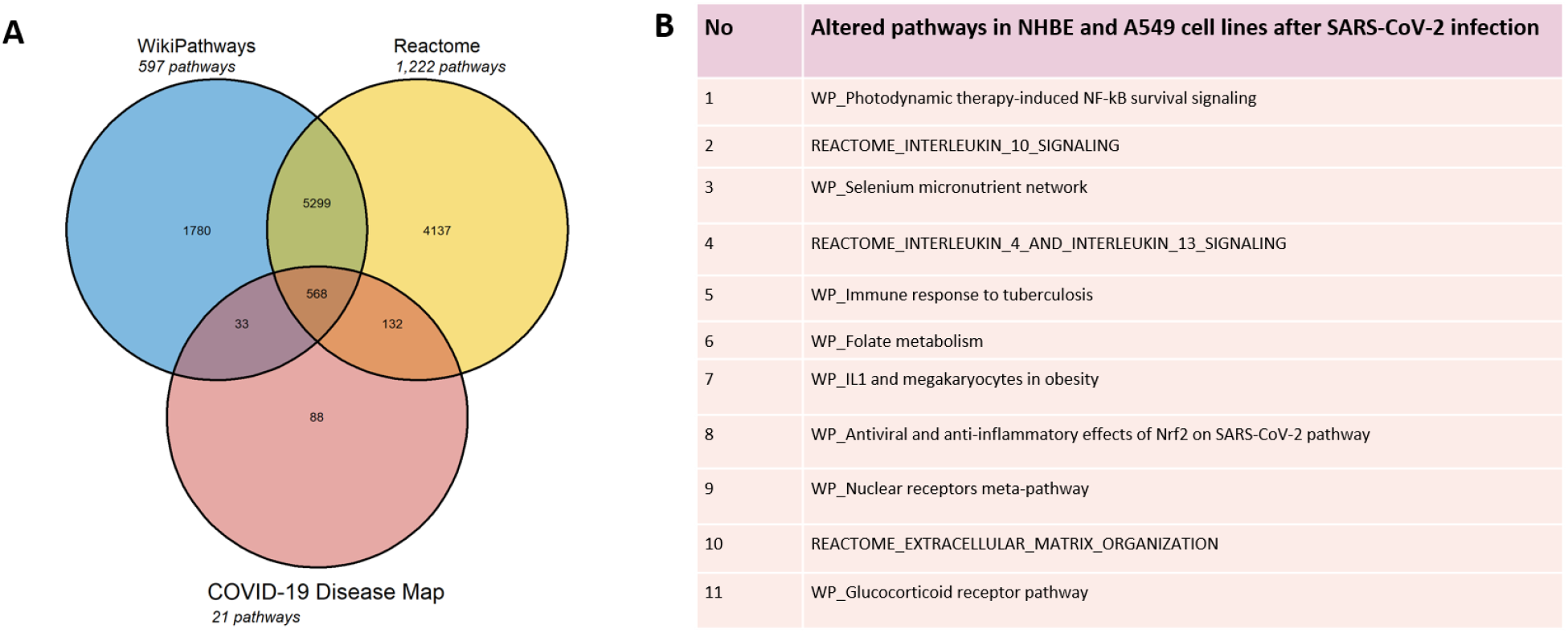
(A) Venn diagram of the combined pathway collection from COVID-19 Disease Map, WikiPathways, and Reactome with 1,840 human pathways containing 12,037 unique genes. (B) Over-representation analysis (criteria: absolute fold change > 1.5 and p-value < 0.05) revealed 11 altered pathways common in both cell lines.

**Figure 3.**
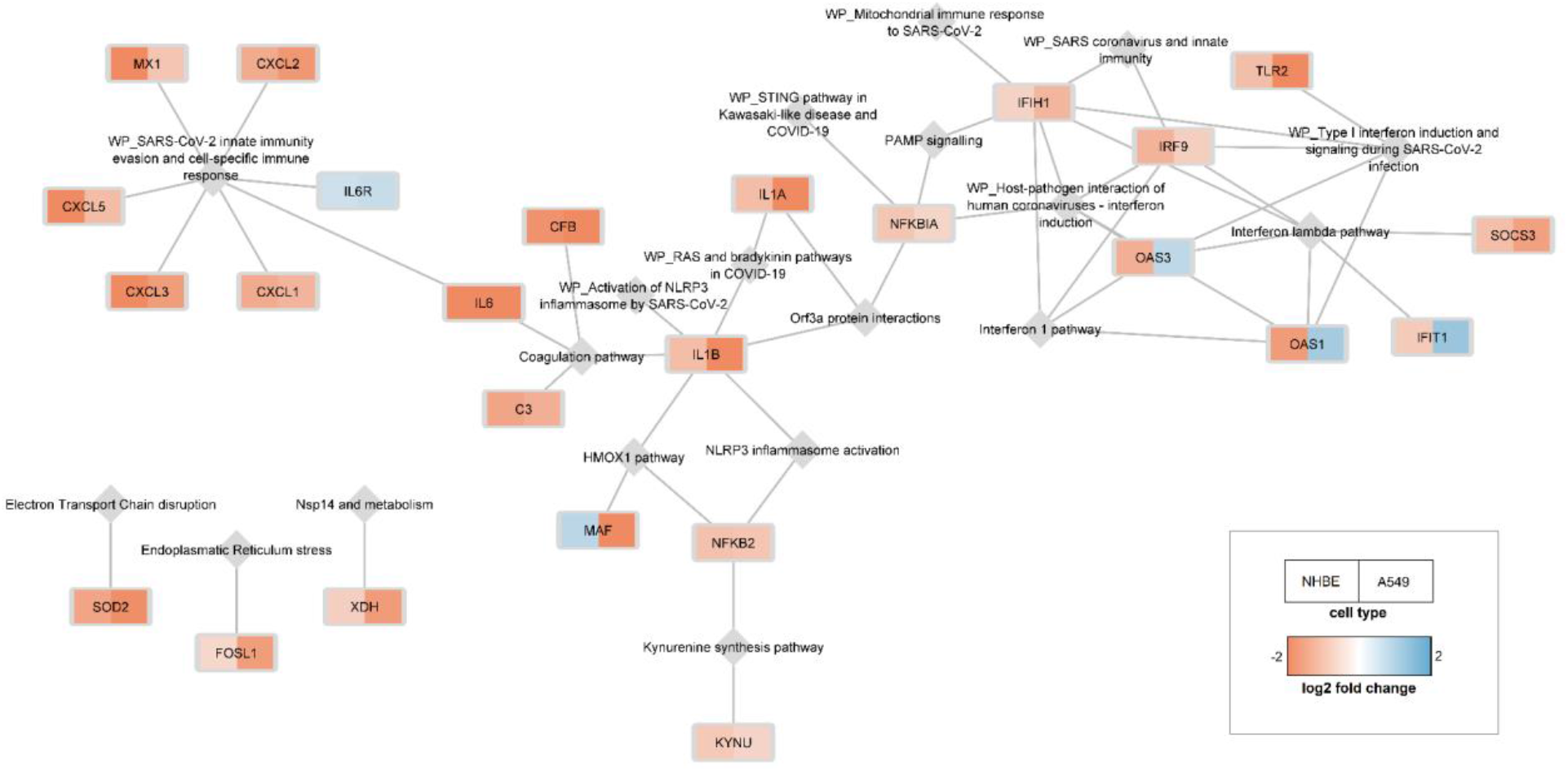
25 genes linked to 19 different pathways, which are differentially expressed in both cell lines.

Interestingly, four genes, namely OAS1, OAS3, IFIT1 from the Interferon pathway, and MAF from the HMOX1 pathway, were found to have opposite expression profiles in the two cell lines. This analysis highlighted that many of the shared differentially expressed genes (134 out of 159) are not yet present in any of the C19DMap pathways, providing an essential resource for future curation efforts to map out and understand the processes affected by SARS-CoV-2 infection.

#### 2.1.4 Single-cell transcriptome analysis in epithelial cell types of COVID-19 patient groups with different severity profiles

Next, we wanted to expand our analyses using patient data. The single-cell RNA sequencing (single-cell RNAseq) dataset was composed of bronchoalveolar lavages from nine COVID-19 patients (GSE145926) (13) and epithelial cells isolated from the lungs of nine healthy subjects (GSE160664) (14). Clustering analysis was carried out on the entire matrix and showed 44 distinct clusters as the best representation of cell types (**Fig. S6**). Five epithelial cell types were selected by cell sample size between groups and gene markers, following the classification of Okuda and collaborators (27). For each cell type, data of moderate, severe, and critical COVID-19 cases were grouped as the category *COVID-19*, and differential expression analysis was performed between *COVID-19* and healthy controls. Among all the DEGs overexpressed in COVID-19 patients in each cell type (**Table S1**), 26 were shared among all lung epithelial cell types (**Table S2**). The overexpressed genes in five cell types of COVID-19 patient groups were reported on the C19DMap to evaluate the activation of specific pathways. In all the epithelia cell types of the COVID-19 group, the genes IFIH1, OAS1, STAT1, OAS2, OAS3, and IRF7, which belong to the type I interferon pathway (WP4868), are found to be overexpressed, meaning that they get activated during SARS-CoV-2 infection. In addition, evidence of direct infections of SARS-CoV-2 in these cell types was confirmed using the databases Reactome (12) and KEGG (15), with activation of pathways linked to Interferon and Influenza A infection, respectively. Interestingly, OAS1, OAS3, and IFIH1 were also found to be differentially expressed in NHBE and A549 infected cells, with OAS1 and OAS3 having an opposite expression profile. STAT1 was also found to be overexpressed in both cell lines. However, IRF7 was not previously identified though members of the same protein family were present in both NHBE and A549 infected cells (IRF3, IRF9). The positive DEGs were reported in the C19DMap as an overlay, viewing only DEGs with a false discovery rate (FDR) ≤ 5 % and |log fold change (logFC)| > 1. Affected pathways were: NLRP3 inflammasome activation, Interferon 1 and Interferon lambda pathways, Virus replication cycle, PAMP signaling, Electron transport chain disruption, E protein interactions, Nsp9 protein interactions, Nsp4/6 protein interactions, Nsp14 protein and metabolism, Orf3a protein interactions, TGFbeta signaling, Orf10 Cul pathway, Endoplasmic reticulum stress, Apoptosis pathway, Kynurenine synthesis pathway, HMOX1 pathway and Renin-Angiotensin pathway (**Fig. 4**). No DEG was mapped onto SARS-CoV-2 RTC and transcription, Pyrimidine deprivation, Autophagy, JNK or coagulation pathways.

**Figure 4.**
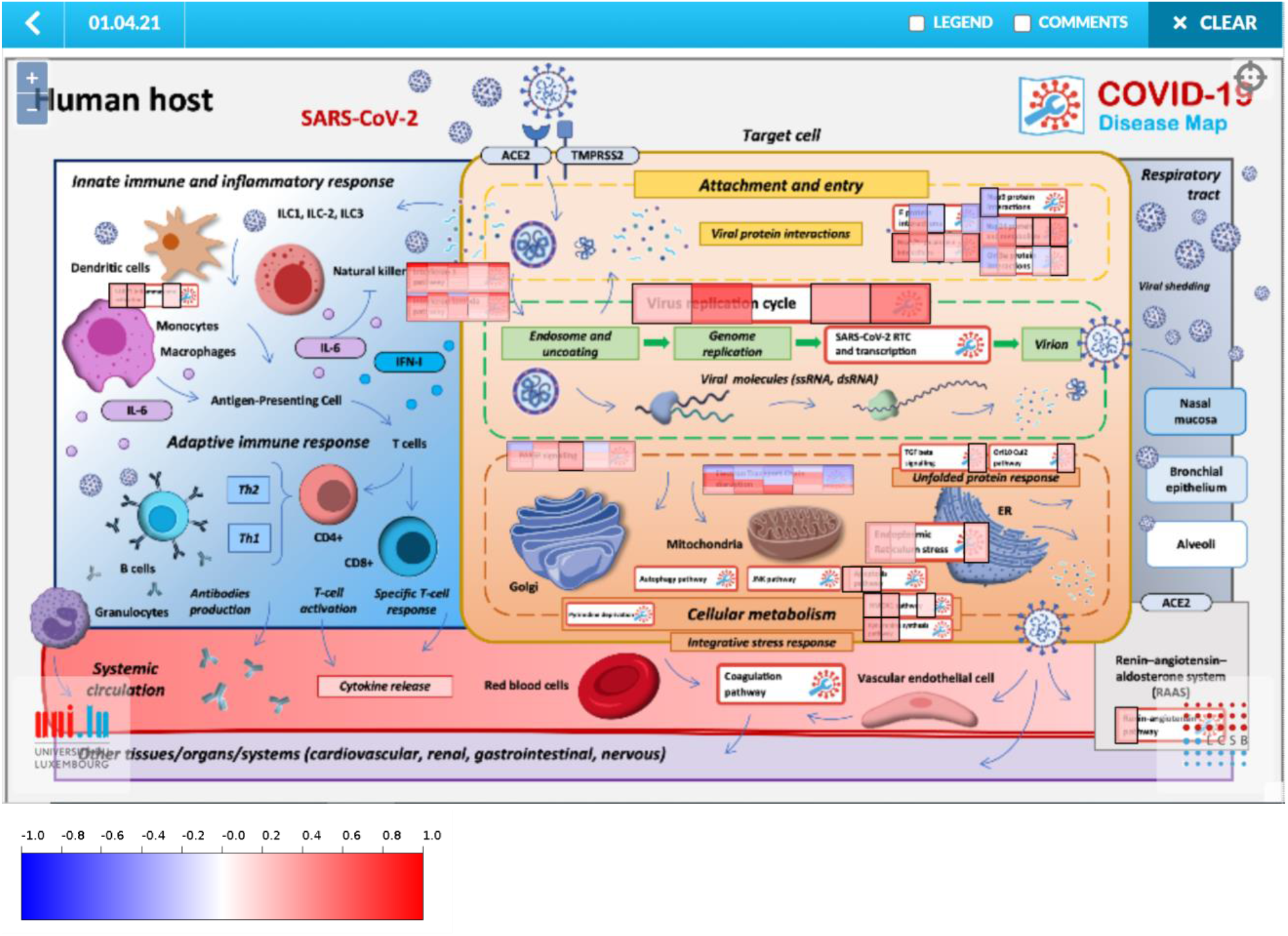
Positive DEGs in the COVID-19 Disease Map repository were reported as an overlay. Only DEGs with FDR ≤ 5 % and |logFC| > 1 are shown.

#### 2.1.5 Combining omics data with mechanistic pathway modeling

To expand on patient data and use the available diagrams in a more active way than mapping, we decided to employ the HiPathia approach (16) that effectively combines RNAseq data with mechanistic diagrams and pathway modeling.

In this step, a public RNAseq dataset of nasopharyngeal swabs from 430 individuals with SARS-CoV-2 and 54 negative controls (17) (GSE152075) was used. Following the pipeline developed for this study, 16 of the 23 pathways were suitable for the HiPathia algorithm. We found that 47 of the 145 circuits analyzed using the HiPathia algorithm were differentially activated (adjusted p-value < 0.05), showing global deregulation of the pathways involved in SARS-CoV-2 infection (**Table S3**). The most representative pathways are shown in Table 3. We then evaluated each pathway containing the deregulated circuits. Almost all of the pathways showed differential activity between infected and normal cells, confirming the relevance of the C19DMap. The apoptosis of the infected cells is a process generally activated in the COVID-19 host response—in fact, the involvement of caspase-3 in SARS-CoV-2-related apoptosis has already been described (18). Moreover, caspase inhibitors have been thoroughly studied because of their therapeutic potential due to the exuberant caspase response in COVID-19 that may facilitate immune-related pathological processes leading to severe outcomes (19).

**Table 3.** Significant pathway activity values after Wilcoxon test comparison between 430 SARS-CoV-2-infected vs 54 non-infected individuals. The results are obtained after running the CoV-Hipathia web tool with the GSE152075 dataset.

| Pathway:<br>Effector Circuit name | UP/DOWN | statistic | FDR | Fold Change | logFC |
| --- | --- | --- | --- | --- | --- |
| Interferon lambda pathway: STAT1, STAT2, STAT3 | UP | 7.16E+00 | 5.77E-11 | 2.01E+00 | 1.01E+00 |
| Interferon 1 pathway: OAS1, OAS2, OAS3 | UP | 6.89E+00 | 2.77E-10 | 2.88E+00 | 1.53E+00 |
| Interferon 1 pathway: ISG15 | UP | 4.26E+00 | 1.81E-04 | 2.55E+00 | 1.35E+00 |
| JNK pathway: JUN, JUND | DOWN | -5.47E+00 | 8.14E-07 | 5.83E-01 | -7.78E-01 |
| Renin-angiotensin pathway: LNPEP | UP | 4.87E+00 | 1.45E-05 | 1.25E+00 | 3.23E-01 |
| Kynurenine synthesis pathway: AHR | UP | 4.28E+00 | 1.81E-04 | 1.18E+00 | 2.35E-01 |
| Coagulation pathway: MAS1 | UP | 3.85E+00 | 5.88E-04 | 1.28E+01 | 3.68E+00 |
| HMOX1 pathway: RBX1, KEAP1, CUL3 | DOWN | -3.87E+00 | 6.33E-04 | 2.73E-01 | -1.87E+00 |
| Orf3a protein interactions: HMOX1, ALG5, ARL6IP6 | DOWN | -3.46E+00 | 7.28E-04 | 2.45E-01 | -2.03E+00 |
| Apoptosis pathway: CASP7 | UP | 3.37E+00 | 3.66E-03 | 1.34E+00 | 4.22E-01 |

As thoroughly described in the scientific literature, impaired coagulation is one of the main complications of severe COVID-19, leading to thrombosis and microthrombosis episodes (20). When examining the C19DMap submap “Renin-angiotensin pathway” (**Fig. 5A**), we found that only one circuit out of the 12 included in the pathway is differentially activated in infected cells. Curiously, this circuit relates to ACE2 and MAS1 as its effector gene is up regulated. The role of ACE2 has been widely associated with SARS-CoV-2 infection (21). Interestingly, it is accompanied by upregulation of the MAS1 circuit related to the normal functioning of the vascular system. The receptor Mas1 induces vasodilation and attenuates vasoconstriction. Moreover, in endothelial cells, activation of the ACE2/Ang-(1-7)/Mas1 axis increases the production of nitric oxide and prostacyclin, both with vasodilator properties and in vascular smooth muscle cells, it inhibits pro-contractile and pro-inflammatory signaling (22). Therefore, the activation of this axis may be a result of a vasoprotective response of the host to the systemic inflammation and vascular injury occurring in COVID-19. On the other hand, the presence of glycoproteins, such as GPVI and vWF, is involved in thromboembolism and thromboinflammation, and other coagulopathies (23). However, recent studies have shown that platelets are indeed hyperactivated in COVID-19 but show reduced glycoprotein VI (GPVI) reactivity in COVID-19 patients (24), which is consistent with our results.

**Figure 5.**
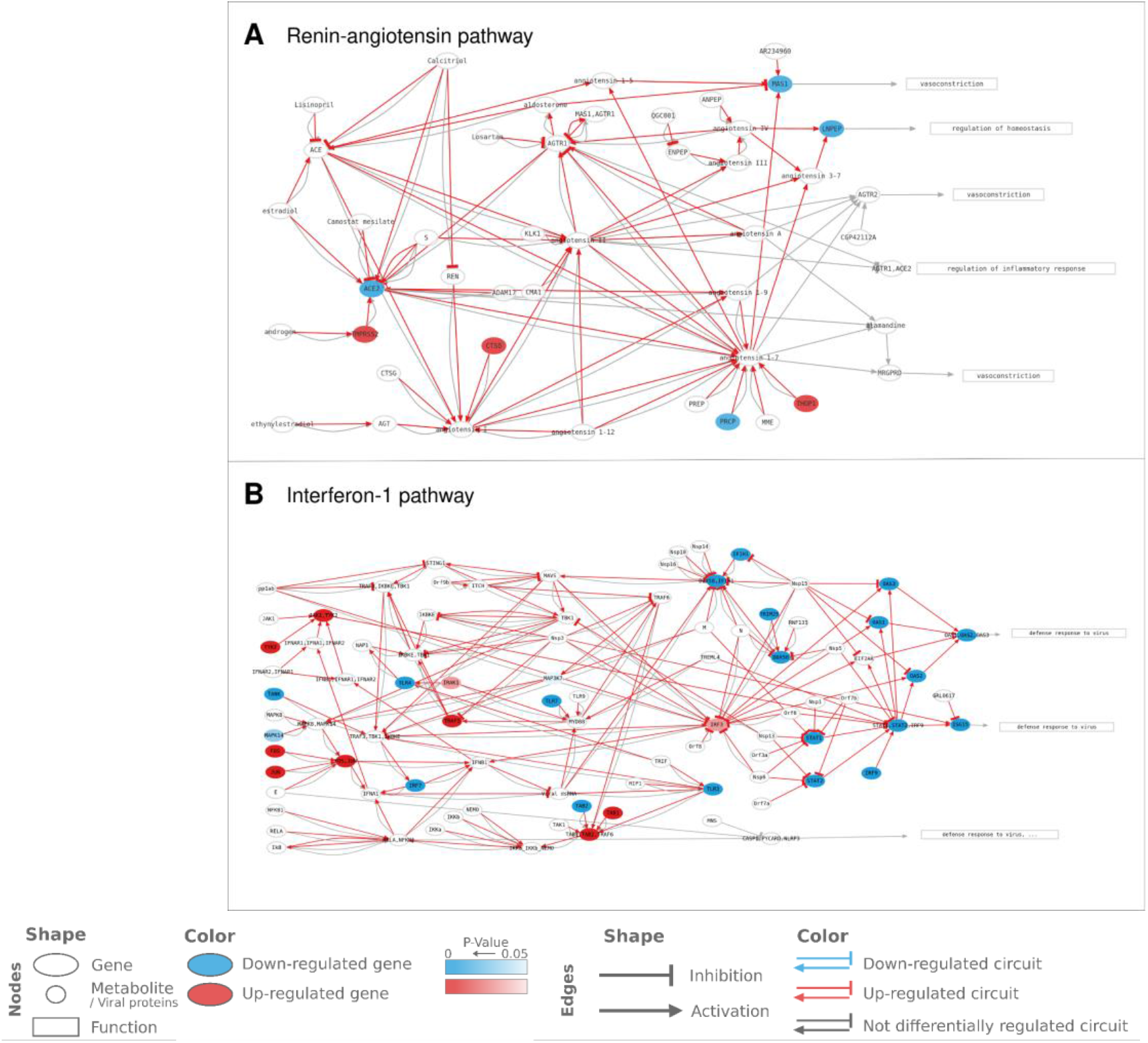
Representation of the activation levels of significant C19DMap pathways in SARS-CoV-2 infected nasopharyngeal tissue. The activation levels have been calculated using transcriptional data from GSE152075 and HiPathia mechanistic pathway analysis algorithm. Each node represents a gene (ellipse), a metabolite/non-gene element (circle), or a function (rectangle). The pathway is composed of circuits from a receptor gene/metabolite to an effector gene/function, which take into account interactions simplified to inhibitions or activations. Circuits activated in infected cells are highlighted by red arrows. The color of each node corresponds to the level of differential expression in SARS-CoV-2 infected cells vs. normal lung cells. Blue: down-regulated elements, red: up-regulated elements, white: elements not differentially expressed. HiPathia calculates the overall circuit activation and can indicate deregulated interactions even if interacting elements are not individually differentially expressed.

The Interferon-1 pathway was highly activated, showing an expected response to virus infection (**Fig. 5B**). However, not all of the genes were overexpressed. The identification of genes which are the most relevant to activate each circuit highlights promising drug target candidates against the downstream processes related to the circuits. Moreover, interferon lambda-1 is a type III interferon involved in innate antiviral responses with activity against respiratory pathogens. In fact, the upregulated circuits show an overall activation of the GO biological process, “defence response to the virus.” Therefore the observed overactivation of the IFN-lambda signaling pathway (**Table 3**) was expected and consistent with studies showing promising results when targeting this pathway as a treatment approach (25).

### 2.2 Dynamical modeling of host-pathogen interactions on a molecular, cellular, and multicellular level

We next studied the impact of upstream regulators on the functional outcome of pathways using dynamical computational modeling. We focused on the Interferon 1 pathway in two different contexts: on a pathway level and on a cellular level integrated into a macrophage model. We then modeled the effects of SARS-CoV-2 on the apoptosis of the epithelium and on the influence of the virus on the recruitment of immune cells by macrophages. In all cases, CaSQ (22) was used to convert the mechanistic diagrams into Boolean models.

#### 2.2.1 A dynamic Boolean model of type I IFN responses in SARS-CoV-2 infection

Type I Interferon (IFN) signaling is an essential pathway of host defence against viral attacks, as highlighted in previous analyses of omics data in both cell lines and patients’ samples. To go one step further in the analysis, we used the type I IFN graphical model available in the C19DMap repository (**Fig. 6**) and the map-to-model translation framework developed in (26) to obtain an executable, dynamic model of type I IFN signaling for *in silico* experimentation. The model obtained included 121 nodes, including three drugs, namely 3,4-methylenedioxy-β-nitrostyrene (MNS) (27), Azithromycin (28), and GRL0617 (29) (**Fig. S7**).

**Figure 6.**
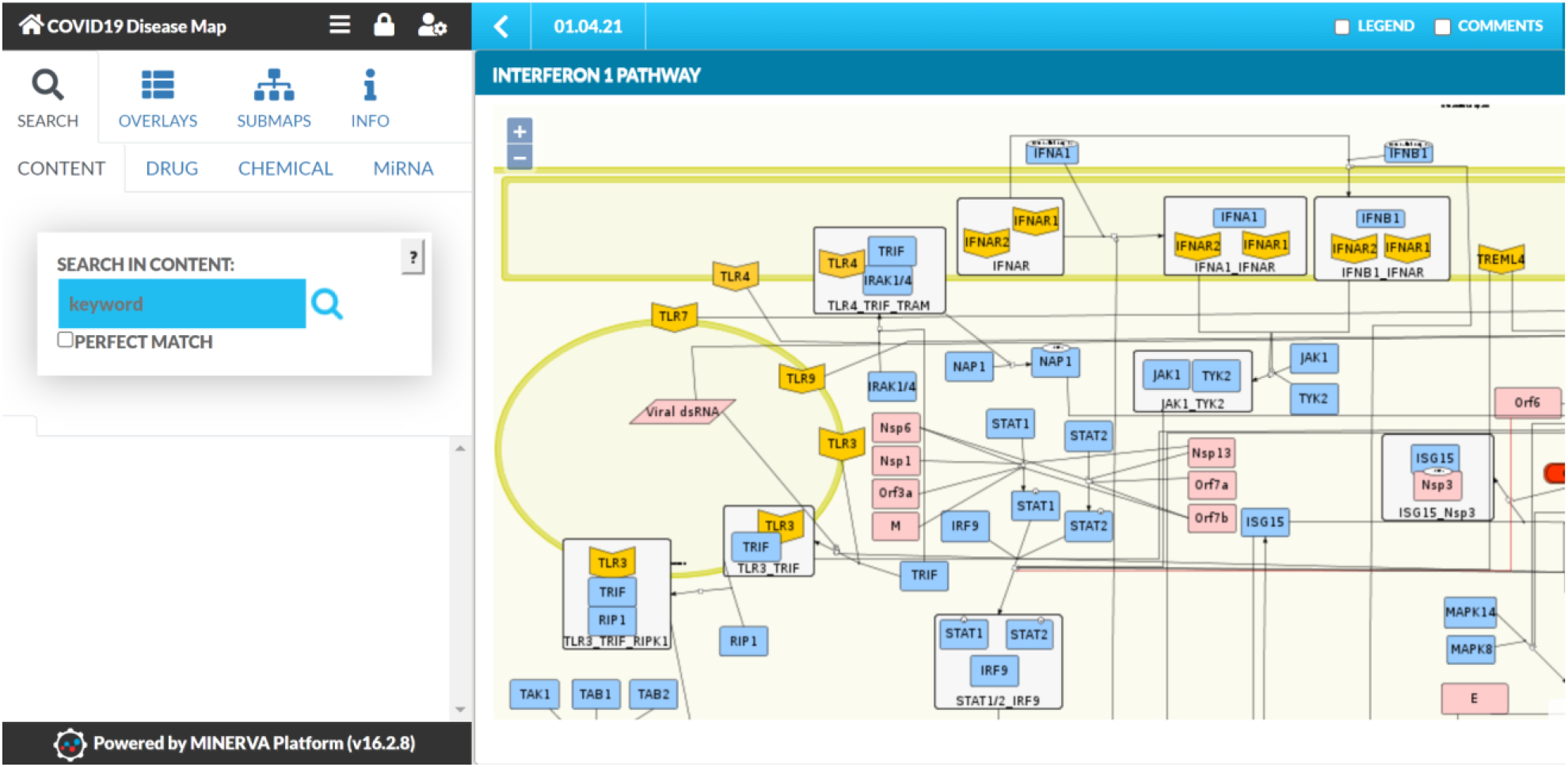
Type I Interferon pathway on the C19DMap repository that was used as a template to produce the Boolean model.

First, to evaluate the model’s ability to reproduce established biological behavior, we performed simulations for seven scenarios derived from the scientific literature (**Table S4**). The model was able to reproduce the behavior for five observations, partially reproduce the behavior for one, and failed to reproduce one biological scenario (**Fig. S8**). Global environmental sensitivity analysis based on partial correlation coefficients using Cell Collective (30) suggested that, in the presence and absence of the drugs, viral E protein had the highest impact on the inflammation phenotype. On the other hand, Nsp3 showed a negative association with the body’s antiviral response. Results of sensitivity analysis with drugs present in the diagram showed that treatment with MNS could reduce inflammation, while Azithromycin was shown to increase the antiviral response (**Fig. S9**).

#### 2.2.2 Sensitivity analysis against knockout and overexpression perturbations

We also performed sensitivity analysis against virtual knockouts (KOs) and knock-ins (KIs), aiming to i) identify the molecules capable of reducing the inflammatory responses and ii) identify the most sensitive viral proteins against knockouts to reduce the viral activity. Results suggested that the overexpression of the IFNB1 RNA had a significant role in the inflammatory process by activating the AP-1 and p50_p65 complexes. IFNB1-induced overexpression is in line with the gene signatures in the Library of Integrated Network-Based Cellular Signatures (https://systemsbiology.columbia.edu/lincs) (31). The IFNB1 RNA increases pro-inflammatory cytokines by activating the NLRP3 inflammasome, while 3,4-methylenedioxy-β-nitrostyrene (MNS) selectively inhibits it (27, 32). However, overexpression of p50_p65 stimulates the inflammatory cytokines via nuclear reactions regardless of the NLRP3 inflammasome inhibition. Therefore, MNS may require a drug combination to reduce the inflammation from nuclear reactions. The viral dsRNA and proteins (Nsp13, Nsp14, and Nsp15) can be significant drug targets since they show potent antagonistic effects on interferon. Literature evidence indicates that such viral molecules have an inhibitory effect on interferons (33). Further, TLR7/9 and TREML4 are the most significant viral binding proteins, suggesting TLR antagonists may be used to control exaggerated inflammations via the MYD88_TRAM complex. Recent proposals consider TLR7/9 as a potential drug target for COVID-19 (34, 35, 36).

#### 2.2.3 Calculating stable states of the IFN model

We used input propagation (37,38) and control nodes to regroup the inputs of the model and simplify the analysis. We regrouped inputs into six categories: 3 meta-inputs that correspond to Inflammatory stimulus, IFN response, and viral stimulus, and three components representing the drugs present in the model (GRL0617, Azithromycin, and MNS). Using this modified model, we could identify 128 stable states and no oscillations. All signatures lack IFN secretion and exhibit either viral replication or antiviral response (or both). To investigate further the behavior of the model, we selected eight configurations for the inputs that cover different biological scenarios of the type I IFN pathway with or without infection and in the presence or absence of drugs (**Table 4**). We then clustered the stable states according to the four outputs of interest, namely viral replication, antiviral response, inflammation, and secretion of IFNA1. For each selected input condition, we have a single attractor (after projection on the outputs; **Table 5**).

**Table 4.** Input configurations that cover different biological scenarios of the type I IFN pathway with or without infection and in the presence or absence of drugs.

|  | C1 | C2 | C3 | C4 | C5 | C6 | C7 | C8 |
| --- | --- | --- | --- | --- | --- | --- | --- | --- |
| viral_components | 1 | 1 | 0 | 1 | 1 | 1 | 1 | 1 |
| immune_response | 0 | 0 | 1 | 1 | 1 | 1 | 1 | 1 |
| IFN_secretion | 1 | 1 | 1 | 1 | 0 | 1 | 1 | 1 |
| Azithromycin_drug | 1 | 0 | 0 | 1 | 0 | 0 | 0 | 0 |
| GRL0617_drug | 1 | 0 | 0 | 0 | 0 | 0 | 1 | 0 |
| MNS_drug | 1 | 0 | 0 | 0 | 0 | 0 | 0 | 1 |

**Table 5.** Projection of the stable states to the four outputs, namely viral replication, antiviral response, inflammation, and secretion of IFNA1. For each selected input condition, we have a single attractor.

|  | C1 | C2 | C3 | C4 | C5 | C6 | C7 | C8 |
| --- | --- | --- | --- | --- | --- | --- | --- | --- |
| ISG_expression_antiviral_response_phenotype | 1 | 0 | 1 | 1 | 0 | 0 | 0 | 0 |
| Viral_replication_phenotype | 1 | 1 | 0 | 1 | 1 | 1 | 1 | 1 |
| Proinflammatory_cytokine_expression_Inflammation | 0 | 1 | 0 | 1 | 1 | 1 | 1 | 0 |
| type_I_IFN_response_phenotype | 0 | 0 | 0 | 0 | 0 | 0 | 0 | 0 |

In these conditions, the propagation of the input values is sufficient to control most components of the model, and in particular, all selected output components. The results of the stable state analyses corroborate the results of experimental studies in patients with COVID-19 with various degrees of severity that showed hampered IFN-I responses in patients with severe or critical COVID-19 (39). These patients had low levels of IFN-I and ISGs, and increased production of tumor necrosis factor (TNF-), IL-6-, and NFkB-mediated inflammation. The result of input propagation can be visualized in a heatmap where lines represent all 121 components of the system and columns represent the eight selected input conditions (**Fig. S10**).

#### 2.2.4 Integration of the Type 1 IFN, the RA system, and the NLRP3 inflammasome curated pathways into a macrophage-specific Boolean model

The population of macrophages expands during SARS-CoV-2 infection, and hyperactivation of these cells can lead to severe immunopathologies (40). To be able to computationally simulate the effects of SARS-CoV-2 on several COVID-related pathways in macrophages, we extended a previously built macrophage polarization model to incorporate biological processes related to SARS-CoV-2 infection, including the Type 1 (T1) IFN response, the Renin-Angiotensin (RA) system, and the NLRP3 inflammasome modules from the C19DMap. The resulting COVID19 Macrophage Model, named MacCOV (https://gitlab.lcsb.uni.lu/computational-modelling-and-simulation/macrophage-model), comprises 131 nodes and 271 interactions manually verified against the macrophage-specific literature. When an inflammatory microenvironment stimulus is simulated, the model reaches a stable state with the respective signaling cascades and inflammatory biomarkers rendered active (inflammatory response; **Fig. 7**). Infection with SARS-CoV-2 stimulates the RA system module, which potentiates inflammation through specific mediators and effectors, like AGTR1/2. Consistent with the literature (41, 42), the virus, through an Orf3a_TRAF3 complex, also triggers the activation of the NLRP3 inflammasome, thus leading to cleavage of proIL-1b and proIL-18 into their functional forms. In addition, although the inflammatory stimuli remain, the stable state analysis indicates that the virus is able to directly activate the expression of proinflammatory markers without the activation of the main signaling cascades. This is because when both inflammatory and viral stimuli are applied together, the model reaches a state similar to virus infection, indicating that the virus ‘overrules’ some of the inflammatory responses that would typically be activated by the inflammatory stimulus alone, namely blocking the activation of TBK1, Pell1, STAT1, and IRF3 (although their expression is increased; see section 2.1), and key effectors in the type 1 IFN cascade (e.g., OAS1-4, expressed upon IRF9 activation). Therefore, the virus itself can trigger the expression of inflammatory biomarkers, whereas, at the same time, it appears to inhibit signal transduction through proinflammatory pathways that crosstalk with the type 1 IFN response. In all cases, the model reaches an inflammatory state, and with viral stimulation, the activation of viral replication and phagocytosis response are also displayed in the macrophage stable state.

**Figure 7.**
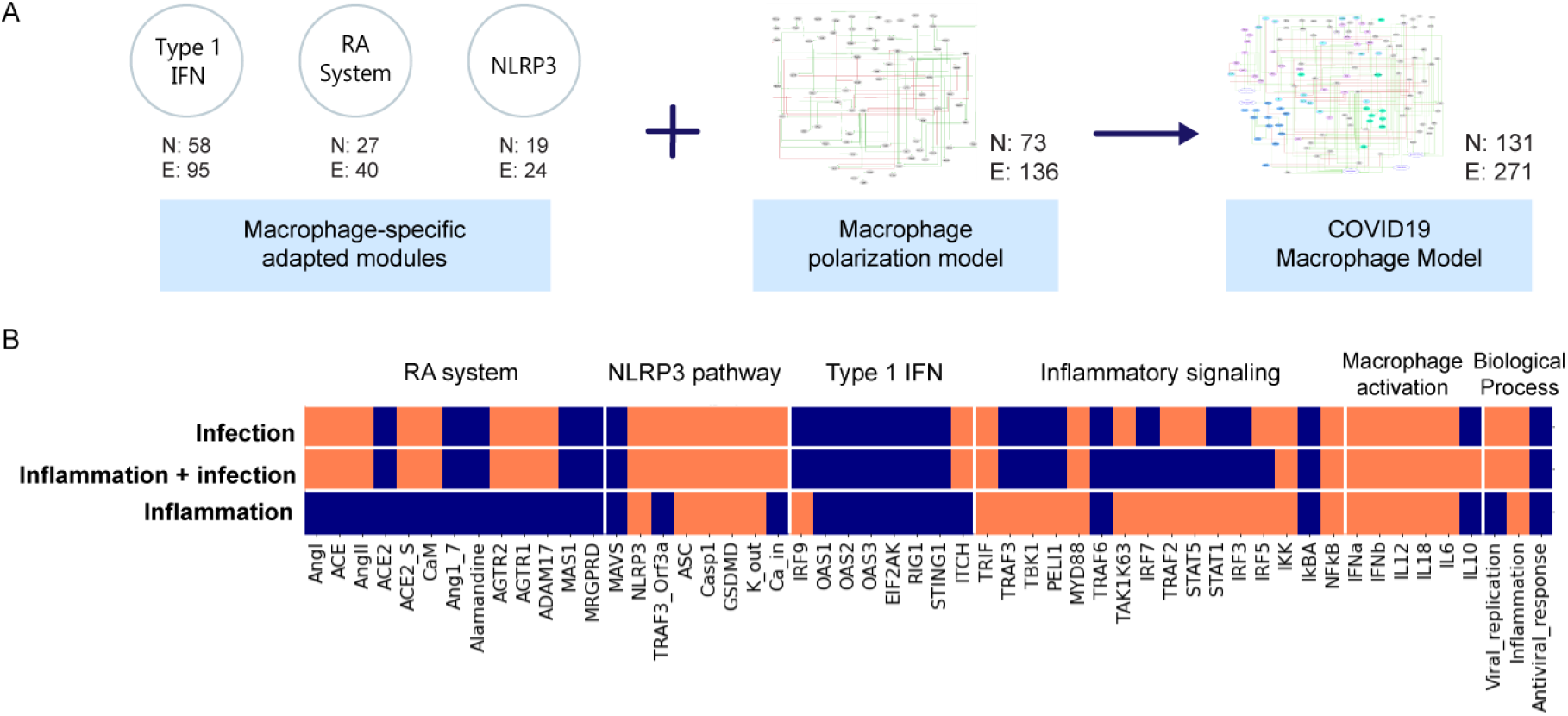
Construction and simulation of a macrophage Boolean model specific for SARS-CoV-2 infection. (A) Modules for the T1 IFN response, the RA system, and the NLRP3 inflammasome were processed with CaSQ to generate Boolean modules, refined, and adapted to be macrophage-specific, and then integrated with a general macrophage polarization model to generate the COVID-19 Macrophage Model (MacCOV). The total number of nodes and interactions in each stage of the processing is indicated in the different panels (N: nodes, E: edges). (B) Model stable states upon different inputs (virus infection, inflammatory conditions + virus infection, and inflammatory condition) are presented in a heatmap. Each input evolves into a unique stable state (rows, delimited by white horizontal lines), where node activity is shown in orange when active and blue when inactive. Nodes, listed at the bottom of the heatmap, are clustered (delimited with white vertical lines) by their relation with specific modules, with the activation of macrophage phenotypes, or with biological processes.

The above results demonstrate that SARS-CoV-2 itself is sufficient to trigger an inflammatory response in macrophages. The virus is also able to block the type I IFN signaling at different levels of the cascade, as demonstrated in the molecular-level model. Lastly, nodes from inflammatory pathways that crosstalk with the type I IFN pathway are also blocked by the virus. By binding to their cognate receptors, proinflammatory mediators activate their downstream signaling effectors, which typically converge in a core pathway (i.e., one that captures signaling from other cascades) or a key proinflammatory transcription factor such as NFkB.

#### 2.2.5 Multiscale and multicellular simulation of SARS-CoV-2 infection uncover points of intervention to evade apoptosis and increase immune cell recruitment

We further expanded our modeling analysis by incorporating two Boolean models into a multiscale simulator of the infection of lung epithelium by SARS-CoV-2 (43) [https://git-r3lab.uni.lu/computational-modelling-and-simulation/pb4covid19]. The two Boolean models focus on the effects of SARS-CoV-2 on the apoptosis of the epithelium and on the influence of the virus on the recruitment of immune cells by macrophages (**Fig. 8**). As previously, CaSQ (26) was used to convert the apoptosis map into a Boolean model.

**Figure 8.**
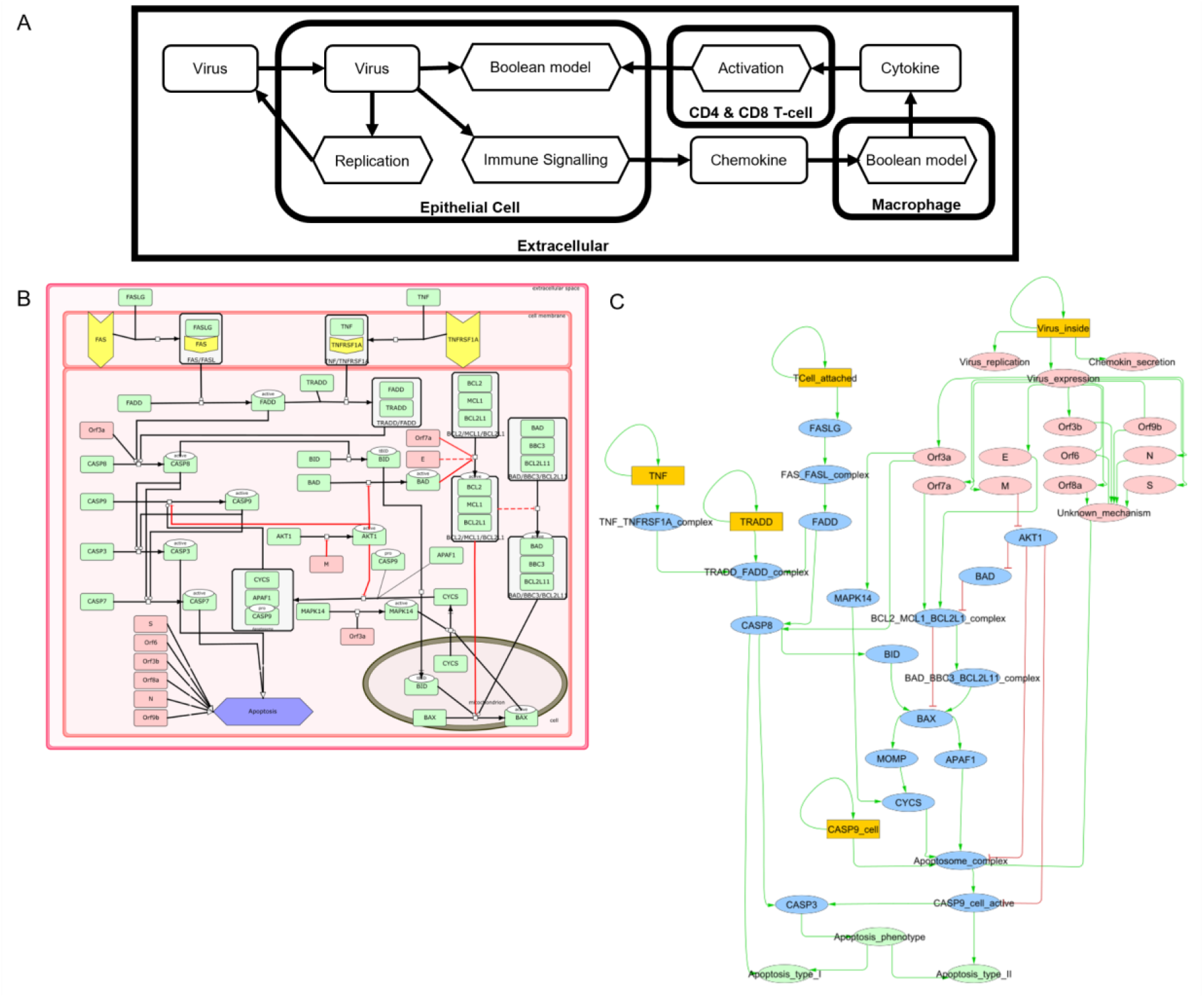
Multiscale simulation workflow. (A) Overview of the top-level interaction model that integrates virus infection, epithelial host cell demise, and the response of different immune cells. (B) The apoptosis model from C19DMap (https://fairdomhub.org/models/712) was used. (C) The modified version of the model was included in each epithelial cell.

We first analyzed the models individually. We studied all the KO of each of the Boolean models (44) to explore and suggest potential drug targets. We identified two perturbations, one that evades apoptosis in infected human host cells and one that increases the immune cell response in macrophages (**Fig. S11**). The first perturbation involved the inhibition of FADD, a downstream actuator of FASLG reception upon T-cell activation promoting apoptosis. In the FADD knockout simulation, CD8-T-cell-mediated apoptosis was abrogated, but the cells were still able to undergo virus-mediated apoptosis through activation of the apoptosome by the virus (**Fig. S12A**). The second perturbation inhibited the macrophages’ p38, a MAP kinase that phosphorylates various proteins in response to stress. We found that the knockout of p38 in this macrophage model increased the recruitment of immune cells by 10% (**Fig. S12B**). In this model, p38 is an activator of pro-inflammatory downstream targets such as AP1, IL1RN, IL1b, IL12, and TNF and is an ERK inhibitor. Thus, p38 knockout having a pro-immune effect is apparently counter-intuitive (45), even though p38 has been described as forming immunity-inhibiting complexes with sestrin during ageing (46). In addition, the pro-immune effect of p38 knockout should also be studied in combination with the SARS-CoV-2 proteins’ triggering of the p38 MAPK signaling pathway to induce apoptosis, as stated above.

We studied the population of epithelial cells and their status (**Fig. 9A**) and the recruitment of immune cells (**Fig. 9B**). Additionally, we incorporated the effect of the mutations in the multiscale simulation: FADD KO behavior in the multiscale model corresponded to the expected behaviors observed in the Boolean model as it reduced the commitment of epithelial cells to apoptosis (**Fig. 9C**). On the other hand, p38 KO in the multiscale model did not substantially change immune cell recruitment by macrophages (**Fig. 9D**). The 10% increase in the recruitment of immune cells seen in the signaling model was not sufficient to see consistent differences when doing replicates of the multiscale simulation.

**Figure 9.**
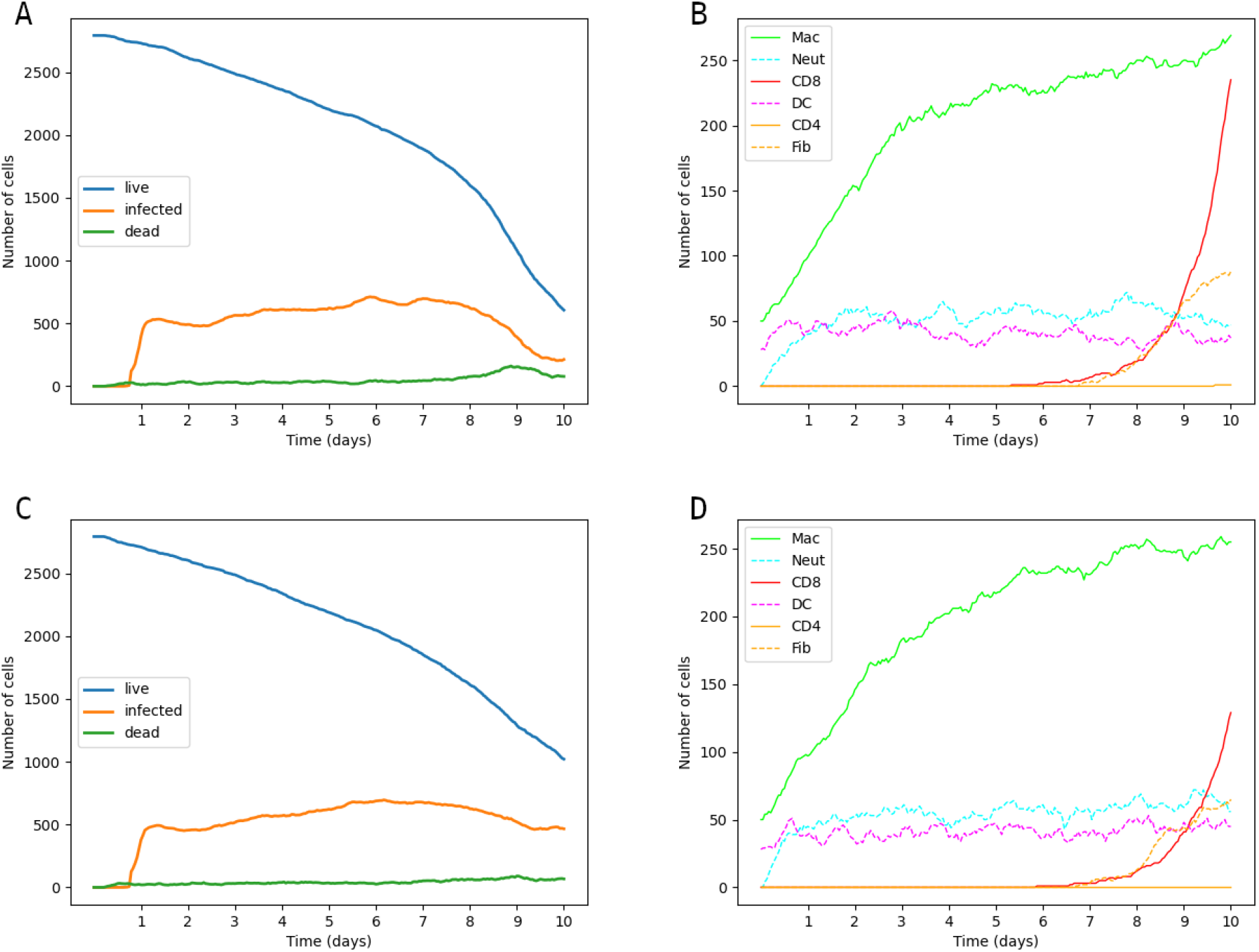
Simulation of wild type and mutants using PhysiBoSS. Our framework can simulate wild-type epithelial cell state (A) and wild-type immune cell recruitment (B) and study the effect of knockouts such as FADD in epithelial cell apoptosis (C) or p38 in immune cell recruitment (D).

### 2.3 Text-mining and AI-assisted drug target enrichment

We used two AI assistants, INDRA and AILANI, to keep the C19DMap up to date and to expand and enrich it with new knowledge. All analyses were performed using a harmonized bipartite graph that included the diagrams from MINERVA, REACTOME, and WikiPathways (See Materials and Methods). In the C19DMap, we now have a collection of 21 MINERVA (5) hosted diagrams, two REACTOME (6) pathways, and 19 WikiPathways (7) diagrams. We created a list of drugs and drug targets using the repository’s content and information from various sources. For example, we provide a list of content-related information from Clinical trials DB, Transcription Factors, drug and protein targets, and miRNA. The corresponding file can be accessed on our public repository. From an initial list of 3,573 proteins extracted from the C19DMap and the drug-target information compiled for the C19DMap, we obtained 1,476 drugs associated with 1,120 drug targets to populate our C19DMap drug target database. We identified 54 targets from the omics data and C19DMap diagrams integrative analysis and the computational modeling analysis (**Table 6**). Using our C19DMap drug target database, we could infer drugs, chemicals, and miRNAs that target these identified nodes (**Table S5**).

**Table 6.** A list of 54 identified targets from the omics data and C19DMap diagrams integrative analysis and the computational modeling analysis.

|  |  |  |
| --- | --- | --- |
| MAPK11 | STAT1 | TREML4 |
| SMAD1 | FOS | TBK1 |
| TICAM1 | JUN | ARL6IP6 |
| TBK1 | RELA | CASP7 |
| IKBKE | NFKB1 | LNPEP |
| IRF3 | STAT2 | HMOX1 |
| ATF4 | IRF9 | FADD |
| ATF6 | BACH1 | AKT1 |
| MBTPS1 | TBP | ALG5 |
| TP53 | TCF12 | AGTR1/2 |
| STAT3 | IFIH1 | EGFR |
| ISG5 | OAS1 | KEAP1 |
| JUND | OAS2 | CUL3 |
| AHR | OAS3 | E |
| FOSL1 | IRF7 | nsp15 |
| MAS1 | BAX | nsp14 |
| RBX1 | IFNB1 RNA | Nsp3 |
| TLR9 | TLR7 | nsp13 |

### 2.4 Pharmacogenomics of drugs targeting the COVID-19 disease map

We collected pharmacogenomic information available in the public domain for the drug targets already present in the C19DMap and assessed the frequency of these genomic variants. We used the “Cumulative Allele Probability” (CAP) and the “Drug Risk Probability” (DRP) scores to summarize the data. The CAP score estimates the likelihood of a particular gene carrying pharmacogenomic variants. In contrast, the DRP score estimates the likelihood of the response to a drug being affected by pharmacogenomic variants (47). The CAP score depends on the number of pharmacogenomic variants and their population frequency. We focused on 79 genes with pharmacogenomic information and allelic frequency information from gnomAD and PharmGKB and calculated CAP scores using gnomAD global exomic information (**Fig. S13**).

The individual CAP scores for the drug target genes were aggregated by drug (**Fig. S14**). Drugs like dabrafenib, migalastat, and erlotinib show low DRP scores across all populations and sexes, whereas others such as methotrexate, capecitabine, and gemcitabine display higher values of DRP scores. Some drugs, such as losartan, show population-related differences in the values of their DRP scores, being higher in Latino/Admixed American and Ashkenazi Jewish than in African/African American populations. Losartan is used to treat hypertension due to its antagonistic effect on the angiotensin II receptor, type 1 (AGTR1) (48). Notably, this protein is involved in one of the circuits from the Renin-angiotensin pathway that are differentially activated in the infected cells compared to controls (see the Mechanistic modeling of COVID-19 disease maps using HiPathia and the macrophage model sections). Currently, 16 clinical trials evaluate losartan’s effect on different outcomes in COVID-19 patients. There are two genomic variants in the *AGTR1* gene annotated to losartan response in PharmGKB (rs5186 and rs12721226). The variant rs5186 is located in the 3’ UTR of the AGTR1 gene. It shows a higher frequency in Ashkenazi Jewish, Latino/Admixed American, and European (non-Finnish) populations (approx. 0.3) than in South and East Asians and Africans/African Americans (<0.1). This variant is associated with increased response to losartan in a study performed on a cohort of European ancestry (49). The other variant, rs12721226, is a missense variant with very low frequency across populations (< 0.01), and the alternative allele (A) is associated with a decreased affinity to losartan and its metabolite EXP3174, which could impair the clinical efficacy of the drug (50). AGTR1 is present in the C19DMap repository and is highlighted as structurally important (ranked 37th in the aggregated graph). Moreover, INDRA analysis retrieved, besides losartan, the drugs telmisartan, irbesartan, valsartan, candesartan, 17alpha-ethynylestradiol, estrogen, nitric oxide, glucose and 1,4-dithiothreitol as able to target AGTR1, while AILANI analysis retrieved besides losartan the drugs candesartan, tasosartan, saprisartan, forasartan, eprosartan, irbesartan, azilsartan medoxomil, olmesartan, telmisartan, valsartan and miRNAs hsa-miR-155-5p, hsa-miR-124-3p, and hsa-miR-26b-5p as molecules targeting AGTR1.

Besides AGTR1, the proteins IKBKE, CASP7, and EGFR are among the identified targets from our analyses for which pharmacogenomics data are available. For IKBKE, the CAP score is very low across all populations, with the lower score achieved for African/African American populations. INDRA analysis retrieved many chemical molecules, and two drugs, amlexanox and sunitinib malate, that have as target IKBKE, while AILANI analysis retrieved the miRNAs hsa-miR-124-3p, hsa-miR-155-5p and hsa-miR-296-5p. Sunitinib shows a high DRP score for East Asian and Latino/Admixed American populations, while it has a very low score for African/African American populations (**Fig. S14**). Amlexanox has no pharmacogenomic data available; however, the drug was used in four clinical trials targeting type 2 diabetes and obesity. Regarding CASP7, the CAP score is very high for East Asians, both male and female, and very low for African/African American populations. INDRA analysis retrieved spermine, 1,4-benzoquinone, melatonin, apigenin, zinc, cisplatin, ac-asp-glu-val-asp-h, nac, fica and emricasan, while AILANI analyses retrieved eight miRNAs that can target CASP7. Among the drugs, pharmacogenomic data were available for cisplatin. Cisplatin has a higher DRP score for Latino/Admixed Americans, both sexes and a lower DRP score across Ashkenazi Jewish and East Asian populations (**Fig. S14**). Emricasan was tested in 18 clinical trials, targeting liver diseases, and recently the drug was tested for its efficacy in COVID-19 disease in 13 patients with mild symptoms; however, no results have been published (https://clinicaltrials.gov/ct2/show/NCT04803227?term=emricasan&draw=4&rank=4). Lastly, for EGFR, the CAP score is very low across all populations, with a slightly higher CAP score for African/African American populations. Using our internal drug-drug target database, we retrieved two drugs, namely zanubrutinib and abivertinib. Zanubrutinib is being tested in clinical trials for the treatment of lymphoma patients (88 clinical trials retrieved from https://clinicaltrials.gov/), while abivertinib has been tested in 11 clinical trials for lymphoma, prostate, and lung cancers and recently was also evaluated in two completed clinical trials for COVID-19 according to https://clinicaltrials.gov/.

### 2.5 Graphical exploration and topological analysis

To cope with the size and complexity of the ever-growing content of the mechanistic pathways, we developed and implemented a concept for the hierarchical exploration of the C19DMap and performed a comprehensive analysis of node centralities on two levels: on the level of the individual pathways for all three platforms and on the level of an aggregated network combining all individual pathways. The implementation is based on the biological network analysis tools Vanted (51), SBGN-ED (52) and a customized version of LMME (Large Metabolic Model Explorer) (53), LMME-DM (**Fig. 10**).

**Figure 10.**
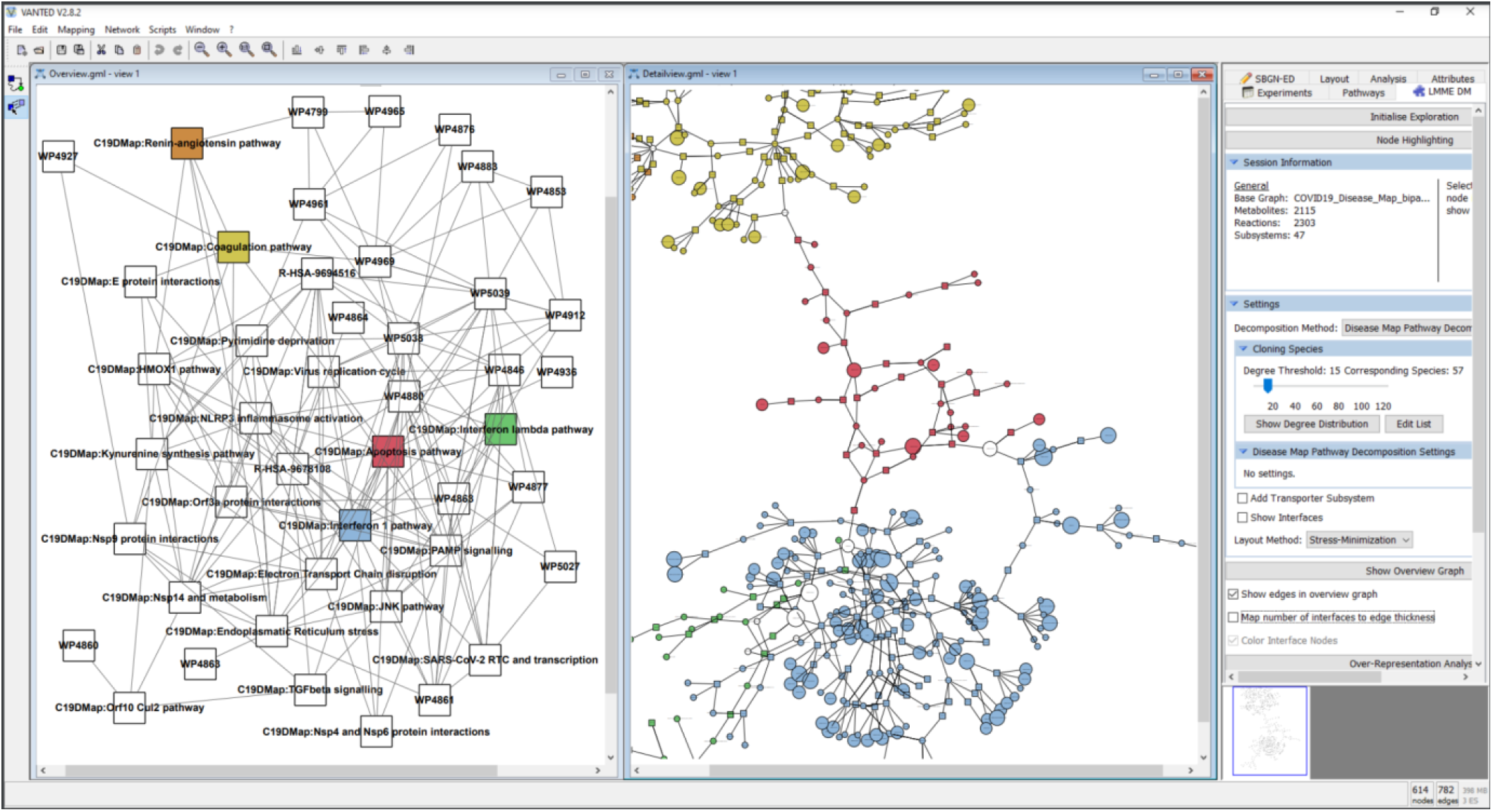
Hierarchical exploration of centrality values in the disease map using LMME-DM. The following pathways are shown in detail: Coagulation: yellow; Apoptosis: red; Interferon 1: blue; Interferon lambda: green; Renin-Angiotensin: orange. The aggregated centrality values are mapped to the node sizes in the detail view.

On all networks combined in the bipartite graph (individual pathways and aggregated network), we performed centrality analysis and computed an aggregated centrality value (see Materials and Methods) to identify the top-ranked species of the C19DMap bipartite graph (**Table S6**). Not surprisingly, the proteins that show up in the top ten are viral proteins and the ACE2 protein that acts as a receptor for the SARS-CoV-2 spike protein. Topological analyses can highlight targets and hubs, providing a basis for linking pathway structure with key findings from text mining, omic data analysis, and modeling pipelines. For the five representative C19DMap pathways, namely Interferon type I, Interferon lambda, coagulation, apoptosis, and renin-angiotensin, we used the aggregated ranks to create a high-level view of the pathways visualizing their connections and also creating nested nodes for coping with complexity (**Fig. 10**). Moreover, from the 54 highlighted targets (**Table 6**), nine of them are characterized as structurally important in the respective pathways, namely TBK1, IKBKE, IRF3, MAS1, IRFNB, CASP7, FADD, AKT1 and AGTR1/2, as they appear in the top ten occurrences of each of the five pathways shown in Tables S7-S11. While the topological features for the aggregated pathway (where all content is unified across the three platforms, MINERVA, WikiPathways, and Reactome) were not so easy to calculate due to incompatibilities that we will need to address in the future versions of the repository (for example, different naming for the same complex, such as AP-1 or AP1, different spellings of nodes using or not capitalized initials, such as nsp13 or Nsp13), we were able to have clean topological features for 26 of the 54 targets. Among these, 18 targets appeared in the top 1000 occurrences of the aggregated pathway (**Table S12)**, with 11 targets characterized as structurally important as they showed up in the top 30% **(Table S13)**.

### 2.6 FAIRness and availability for proper data management

We made considerable efforts to align our work with the four FAIR principles: Findability, Accessibility, Interoperability, and Reusability (54). As this is an ongoing effort, we try to balance our results between timely availability and FAIRness in progress. The tools implemented in our ecosystem are published and indexed on PubMed and searchable online. We try to advance, communicate and exchange with other Systems Biology communities, especially when it comes to the annotation and curation of models (55, 56). All tools are open access, and WikiPathways (11), REACTOME (12), MINERVA (57), AILANI (https://www.biomax.com/products/ailani-for-semantic-integration-and-search/) and CellCollective (30) provide APIs. The developed maps and models are available on GitLab (https://git-r3lab.uni.lu/covid/models/) and FAIRDOMHub (58). We have worked on tool interoperability and promoting community standards; therefore, most input formats are GML, SIF or SBML, and SBML Qual files in an effort to enhance model reusability (59). All maps and models are available under a CC-BY license. Appropriate metadata associated with each of the analyses and modeling results presented in the article is registered and indexed on FairdomHub to facilitate accessibility. Furthermore, we plan to submit the models obtained to model repositories such as the Cell Collective (30), GINsim (60), and BioModels (61). We have also built the C19DMap-Neo4j graph database by integrating the content of the C19DMap diagrams available in MINERVA into the Neo4j framework. This database is available for online exploration at https://c19dm-neo4j.lcsb.uni.lu and is used as a backend solution for efficient access to the resource data. Biological concepts from the C19DMap diagrams available in MINERVA (such as macromolecules and processes) are stored in the database under Neo4j nodes. In contrast, relationships between these concepts (such as consumption and catalysis) are stored as Neo4j relationships. In addition, annotations, such as UniProt identifiers and PubMed publication IDs, are stored in the form of individual nodes that we can easily query (for an example, see **Fig. S15**).

## 3. Discussion

We have explored the high-quality, manually curated mechanistic content of host-pathogen interactions using a number of computational frameworks and bioinformatics analyses that are now combined in interoperable pipelines. To further prioritize targets and contextualize the mechanistic content with different layers of biological data, a set of different omics data was used, ranging from infected cell lines to bulk RNAseq and single-cell omic data from patients affected with SARS-CoV-2. In summary, we used omics data following SARS-CoV-2 infection to infer a causal network describing signaling events perturbed after viral infection. We identified the MAPK protein family as a key mediator of the referred signaling events. Our omics-based approach was able to capture several genes present in the pathways manually curated by the C19DMap community. Furthermore, we found additional causal interactions suggesting the potential mechanism behind the crosstalk between some of the most relevant pathways upon SARS-CoV-2 infection, such as EGFR, PI3K, and the PAMPs/interferon-1 pathway. Focusing further on transcription factors, the analysis revealed new transcription factors not yet included in the C19DMap. Their inclusion may provide an opportunity to reveal more detailed mechanisms of gene regulation hijacked by coronavirus infection. The results showed that, among the drugs targeting transcription factors detected in both cells, 47 were already in external clinical trials, including drugs evaluated for their effectiveness against COVID-19. In addition, we also retrieved 160 drugs that have not yet been tested in clinical trials or tested for efficacy against COVID-19 and could represent potential candidates for further evaluation (**Table S14**). Lastly, over-representation analysis revealed 58 affected pathways in NHBE cells and 39 enriched pathways in A549 cells, including pathways relevant to immune response, the NFkB pathway, glucocorticoid receptor and MAPK signaling pathway, and pathways related to interferon.

The single-cell RNAseq data analysis of a small group of patients confirmed some of the previously identified TFs, DEGs and altered pathways pointed out by the cell line analysis. However, the number of patients in this analysis was relatively small. To expand our analysis, we used an extensive dataset of 450 patients and the HiPathia modeling algorithm to identify affected circuits in the mechanisms described in the repository. We found pathways, such as apoptosis, to be systematically up or downregulated, which means that the whole pathway is relevant to the progression of the disease. Moreover, more extensive pathways showed differential activation in a few or even one of the circuits, which may indicate that, despite the involvement of the whole pathway in the disease progression, only a few processes reflected in the deregulated circuits are critical to the mechanism of infection. These specific key processes may support finding new therapeutic targets. The extensive integrative omic data analysis using RNAseq bulk and single-cell data and the pathway resources revealed interesting TFs,DEGs, and altered pathways after the SARS-CoV-2 infection in the two studied cell lines and in patients’ data. The methodologies used for this step were complementary, covering a wide range of state-of-the-art pipelines and bringing forward two significant points: the coverage and relevance of the C19DMap repository regarding the COVID-19 disease and the identification of additional regulators that would need to be included in the resource.

The COVID-19 disease maps can also be analyzed using computational modeling approaches. Indeed, these disease models can help elucidate mechanisms deregulated at molecular, cellular, and multicellular levels to gain insight into COVID-19 underlying processes. Type I Interferon (IFN) signaling is an essential pathway of host defence against viral attacks, as highlighted in previous analyses of omics data in both cell lines and patients’ samples. We used the executable, dynamic model of type I IFN signaling of our repository for in silico experimentation. The results of the computational modeling showed a complete lack of IFN signatures under relevant conditions matching the experimental results that showed hampered IFN-I responses in patients with severe or critical COVID-19 (36). These patients had low levels of IFN-I and ISGs, and increased production of TNF-, IL-6-, and NF-κB-mediated inflammation. Adding the IFN response, Renin-Angiotensin mechanism, and NLRP3 pathways from the C19DMap to an existing macrophage polarization model helped elucidate the innate immune response that macrophages trigger upon acute COVID-19, in addition to highlighting their contribution to the disease’s pathology. Lastly, the integration of both pathway and cell models in a multicellular-multiscale model helped to reveal the impact of mutations of FADD and p38 on the cellular death of epithelial cells upon infection, as well as on the recruitment of immune cells.

In an effort to further enrich the content, AI-assisted text mining systems, such as INDRA and AILANI, were employed to infer from the vast literature the drugs, miRNAs and chemical molecules that have as targets the biomolecules included in the diagrams of the C19DMap. Text mining and AI solutions can help enrich the content and provide further directions to fill in knowledge gaps. Furthermore, integrating publicly available data from the C19DMap, PharmGKB, and gnomAD allowed us to determine the presence of variants with pharmacogenomic impact and their frequency in human populations. We thus estimated the genomic variability of genes from the C19DMap that was involved in drug response across different populations and sexes. We were able to retrieve pharmacogenomic information for about 79 genes present in the repository, four of which were also identified as potential targets. Topological analyses revealed interesting information about hubs and shared molecules among pathways that could help us better understand the potential upstream and downstream effects of targeting them.

### Perspectives

As mentioned in our previous report (2), most of the diagrams of the CD19DMap repository were initially built using the scientific literature on SARS-CoV-1 and other coronaviruses that were available during the onset of the pandemic. This corpus provided the foundation for rapid curation and a literature triage approach. Annotations for the SARS-CoV-1 viral infection process, including the viral life cycle, host interactions, and therapeutic pathways, were built on this foundation. After more than two and a half years since the appearance of the SARS-CoV-2 virus, the body of scientific literature specific to this type of coronavirus has reached a point where it can now be used to curate complete mechanisms. With the continuous update of pathway information and new datasets related to SARS-CoV-2, reproducible and automated data analysis workflows can be rerun to provide more accuracy and specificity. Generation of Reactome’s SARS-CoV-2 pathway leveraged the database’s foundational manual curation, orthoinference projection, and the collaborative resources of the CD19DMap project. The SARS-CoV-2 infection pathway emerged from a computationally generated rough draft via the orthoinference process from the manually curated, peer-reviewed Reactome SARS-CoV-1 infection pathway (see Materials and Methods). The community can adopt this approach that identifies SARS-CoV-2-specific interactions to increase viral specificity in the mechanisms included in the C19DMap repository.

We made considerable efforts to increase interoperability and communication across three different platforms, MINERVA, WikiPathways and Reactome, support Systems Biology standards such as SBGN (62) and SBML (63), and promote scientific openness with the use of public repositories and the adoption of FAIR (Findability, Accessibility, Interoperability, and Reusability) Data principles (54).

We have successfully built seamless workflows that allow us to use high-quality, curated mechanistic content for integrative analysis and computational modeling. The interoperable pipelines developed and demonstrated here are highly adaptable to new challenges due to standardized formats, can support the testing of combinatorial therapies, as multiple drugs and targets are suggested, and offer a canvas for evaluating the repurposing of existing drugs to fight new waves of COVID-19 or other pandemics, and contribute to elucidating the etiologies of post-acute Covid Symptoms (PASC). By comparing the mechanisms and drug targets, we can further look into the comorbidities of the disease. The C19DMap computational framework is flexible, expandable, accessible, and available freely to the scientific community.

## 4. Materials and Methods

### 4.1 Using the mechanistic diagrams for omics data analysis

#### Footprint analysis

We obtained the transcriptomics dataset from the GEO database with accession number GSE147507 (4). We extracted series number 5 from the dataset, consisting of 2 conditions, A549 cells either mock-treated or infected with SARS-CoV-2, measured in triplicate 24 hours after infection. Differential analysis of the transcript abundances was performed using DESeq2 (64). The resulting t-values of the differential analysis were used as input to estimate pathway activity deregulation using Progeny (65). The differential analysis t-values were also used to estimate the deregulation of TF activities using Dorothea (66) as a source of TF-target regulon and the Viper algorithm (67) to estimate the TF activity score. Phosphoproteomic data of mock-treated and SARS-CoV-2 infected cells were extracted from (5). Phosphosite differential analysis log2FC was used to estimate the deregulation of kinase activities using https://github.com/indralab/protmapper as a source of kinase-substrate interactions and a z-test to estimate kinase activity score (68, 69). Finally, we used Carnival (6) with the COSMOS approach (7) to connect the top 10 deregulated kinases with the top 30 deregulated TFs with a Prior Knowledge Network assembled from OmniPath resources (8). Progeny pathway activity scores were used to weigh the PKN and facilitate the optimal network search to connect kinases and TFs. To place our results in the context of the whole study, we matched the genes obtained in carnival results with those included in the curated pathways by the Covid-19 Disease map community (https://covid.pages.uni.lu/map_contents). In addition, we matched our results with a harmonized list containing drug-targets.

#### TF activity and drug target identification

In this analysis, we inferred the gene regulatory systems that are hijacked by COVID-19, especially the target transcription factors. In order to infer the target transcription factors, we detected transcription factors that statistically significantly regulate the genes whose expression changes were induced by COVID-19. First, the gene groups whose expression changes were induced by COVID-19 in NHBE cells and A549 cells were detected as the DEGs using DESeq2 (64) for the GSE147507 dataset (4, 9) described above. Next, we extracted all the regulatory relationships with Confidence “A”, “B”, and “C” from DoRothEA (66) as information on the regulatory relationships of transcription factors to each of these DEGs for NHBE cells and A549 cells. The transcription factors that regulated each of these DEGs for NHBE cells and A549 cells were detected by LAMP (10) (significance level < 0.05). Next, to gain insight into the biological phenomena affected by the detected transcription factors, i.e. the transcription factors hijacked by COVID-19, gene ontology enrichment analysis of DEGs under the control of these transcription factors was performed using the GOstats package (70) in R (significance level α = 0.05). In order to verify whether these transcription factors are included in the publicly available C19DMap (2), we performed a search based on the HGNC ID of each transcription factor against the SBML file of each Disease Map. Finally, we searched for and picked up the drugs that target each of the transcription factors for NHBE cells and A549 cells that have been in the clinical trials in anticipation of later usefulness for the treatment of COVID-19 as follows. To find the drugs which target the above transcription factors, we conducted a search against GeneCards (https://www.genecards.org/) (71) based on the HGNC IDs of the transcription factors. After that, we performed another search based on those drugs against the list of the drugs in External Clinical Trials for COVID-19 and Related Conditions in the COVID-19 Dashboard of DRUGBANK (https://go.drugbank.com/covid-19) (72). Only approved drugs were listed as candidate drugs in the final results. Finally, to identify gene regulatory systems affected by COVID-19 independent of cell type, DEGs, transcription factors, enriched GO terms, and drug targets detected against NHBE, A549 cells were classified for one or both cell types.

#### Pathway and network analysis in SARS-CoV-2 infected NHBE and A549 cells

We demonstrate an automated and reproducible workflow for transcriptomics data analysis using pathway- and network-based approaches (see our GitLab repository for details;https://gitlab.lcsb.uni.lu/computational-modelling-and-simulation/pathway-analysis-and-extension). The analyses are fully automated in R with clusterProfiler (73) and RCy3 (74) to connect to the widely adopted network analysis software Cytoscape (75) for network visualization. We obtained the transcriptomics dataset from the GEO database with accession number GSE147507 (4). We extracted series numbers 1 (NHBE) and 5 (A549) from the dataset, consisting of 4 conditions in triplicate, NHBE and A549 cells treated with mock (two controls) and NHBE and A549 infected with SARS-CoV-2, measured 24 hours after infection. Pre-processing and differential gene expression analysis was performed in R using the DESeq2 package (64). Next, a combined pathway collection of the COVID-19 Disease Map (21 pathways (76)), WikiPathways (597 pathways (11)) and Reactome (1,222 pathways (12)) was created. Pathway enrichment analysis was performed using the clusterProfiler R package (73). Differentially expressed genes (DEGs; p-value < 0.05 and absolute fold change > 1.5) were used as input for the over-representation analysis. The analysis was performed separately for NHBE, and A549 cells and the overlap in enriched pathways was analyzed. Selected pathways are visualized in Cytoscape using the WikiPathways app (77). A pathway-gene network for the shared pathways was created to study pathway crosstalk and overlap. Next, the harmonized bipartite graph was used to create a pathway-gene network for all C19DMap pathways. By overlaying information about differential expression and filtering for shared differentially expressed genes, we used the network to identify relevant biological processes as well as molecular mechanisms that may be missing in our current pathway collections. This enabled the prioritization of curation efforts.

#### Single-cell transcriptomic data analysis in epithelial cell types of COVID-19 patients

In this section, we provided gene expression analysis to explore differential expressed genes (DEG) on scRNAseq in specific epithelial cell populations in the COVID-19 patient group (moderate, severe, and critical cases), comparing with isolated epithelial cells from the lungs of healthy subjects. An exploratory gene expression data was carried out on single-cell RNAseq analysis of bronchoalveolar lavages from nine COVID-19 patients, three moderate cases, one severe case, and five critical cases (GSE145826) from (13). To obtain high confidence of differential expressions in three different groups, single-cell RNAseq data of isolated epithelial cells (DAPI−, CD45−, CD31−, CD326+) from control lung explant tissue of nine health subjects was chosen as a healthy control specific for epithelial cell types (14). All filtered samples were merged in only one filtered gene-barcode matrix and analyzed with R package Seurat v.3 (78). In parameter settings, the first 50 dimensions of canonical correlation analysis (CCA) and principal component analysis (PCA) were used. Moreover, the filtered gene-barcode matrix was first normalized using ‘LogNormalize’ methods with default parameters. UMAP was performed on the top 50 PCs for visualizing the cells, while clustering was performed on the PCA-reduced data for clustering analysis with Seurat v.3. The resolution was set to 0.5. A UMAP embedding represents the distribution of major cell types in the single-cell RNAseq database (**Fig. S6**). The epithelial cell group (TPPP3, KRT18), directly infected by SARS-CoV-2, was analyzed for every patient group. At first, the classification was provided, following these gene markers, as reported in (13): macrophages (CD68), neutrophils (FCGR3B), myeloid dendritic cells (mDCs; CD1C, CLEC9A), plasmacytoid dendritic cells (pDCs; LILRA4), natural killer (NK) cells (KLRD1), T cells (CD3D), B cells (MS4A1), plasma cells (IGHG4) and epithelial cells (TPPP3, KRT18). For the finest cell annotation of epithelial cell types, specific gene markers were used as reported in the Human Protein Atlas database (https://www.proteinatlas.org/), and markers of health epithelial cells reported by Deprez and colleagues (79) (10.1164/rccm.201911-2199OC) and extracted. In particular, ciliated cells (CFAP157, FAM92B; SARS-CoV-2-infected” cells 15.5%), Secretory cells (BPIFB1, SCGB1A1, SCGB3A1; SARS-CoV-2-infected” cells 6.4%), Suprabasal cells (KRT5, SERPINB4, KRT19, COVID19 cells 37.7%), Alveolar Type 1 cells (AGER, CAV1, EMP2, SARS-CoV-2-infected” cells 6%), Basal cells (KRT5, KTR15, COVID19 cells 11.2%). Alveolar Type 2 cells were not included because of an unbalanced ratio of cell sample size between COVID-19 cases and healthy control (SARS-CoV-2-infected” cells <2%; see **table S1** for a detailed summary of all cell types). The balanced sample size of cells allowed us to compare these two groups. For epithelial cell groups, differential gene expression analysis between patients and specific cell control was carried out. A differential gene expression analysis for all clusters was performed using the FindMarkers function in Seurat v.3, imposing a statistical threshold of 0.05 % FDR, average |logFC| > 1 and the difference between PCs >0.25, in order to maximally increase confidence in the results.

#### Integrative pathway modeling using C19DMap diagrams and RNAseq data from COVID-19 patients

The HiPathia algorithm allows modeling the behavior of signaling pathways, described as directed graphs that connect receptor proteins to effector proteins through a chain of activations and inhibitions exerted by intermediate proteins. HiPathia treats the pathways as if they were composed of elementary circuits, each circuit defined as the sub-pathway, or chain of proteins, connecting receptors to effectors. HiPathia uses expression values of genes as proxies of the levels of activation of the corresponding proteins in the circuit (80). To estimate the activity of a given circuit, an arbitrary signal value is transmitted through the nodes and is modulated by the activity values of the intervening proteins until it reaches the final effector protein, which is annotated with the functions that it triggers in the cell (16). These circuit activation values can be between conditions to obtain profiles of differential signaling and differential functional activity. The first version of the C19DMap has been implemented in the CoV-HiPathia version (81). In addition, extracted SIF files from SBML qual files using CaSQ (26) can be imported to HiPathia containing the Activity Flow (AF) structure of the Process Description (PD) diagrams, enabling new disease maps to be modeled as they are built thus permitting their exploration and analysis. In order to test the methodology, a public RNAseq dataset of nasopharyngeal swabs from 430 individuals with SARS-CoV-2 and 54 negative controls (17) (GSE152075) was used. First, the RNA-seq gene expression data were normalized with the Trimmed mean of M values (TMM) normalization method using the edgeR R package (82). Then, within the CoV-Hipathia web tool (81), the HiPathia algorithm requires the expression data to be rescaled between 0 and 1 for the calculation of the signal. Finally, quantile normalization using the preprocessCore R package (80) was carried out. The normalized gene expression values were used to calculate the level of activation of the sub-pathways, and then a case/control contrast with a Wilcoxon test was used to assess differences in signaling activity between the two conditions: SARS-CoV-2-infected and normal control nasopharyngeal tissue.

### 4.2 Dynamical modeling at the molecular, cellular, and multicellular levels

#### Dynamical modeling of type I IFN responses in SARS-CoV-2 infection

##### Type I IFN model development and computational validation

We used the type I IFN molecular map as a scaffold and auto-generated the dynamic model using the CaSQ tool. We utilized seven biological scenarios from the scientific literature to evaluate the model’s behavior.

##### Global sensitivity analysis

We simulated the model in Cell Collective (30) using varying activity levels of each input. We determined the input-output association using activity levels of 1000 randomly-generated simulations as previously used by our group (83). We performed probabilistic global sensitivity analysis based on the partial correlation coefficient (PCC) using the “sensitivity” package (https://cran.r-project.org/web/packages/sensitivity/sensitivity.pdf) in R (R Core Team, 2016) on data obtained from Cell Collective. It shows the impact of change in the input variable (independent variable) on the output variable (dependent variable) while considering and removing the linear effect of other input variables on the output variable (84). The script used in this analysis is available in our shared GitLab repository (https://git-r3lab.uni.lu/computational-modelling-and-simulation/analysis/-/blob/master/IFN1_modelling/Global_Sensitivity_analysis_of_IFN_model.R).

##### Sensitivity analysis against overexpression and knockouts

The sensitivity of biomolecules was calculated against knockout and overexpression perturbations. The sensitivity values were quantified in macro values for each biomolecule. The bitwise distances were calculated for each biomolecule in the same macro class. The highest sensitivity values were then simulated in Cell Collective. The methodology of the algorithm used to calculate the sensitivities against knockout and over-expression perturbations is described in FairdomHub (https://fairdomhub.org/data_files/4090), and the used script that generates the result is available in our shared GitLab repository (https://git-r3lab.uni.lu/computational-modelling-and-simulation/analysis/-/blob/master/IFN1_modelling/IFN1_sensitivity_against_mutations.R).

##### Input propagation for calculating stable states

The IFN model has 55 input components. These input components always maintain their activity level as they have no upstream regulators, and their initial configuration plays a vital role in the potential outcome. To eliminate unrealistic input configurations, we consider here that all inputs representing viral components share a common state. To encode this constraint, we introduce an additional input node controlling this group of components. We applied the same approach to inputs associated with the immune response and IFN secretion. In the resulting model, only six inputs remain, these three meta-inputs and three components representing drugs (GRL0617, Azithromycin, and MNS). Using this modified model, we identified 128 stable states. The absence of other stable patterns suggests that this model does not generate stable oscillations. We selected four output components to assess the obtained phenotypes (viral replication, antiviral response, inflammation, and secretion of IFNA1). The projection of the 128 stable states on these four outputs gave six distinct signatures among the 16 possibilities. All signatures lacked IFN secretion and exhibited either viral replication or antiviral response (or both). We then studied in more detail a set of 8 input conditions that cover different biological scenarios of the type I IFN pathway with or without the infection and in the presence or absence of drugs (**Table S2**). In these conditions, the propagation of the input values was sufficient to control most components of the model, and in particular, all selected output components. Studies in patients with COVID-19 with various degrees of severity showed hampered IFN-I responses in patients with severe or critical COVID-19. These patients had low levels of IFN-I and ISGs, and increased production of TNF-, IL-6-, and NF-κB-mediated inflammation.

### Integration of the Type 1 IFN, the ACE-ACE2 axis, and the NLRP3 inflammasome curated pathways into a macrophage-specific Boolean model

Three diagrams in the C19DMap repository were selected: the Type 1 IFN, the ACE-ACE2 axis, and the NLRP3 inflammasome. These diagrams were converted into SMBL qual formats using the CaSQ tool (26) and then processed in GINsim (60). Once processed, the pathway modules were integrated into a COVID-19-specific macrophage model. Phenotypic nodes were added to easily link the biomarkers with a biological process by way of an associated GO term name. Next, the functionality and behavior of the COVID-19 macrophage model were evaluated in a stable state analysis (attractors) performed with the following stimulatory conditions: inflammatory microenvironment, virus infection, and both.

### Multiscale and multicellular simulation

We incorporated two Boolean models into a multiscale simulator that consists of the infection of a patch of lung epithelium by SARS-CoV-2 and the immune cells that are recruited (43): macrophages, neutrophils, dendritic cells, CD4- and CD8-T-cells. We expanded this simulator with our tool, PhysiBoSS (85), which incorporates MaBoSS (86), a tool that stochastically simulates Boolean models, into PhysiCell (87), a tool that uses agent-based modeling to simulate cells and their surrounding environment, and their interplay. Two Boolean models were used: first, the epithelial apoptosis model was converted from the map to the model using CaSQ (26) and the C19DMap project (https://fairdomhub.org/models/712) (76). We modified the apoptosis model to capture mechanisms such as BAX activating the apoptosome complex and included output nodes as readouts. We also connected inputs and outputs to different variables in the population model, such as the *Virus_inside* node, which depends on the number of virions inside a cell, or the *Tcell_attached* node, which depends on the attachment of a T-cell to the epithelial cell (**Fig. 10C**). Second, we included the macrophage-specific Boolean model developed for this work. As with the apoptosis model, we connected the models’ inputs and outputs to relevant variables from the agents. For instance, activating the *Apoptotic_cell* node upon encountering an apoptotic epithelial cell, activating the *SARS_CoV_2* node upon encountering a virion, or activating the interferon Boolean nodes when the interferon roaming in the environment is above the detection threshold. Likewise, when *Neutrophil_recruitment, CD4_Tcell_activation* or *CD8_Tcell_activation* nodes are ON, proinflammatory cytokines are released. We found perturbations in the Boolean model that enhanced the recruitment of immune cells and the commitment to apoptosis using our pipeline of tools (44) that uses MaBoSS to simulate stochastic trajectories. Knocking out FADD in the epithelium cells blocked the commitment to apoptosis, which was expected from the regulation of that node. Interestingly, we found that knocking out p38 in the macrophages increased the recruitment of immune cells by 10%.

### Pharmacogenomic analysis

We obtained the list of proteins in the C19DMap as well as lists of proteins targeted by drugs and chemicals from annotations from the AILANI COVID-19 research assistant (https://ailani.ai) based on an NLP pipeline (88), INDRA (Integrated Network and Dynamical Reasoning Assembler) (1), and from the Clinical Trials DB. We used information from the cross-references from DrugBank (72) to map ChEBI and PubChem identifiers to DrugBank identifiers. We further enriched the list of drug/chemical targets using the information from DrugBank (accessed June 2022). A list of 16 drugs used for the treatment of COVID-19 was obtained from (89), and their targets were obtained from DrugBank. After merging the lists, a final dataset of 1,476 drugs and chemicals (identified by DrugBank IDs) and 1,120 drug targets (identified by NCBI Gene Id) was obtained. Information on pharmacogenomic variants for the drug targets was retrieved from PharmGKB (90) (accessed on Feb 14, 2021). For each gene that encodes a drug target, the list of variants with pharmacogenomic annotations that are significant and are annotated to a dbSNP identifier was retrieved. We used the cross-references from PharmGKB to map the PharmGKB drug accessions to DrugBank identifiers. Data on the allelic frequency of the pharmacogenomic variants were retrieved from The Genome Aggregation Database (gnomAD) (91) (version 2.1.1). gnomAD is a resource developed by an international coalition of investigators with the goal of aggregating and harmonizing both exome and genome sequencing data from a wide variety of large-scale sequencing projects and making summary data available for the broader scientific community. To aggregate the data on the pharmacogenomic impact and allelic frequency of the variants, we computed a modified version of the Cumulative Allele Probability (CAP) and the “Drug Risk Probability” (DRP) score (47). The CAP score considers the number of pharmacogenomic variants and their frequency in the population for a specific gene. The DRP score combines the CAP scores for all drug target genes for a specific drug. The code to compute the CAP and DRP scores is available at https://github.com/jpinero/pharmacogenomics_covid19_minerva_map/.

### Topological analysis

For each of the available pathways, we calculated values for a set of 17 network centrality measures as implemented in Vanted’s Centilib extension (92). Taking into account the results of correlation analysis and the requirements of centrality calculation on the network structure, such as connectivity, we restricted the 17 measures to a base set of ten measures (Eccentricity, Degree, Eigenvector, HITSAuths, Current Flow Betweenness, Radiality, Stress, Shortest Path Betweenness, Centroid Rank, Closeness)(93). For these measures, we calculated the values for each network node (excluding reactions) and provided rankings of nodes for each measure per network. Additionally, we computed aggregated rankings using the residual sum of squares for each node per network as well as on the aggregated network. The results from our centrality calculations can also be explored and put in context using the software LMME-DM (https://github.com/LSI-UniKonstanz/lmme-dm) that was developed as part of the C19DMap project. It follows an overview and detail approach showing an overview graph containing one node per pathway, and a detailed pathway view, including the detailed crosstalks. The centrality values can now be mapped on both the size and the color of the nodes (see **Fig. 10**).

### Orthoinference process for converting from SARS-CoV-1 to SARS-CoV-2 diagrams

The standard orthoinference process is used to infer reactions electronically in fifteen evolutionarily divergent eukaryotic species for which high-quality whole-genome sequence data are available. Eligible reactions are checked to determine whether each involved protein has at least one homologous protein in the reaction’s input, output, and (if present) catalyst in the organism undergoing inference. If a human reaction involves a complex, at least 75% of the accessioned protein components of the human complex must have homologous proteins in the model organism. The first (V74) draft of this SARS-CoV-2 pathway consists of 101 reactions involving 489 molecular entities (279 proteins, 12 RNAs, and 198 others) and is supported by citations from 227 publications. Reactome developed a computational triaging strategy to review and identify publications appropriate for manual curation (66,100 SARS-CoV-2 articles on PUBMED, tallied on 30/October/2020).

## Acknowledgements

The authors would like to acknowledge all members of the COVID-19 Disease Map community for fruitful exchanges and discussions, the LCSB and Elixir Luxembourg for the hosting of the C19DMap resource and repository, the FAIRDOMHub community for supporting our project, and members of the SysMod, CoLoMoTo, BioModels and COMBINE communities for fruitful exchanges and feedback.

## Author contribution statement

Planned and coordinated the project: AN, MO

Advised the project as domain experts: HK, EB, AV, OW, RS, CA, CE, ELW, JD, JSR, RB, FS, JSB, RP, PdE, MK, MEG

Topological analysis: FS, KK, MA, FB, TC

AI-assisted biocuration, text mining and data retrieval for drugs and drug targets: DM, AB, BMG, JAB, AF, AL

Created harmonized content, ensured interoperability of infrastructure and technical support: MO, AN, SS, PG, MG

Contributed code: AH, ANa, MK, AMo, MO, AF, TGY, YH, NFH, FH, NP, FE, VN, MPdL, BLP, SS, JP, TC, AV, AD, MA, KK, AV, AD, FM, MEM, MPC, KR

Omic data and pathway analysis: MK, FH, NP, FE, AF, TGY, YH, NEH, JSR, AD, AV, FM, MPC, KR, MEM

Molecular, Cellular and Multicellular modeling: AN, SSA, TH, BLP, ANa, SS, VS, MK, VB, MFF, ET, LC, VN, AMo, MPdL, AV

Pharmacogenomics analysis: JP, LIF, AN, ELW Neo4j implementation: AMa, IB, AR, YJ, AL

Protocol and advice on contextualization of the diagrams: MEG, PdE, BJ, VS, GW, IK, MGLC, JMR, MLA, BDM, LP, LM, MOM, DABR, RWO

Analyzed and synthesized the results: AN, MO, AF, MK, FM, MA, KK, TC, FS Designed the Figures: AN, MK, AF, AV, FM, MEM, KR, MPC, BLP, VB, MFF, MA

Wrote the draft of the manuscript: AN, MO

Contributed significantly to the structure of the manuscript: AN, MO, CE, MEG, DM, JSB, AF, MLA, MK, NFH, FM, AMo, BMG, JP, LIF, RWO, MOM

Revised, edited, and contributed text to the manuscript: all authors Read and approved the final version of the manuscript: all authors

## Data availability statement

All data, code and files are available in our GitLab repository https://gitlab.lcsb.uni.lu/computational-modelling-and-simulation

Pharmacogenomics analysis:

https://github.com/jpinero/pharmacogenomics_covid19_minerva_map/blob/main/drp_score_drugs.tsv https://github.com/jpinero/pharmacogenomics_covid19_minerva_map/blob/main/cap_score_genes.tsv

Topological analysis:

https://github.com/LSI-UniKonstanz/lmme-dm

Fairdomhub: https://fairdomhub.org/projects/190

## Supplementary material

All supplementary tables and figures are included in a pdf file, and supplementary table 5 is provided as a separate excel file.

