## Supplementary Material for "A versatile and interoperable computational framework for the analysis and modeling of COVID-19 disease mechanisms"

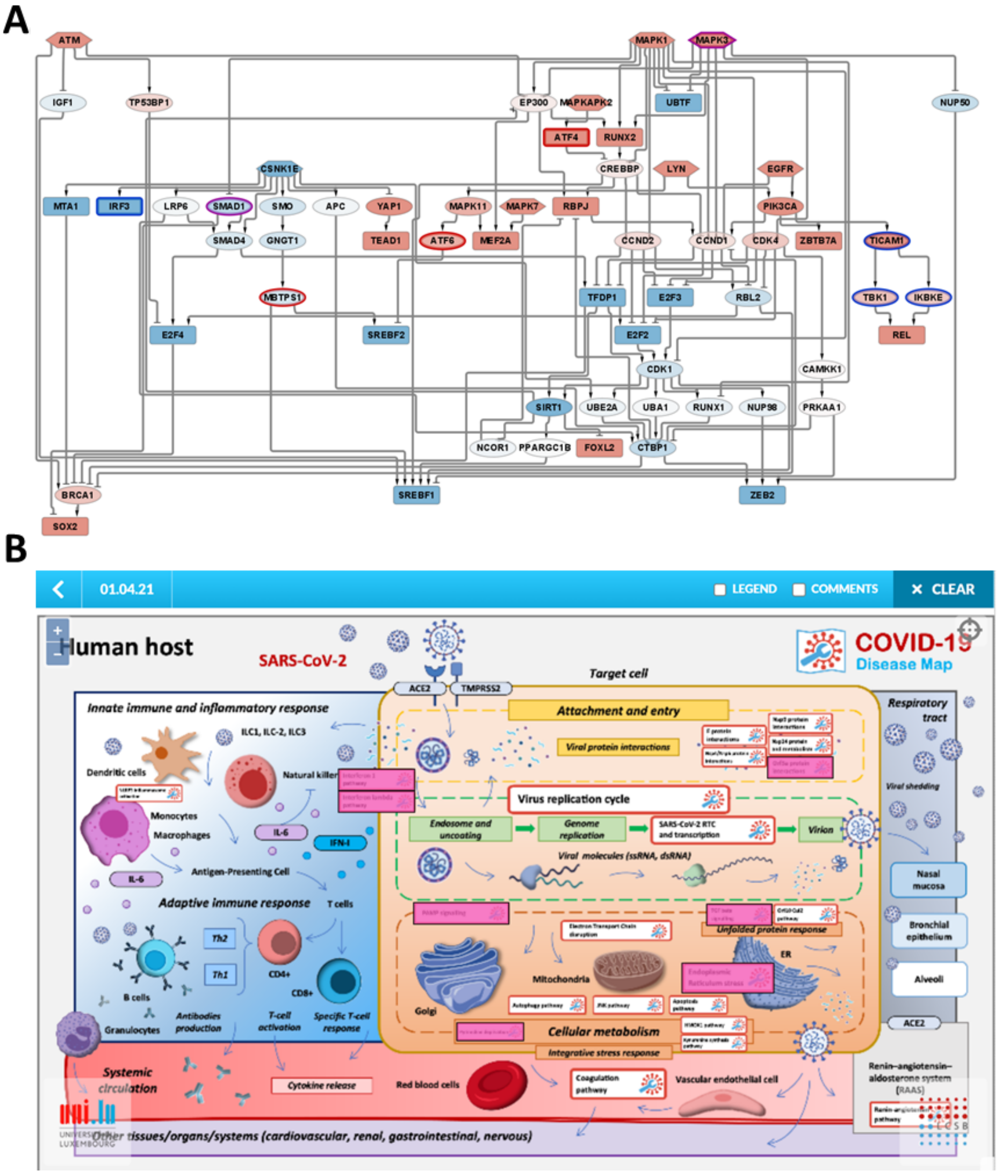


***Supplementary Figure 1A)*** Carnival network of mechanistic hypotheses connecting the top 10 deregulated kinases with the top 30 deregulated TFs. Kinases are represented as hexagon-shaped nodes, TFs as rounded rectangles, and intermediary signaling proteins as elliptical nodes. Up-regulated nodes are indicated in red while down-regulated nodes are colored in blue. Activatory reactions are indicated with arrow-shaped edges whereas inhibitory ones are represented by T-shaped edges. Bold outlines denote proteins already present in the COVID-19 Disease Map. Blue outlines denote PAMPs and Interferon-1 pathway members, red outlines are ER stress pathway members and purple outlines indicate TGFbeta pathway members of the pathways manually curated by the COVID-19 Disease Maps community. ***B)*** Highlighted C19DMap diagrams containing deregulated kinases and TFs are shown in dark pink. Users can click on the highlighted boxes and enter the diagram.

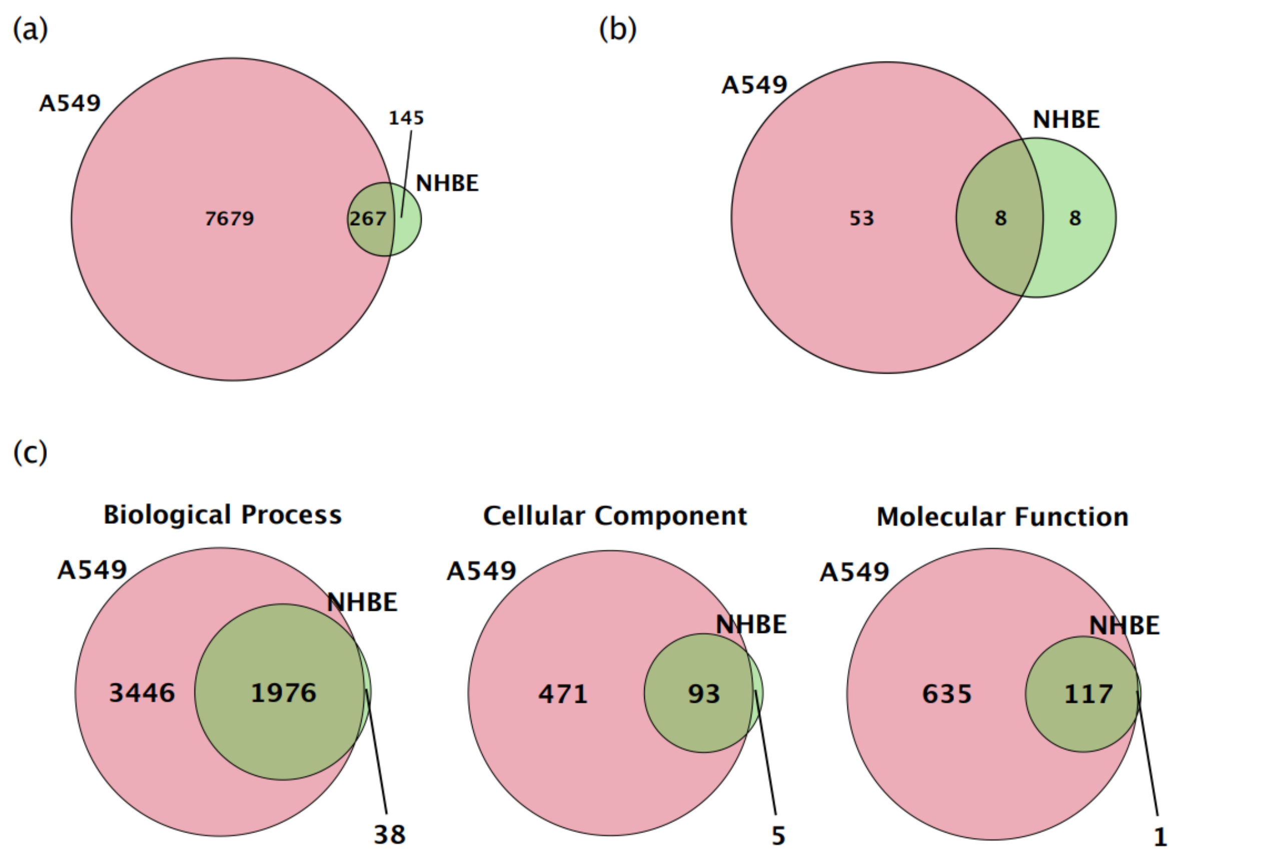


***Supplementary Figure 2:*** Overlap of (a) DEGs ,(b) TFs, and (c) GO terms between coronavirus-infected NHBE and A549 cells by LAMP and GO Enrichment analysis.

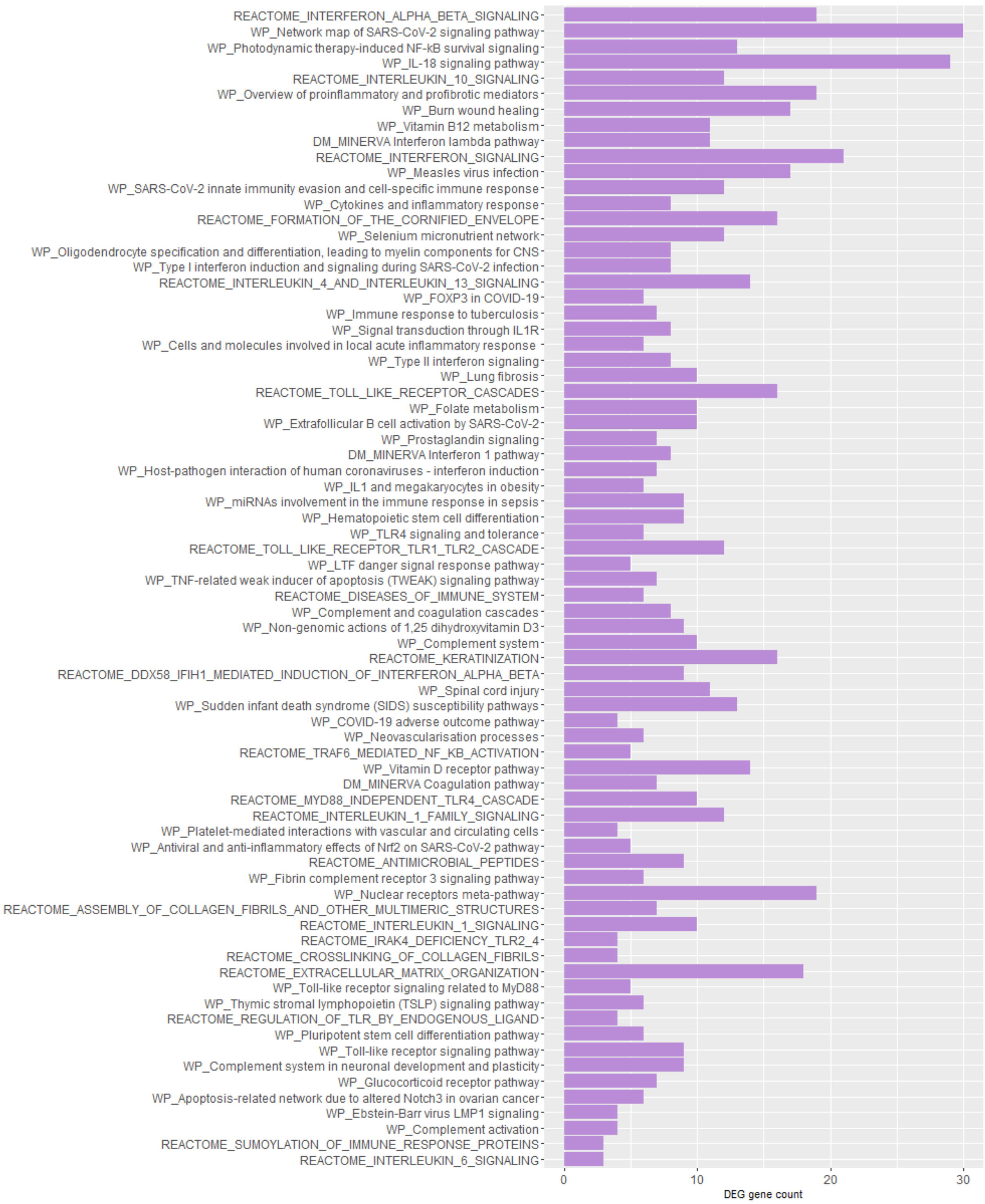


***Supplementary Figure 3.*** C19DMap (DM), WikiPathways (WP) and Reactome pathways significantly enriched for DEGs in SARS-CoV-2-infected NBHE cells.

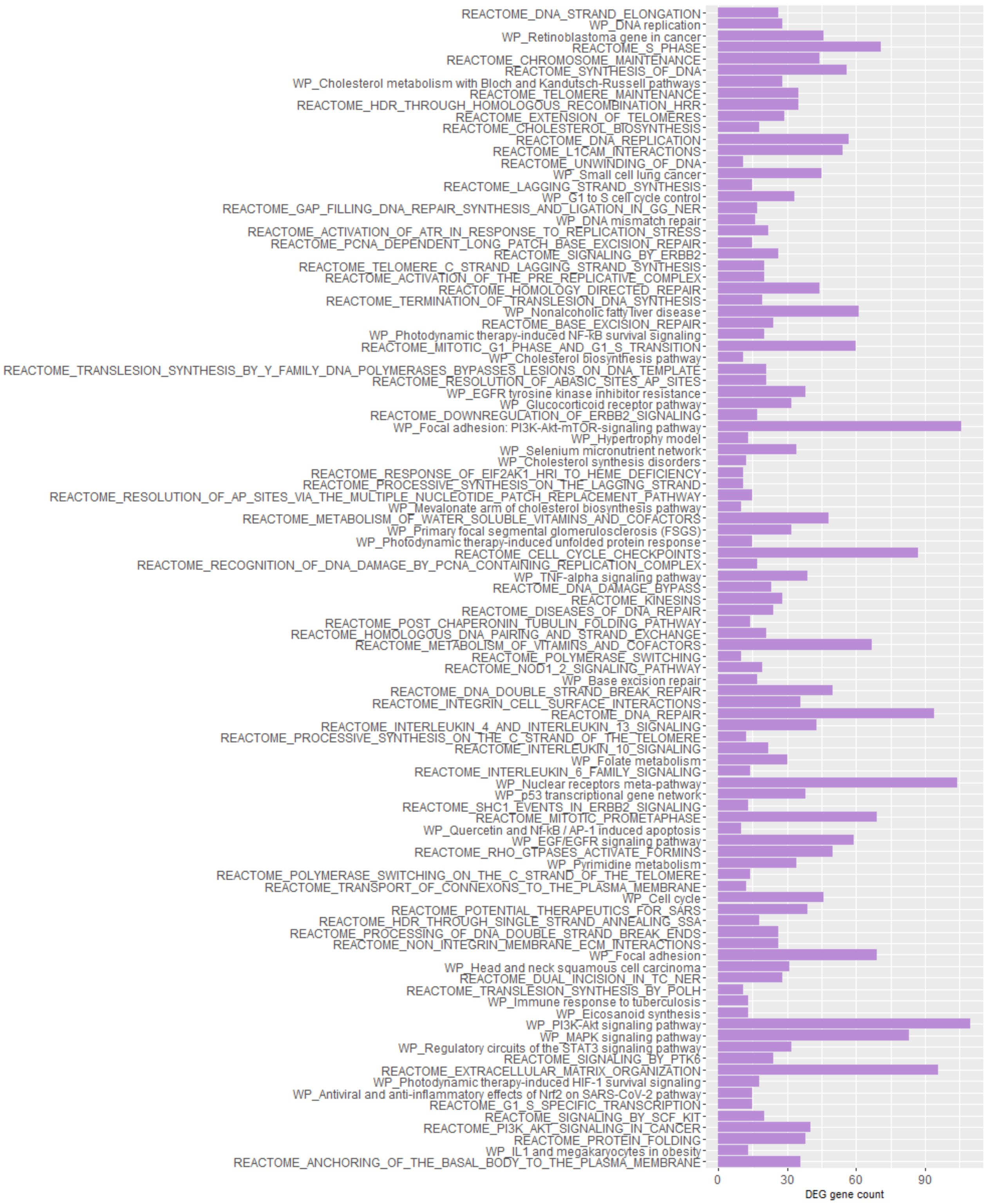


***Supplementary Figure 4.*** C19DMap (DM), WikiPathways (WP) and Reactome pathways significantly enriched for DEGs in SARS-CoV-2-infected A549 cells.

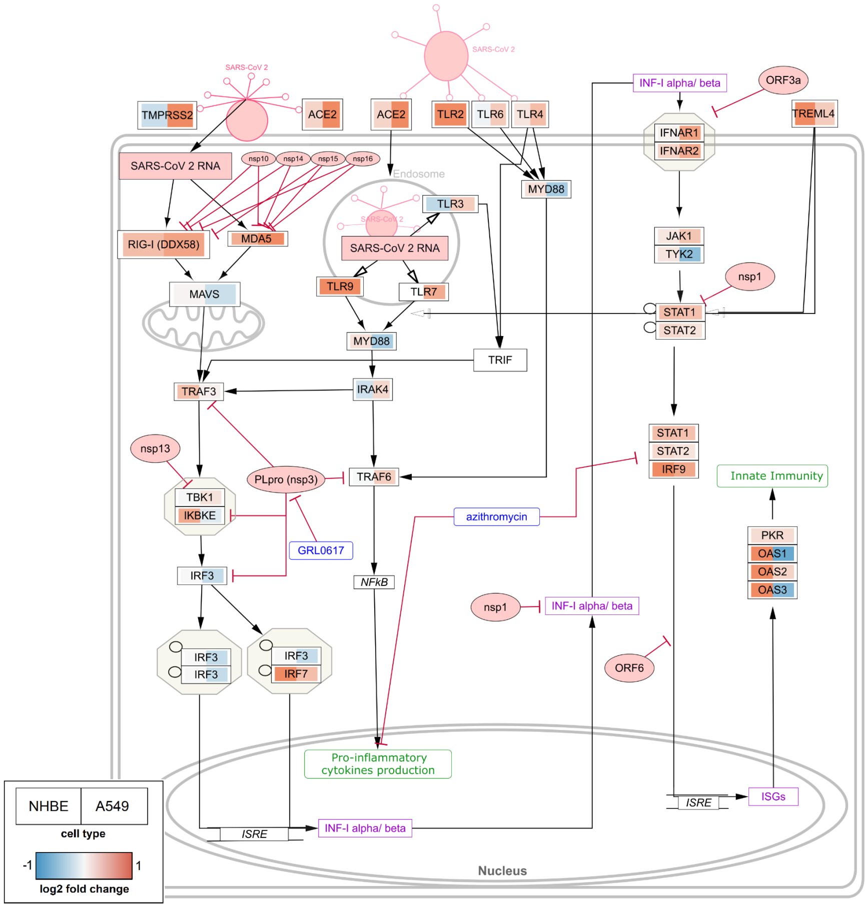


***Supplementary Figure 5: Data visualization on the type I interferon induction and signaling during SARS-CoV-2 infection.*** Transcriptomics data can be easily visualized on the altered pathways as shown here on the Type I interferon induction and signaling during SARS-CoV-2 infection pathway (wikipathways:WP4868). The boxes are split in half, left side shows the expression change in NHBE cells and the right side shows the change in A549 cells. The log2 fold change is visualized, red indicating up-regulation, blue indicating down-regulation. Grey nodes are not measured in the dataset.

***Supplementary Table 1:*** Differential cell type-specific gene expression between COVID-19 patients and healthy controls. COVID-19 cell sample size (%) represents absolute count and percentage of cell types in COVID-19 patients-derived samples; Overexpressed DEGs and Downexpressed DEGs represent numbers of overexpressed and downexpressed DEGs, respectively, in COVID-19 patients vs. healthy controls. 26 COVID-19 overexpressed DEGs were shared between all cell types.

| | **COVID-19 cell sample size (%)** | **Overexpressed DEGs** | **Downexpressed DEGs** |
|---|---|---|---|
| **Ciliated cells** | 952 (13.5%) | 61 | 93 |
| **Alveolar cells type 1** | 139 (5.7%) | 183 | 81 |
| **Basal cells** | 469 (10.1%) | 166 | 77 |
| **Secretory cells** | 237 (6%) | 83 | 42 |
| **Suprabasal cells** | 958 (27.3%) | 94 | 116 |

A)

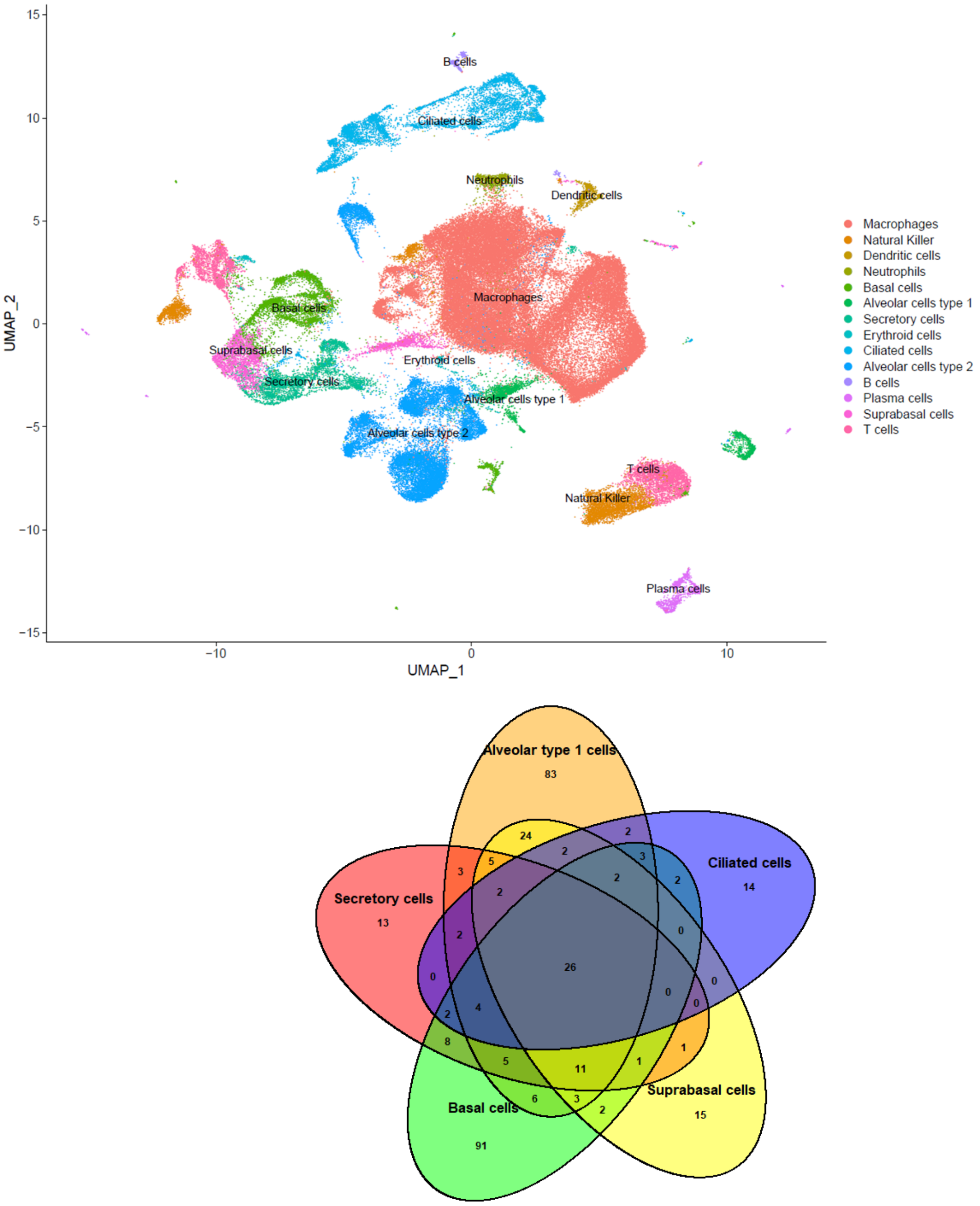


B)

***Supplementary Figure 6.*** *A) UMAP of both whole data sets (GSE145926 and GSE160664), where all cell types were highlighted and reported. B)Overlap of overexpressed (positive) DEGs among all detected lung epithelial cell types of COVID-19 patients relative to healthy controls.*

***Supplementary Table 2.*** *26 DEGs shared among all detected lung epithelial cell types. The genes in red belong to Type I interferon induction and signalling during SARS-CoV-2 infection (WikPathway WP4868).*

| | |
|---|---|
| ATP6V0C | MX1 |
| OAS1 | SRGN |
| MX2 | IFIT2 |
| IFI44L | IFIT3 |
| OAS3 | IFI44 |
| EPSTI1 | GBP1 |
| IFITM1 | CCL2 |
| OAS2 | RSAD2 |
| XAF1 | IRF7 |
| IFIT1 | MT-ATP8 |
| ISG15 | NME2 |
| SAMD9 | IFI6 |
| TAOK1 | STAT1 |

***Supplementary Table 3.*** Significant pathways-circuits activity values after Wilcoxon test comparison between 430 SARS-CoV-2-infected and 54 non-infected individuals. The results are obtained after running CoV-Hipathia web tool with GSE152075 dataset.

| Pathway: Effector Circuitname | UP/DOWN | statistic | FDR-p.value | Fold Change | logFC |
|---|---|---|---|---|---|
| Nsp9 protein interactions: EIF4H | DOWN | -7.56E+00 | 6.09E-12 | 6.29E-01 | -6.68E-01 |
| **Interferon lambda pathway: STAT1,STAT2,STAT3** | **UP** | **7.16E+00** | **5.77E-11** | **2.01E+00** | **1.01E+00** |
| **Interferon 1 pathway: OAS1,OAS2,OAS3** | **UP** | **6.89E+00** | **2.77E-10** | **2.88E+00** | **1.53E+00** |

| | | | | | |
|---|---|---|---|---|---|
| Nsp9 protein interactions: MYCBP2 | UP | 6.57E+00 | 1.62E-09 | 1.26E+00 | 3.32E-01 |
| Nsp9 protein interactions: COMT | DOWN | -6.54E+00 | 1.62E-09 | 4.13E-01 | -1.28E+00 |
| Nsp9 protein interactions: DCAF7 | UP | 5.62E+00 | 4.65E-07 | 1.23E+00 | 2.93E-01 |
| Nsp9 protein interactions: CYB5R3 | DOWN | -5.57E+00 | 5.35E-07 | 7.47E-01 | -4.20E-01 |
| JNK pathway: JUN,JUND | DOWN | -5.47E+00 | 8.14E-07 | 5.83E-01 | -7.78E-01 |
| Nsp9 protein interactions: MIB1,DLL1 | DOWN | -4.83E+00 | 1.38E-06 | 1.31E-01 | -2.93E+00 |
| E protein interactions: STOML3,ASIC1 | DOWN | -5.12E+00 | 2.10E-06 | 3.48E-01 | -1.52E+00 |
| **Renin-angiotensin pathway: LNPEP** | **UP** | **4.87E+00** | **1.45E-05** | **1.25E+00** | **3.23E-01** |
| Nsp9 protein interactions: AP2M1 | DOWN | -4.56E+00 | 6.26E-05 | 8.04E-01 | -3.14E-01 |
| E protein interactions: CCNT1,CDK9 | DOWN | -4.52E+00 | 6.73E-05 | 5.00E-01 | -1.00E+00 |
| **Interferon 1 pathway: ISG15** | **UP** | **4.26E+00** | **1.81E-04** | **2.55E+00** | **1.35E+00** |
| Kynurenine synthesis pathway: AHR | UP | 4.28E+00 | 1.81E-04 | 1.18E+00 | 2.35E-01 |
| Nsp9 protein interactions: GFER | DOWN | -4.10E+00 | 1.98E-04 | 4.41E-01 | -1.18E+00 |
| Nsp9 protein interactions: IMPDH2 | DOWN | -4.13E+00 | 3.10E-04 | 6.98E-01 | -5.19E-01 |
| Renin-angiotensin pathway: MAS1 | UP | 4.06E+00 | 3.50E-04 | 2.45E+00 | 1.29E+00 |

| | | | | | |
|---|---|---|---|---|---|
| Nsp4 and Nsp6 protein interactions: Nsp3:Nsp4:Nsp6 | DOWN | -3.73E+00 | 5.78E-04 | 5.55E-01 | -8.49E-01 |
| E protein interactions: CRB3,PATJ,MPP5 | DOWN | -3.69E+00 | 5.88E-04 | 4.71E-01 | -1.09E+00 |
| Coagulation pathway: MAS1 | UP | 3.85E+00 | 5.88E-04 | 1.28E+01 | 3.68E+00 |
| HMOX1 pathway: RBX1,KEAP1,CUL3 | DOWN | -3.87E+00 | 6.33E-04 | 2.73E-01 | -1.87E+00 |
| Orf3a protein interactions: HMOX1,ALG5,ARL6IP6 | DOWN | -3.46E+00 | 7.28E-04 | 2.45E-01 | -2.03E+00 |
| Nsp4 and Nsp6 protein interactions: DNAJC11 | DOWN | -3.63E+00 | 8.61E-04 | 5.67E-01 | -8.19E-01 |
| Nsp9 protein interactions: SEPSECS | UP | 3.65E+00 | 1.48E-03 | 1.66E+00 | 7.35E-01 |
| Nsp4 and Nsp6 protein interactions: TIMM29,TIMM22,TIMM10B | DOWN | -3.24E+00 | 1.70E-03 | 3.85E-01 | -1.38E+00 |
| Nsp4 and Nsp6 protein interactions: IDE | DOWN | -3.44E+00 | 1.77E-03 | 5.59E-01 | -8.40E-01 |
| Nsp9 protein interactions: MRPS5,MRPS2 | DOWN | -3.50E+00 | 2.13E-03 | 6.66E-01 | -5.86E-01 |
| Nsp9 protein interactions: FBLN5,LOXL1 | DOWN | -3.22E+00 | 2.68E-03 | 3.35E-01 | -1.58E+00 |
| Apoptosis pathway: CASP7 | UP | 3.37E+00 | 3.66E-03 | 1.34E+00 | 4.22E-01 |
| Nsp9 protein interactions: COPS6,EDN1 | DOWN | -3.00E+00 | 4.52E-03 | 4.26E-01 | -1.23E+00 |

| | | | | | |
|---|---|---|---|---|---|
| Nsp9 protein interactions: POLR2A,GTF2F2,POLR 2G | DOWN | -2.99E+00 | 9.43E-03 | 5.54E-01 | -8.53E-01 |
| Nsp9 protein interactions: BAG6,EDN1 | DOWN | -2.68E+00 | 1.51E-02 | 8.07E-01 | -3.10E-01 |
| Nsp9 protein interactions: AP2A2 | DOWN | -2.74E+00 | 2.65E-02 | 8.02E-01 | -3.18E-01 |
| Nsp9 protein interactions: RALA | UP | 2.70E+00 | 2.84E-02 | 1.31E+00 | 3.87E-01 |
| Nsp9 protein interactions: SBNO1 | UP | 2.64E+00 | 3.18E-02 | 1.06E+00 | 8.03E-02 |
| Nsp9 protein interactions: DDX10 | UP | 2.64E+00 | 3.18E-02 | 1.28E+00 | 3.53E-01 |
| Nsp9 protein interactions: UBQLN4,EDN1 | DOWN | -2.37E+00 | 3.18E-02 | 9.24E-01 | -1.13E-01 |
| PAMP signalling: ITCH | UP | 2.62E+00 | 3.26E-02 | 1.08E+00 | 1.18E-01 |
| Nsp9 protein interactions: ZNF503 | DOWN | -2.47E+00 | 3.39E-02 | 6.31E-01 | -6.63E-01 |
| Nsp9 protein interactions: CCDC86 | DOWN | -2.53E+00 | 3.42E-02 | 7.05E-01 | -5.05E-01 |
| Nsp9 protein interactions: TCF12 | UP | 2.53E+00 | 3.97E-02 | 1.04E+00 | 6.07E-02 |
| Nsp9 protein interactions: MAT1A,MAT2A,MAT2B | DOWN | -2.36E+00 | 4.34E-02 | 6.69E-01 | -5.81E-01 |
| Nsp9 protein interactions: COPS2,COPS4,COPS5 | DOWN | -2.44E+00 | 4.57E-02 | 6.84E-01 | -5.49E-01 |
| Nsp9 protein | DOWN | -2.45E+00 | 4.57E-02 | 8.06E-01 | -3.11E-01 |

| | | | | | |
|---|---|---|---|---|---|
| interactions: MEPCE,LARP7 | | | | | |
| Nsp4 and Nsp6 protein interactions: ATP6AP1 | DOWN | -2.41E+00 | 4.99E-02 | 8.33E-01 | -2.63E-01 |
| Nsp9 protein interactions: LARP4B | UP | 2.39E+00 | 5.07E-02 | 1.03E+00 | 4.31E-02 |
| Nsp9 protein interactions: ZNF503,DCAF7 | DOWN | -2.27E+00 | 5.07E-02 | 7.46E-01 | -4.23E-01 |

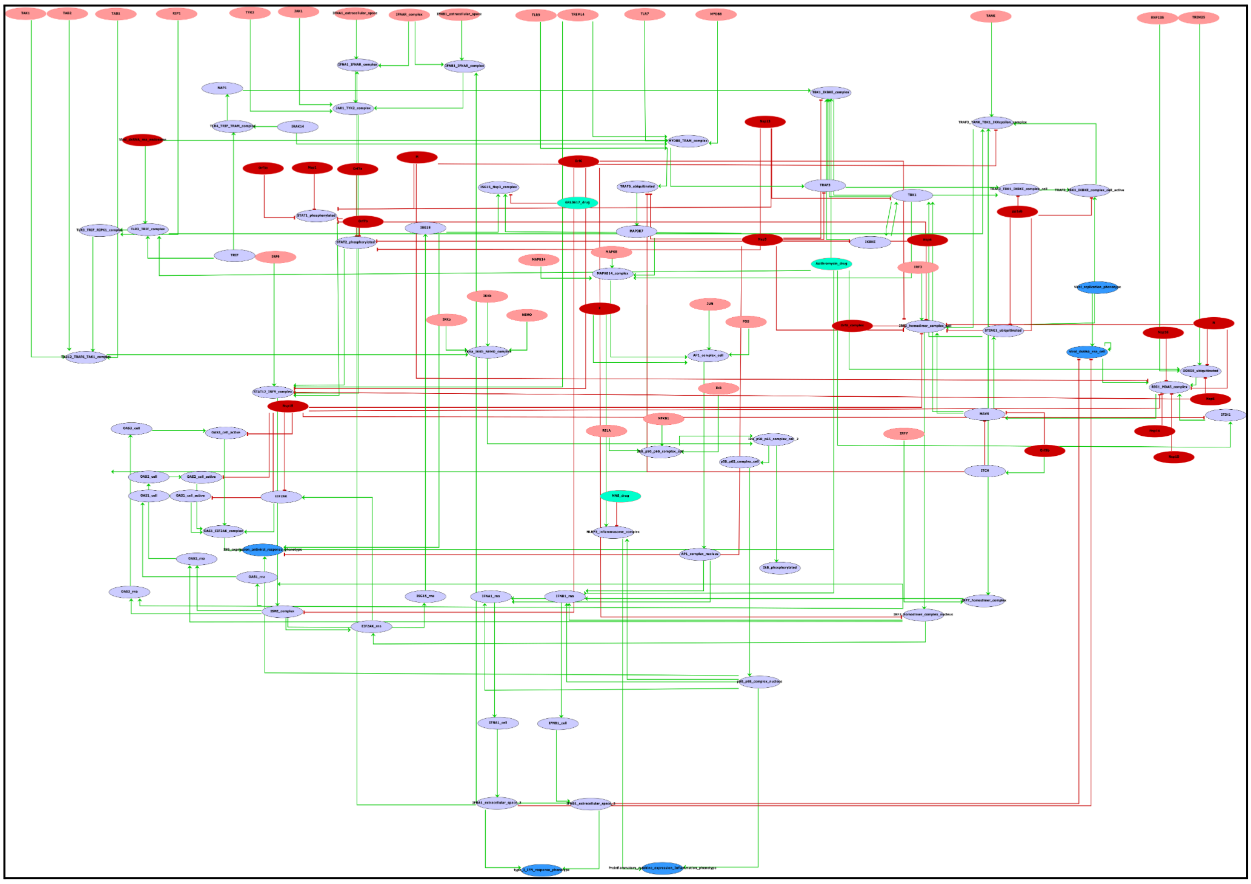

***Supplementary Figure 7:*** The Boolean network obtained after the processing of the CellDesigner XML file of the Type I IFN of the COVID-19 Disease Map repository with CaSQ 9.0.2. After post-processing, the final version of the model contains 121 nodes and 190 edges. Colour code: Red: Viral proteins, Turquoise: Drugs, Pink: host cell inputs, Purple: Host cell proteins, Light blue: Phenotypes of interest

*Supplementary Table 4:* Simulation scenarios and model behaviour.

| No | Biological Scenarios | Simulation results | Model performance (agreement/ partial agreement/ disagreement) |
|---|---|---|---|
| **1** | An unbalanced immune response, characterized by a weak production of type I interferons (IFN-Is) and an exacerbated release of proinflammatory cytokines, contributes to the severe forms of the COVID-19 disease. [PMID: 32726355] | When all of the viral components are active, the inflammation phenotype is increased. | Agreement |
| **2** | SARS-CoV-2-inhibition of type I IFN responses in infected cells leads to delayed or suppressed type I IFN responses, allowing the virus to replicate unchecked and induce tissue damage. [PMID: 33619493, 33097660] SARS-Cov-2 is a poor inducer of IFN-I response *in vitro* and in animal models compared to other respiratory RNA viruses. [PMID: 32726355] Multiple viral structural and non-structural proteins antagonize interferon responses, contributing to inflammation. [PMID: 33619493, 32346093] The suppression of interferon signalling is a mechanism widely used by SARS-CoV-2 in diverse tissues to evade antiviral innate immunity, and targeting the viral mediators of immune evasion may help block virus replication in patients with COVID-19. [PMID: 33140044] | When all of the viral components are active, the type 1 interferon immune response is inactive while the inflammation phenotype increases. We observe the inflammation reduction and activation of ISG expression_antiviral response_phenotype after deactivating all of the viral components. | Agreement |
| **3** | The viral ORF6, ORF8 and nucleocapsid proteins were potent inhibitors of the type I interferon signalling pathway, a | In the presence of the ORF6 and ORF8 complex, and nucleocapsid protein | Agreement |

| | | | |
|---|---|---|---|
| | key component for the antiviral response of host innate immunity. All three proteins strongly inhibited type I interferon (IFN-β) and NF-κB-responsive promoters. [PMID: 32589897] | (N), NFkB pathways and the type I interferon secretion are inactive. | |
| **4** | SARS-CoV-2 desensitizes host cells to interferon by inhibiting the JAK-STAT pathway.<br>[PMID: 33140044] | By increasing the viral components, we observe the decrease of STAT1 and STAT2_phosphorylated, which are part of the JAK/STAT pathway. | Agreement |
| **5** | The antiviral response is intensified by various signalling factors, including sensors and transcriptional regulators, which are themselves ISGs induced by ISGF3 and/or directly by the IRF3/IRF7 transcriptional activators. [PMID: 32726355]<br>The IRF3 gene is highly expressed in the activated CD4+ T cells of COVID-19 patients.[PMID: 32783921] | Interferon regulatory factor 3 (IRF3 homodimer complex_nucleus) is activated by adding Azithromycin drug to the viral components including orf7a/b, orf3a, viral dsRNA, Nsp1, Nsp13, and E protein.<br>(Initial condition of the IRF3 affects complex formation) | Agreement |
| **6** | Several interferon (IFN)-stimulated genes (ISGs; including ISG15, IFI44, IFI44L, and RSAD2) were specifically upregulated in PBMCs from COVID-19 patients, enhancing antiviral and immune-modulatory functions after viral infection. [PMID: 32783921] | When all of the viral components are active, the ISG expression_antiviral response_phenotype is inactivated; however, adding the Azithromycin_drug increases the ISG antiviral response.<br>ISG15 ubiquitin-like modifier is inactive in this condition; however, blocking orf6 and activating STAT1/2_IRF9_complex can upregulate the expression of the ISG15 by affecting ISRE_complex pathways. | Partial agreement |

| | | | |
|---|---|---|---|
| **7** | Inhibition of SCoV2-PLpro with GRL-0617 impairs the virus-induced cytopathogenic effect, maintains the antiviral interferon pathway and reduces viral replication in infected cells. [PMID: 32726803] | When all of the viral components are active, the type 1 interferon response is inactive; By increasing the GRL0617 drug, we didn't observe any effects on the antiviral interferon pathways or reduction of viral replication. The model suggests that applying GRL-0617 alone cannot impact this pathway. | Disagreement |

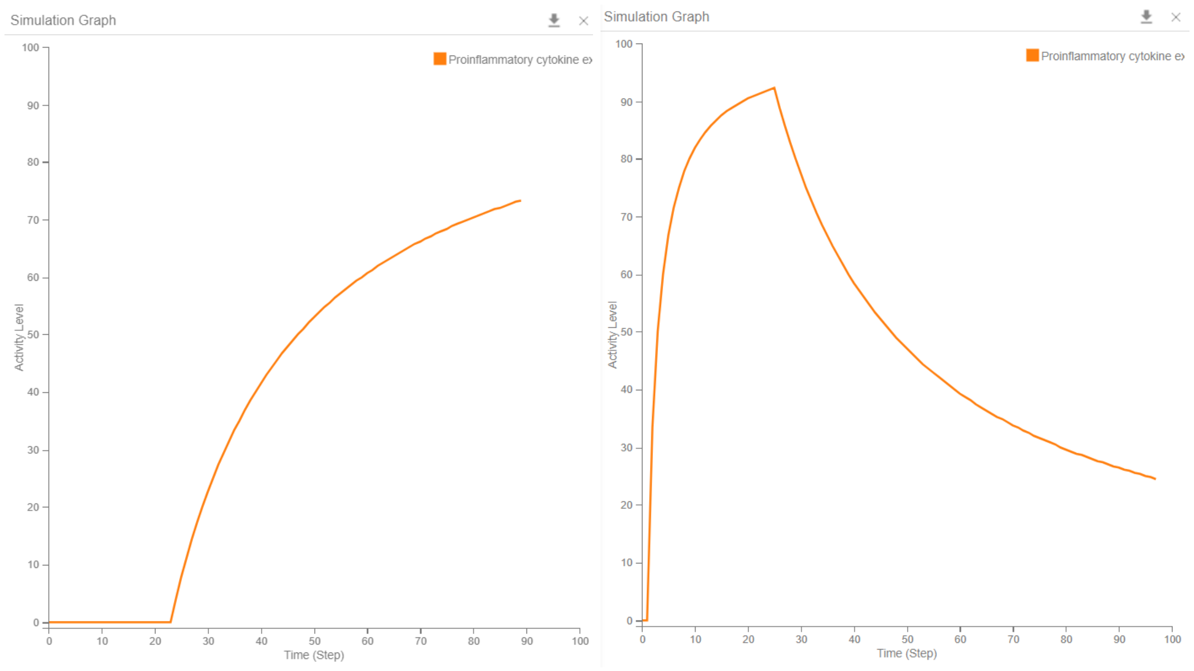


***Supplementary Figure 8.*** **Example of the *in silico* simulation in CellCollective.** The orange line presents the inflammation activity level. The left graph presents the increase of inflammation phenotype when viral components are activated after the time step 20. The right graph presents the decrease of inflammation phenotype in the presence of viral components upon addition of the GRL0617, Azithromycin, and MNS drugs to the system (at time step 20).

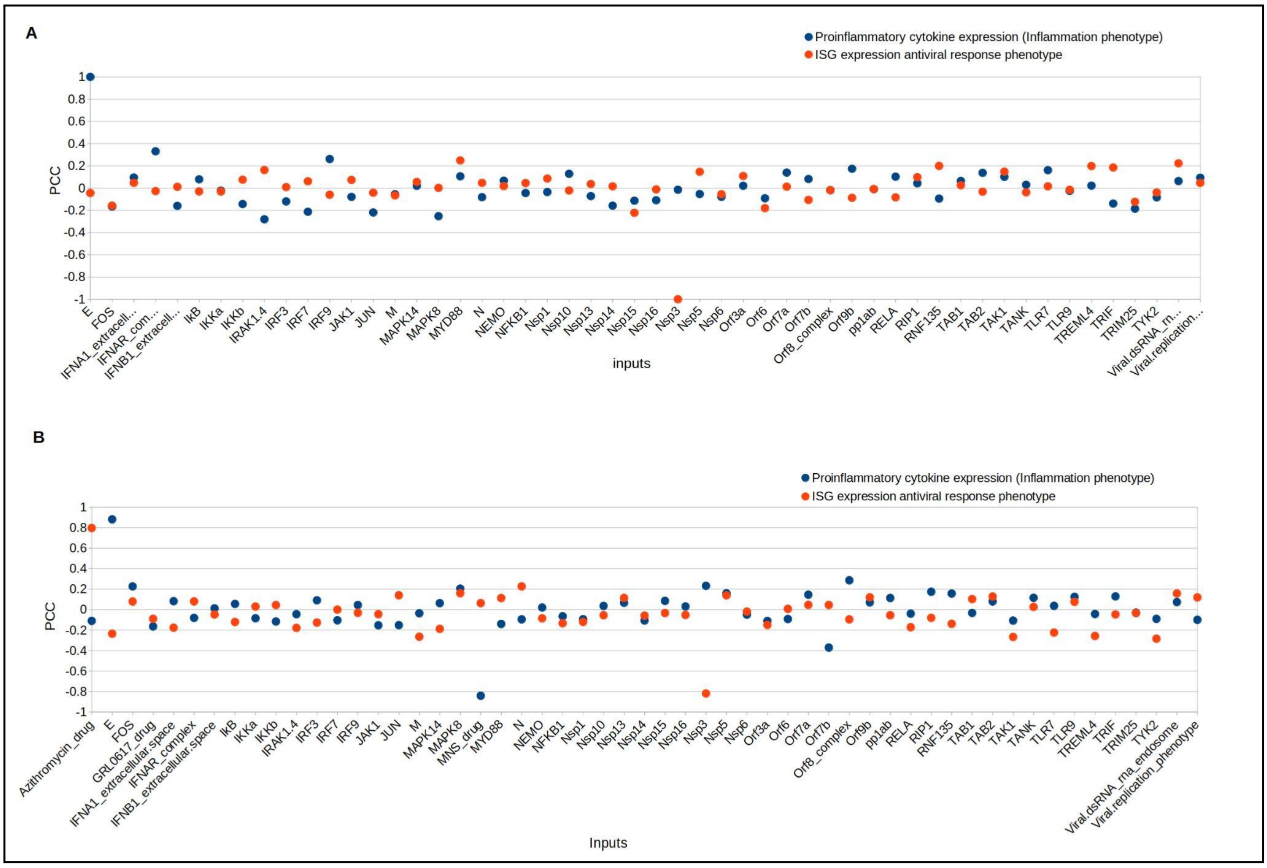


***Figure 9*** Sensitivity analysis showing the association between inputs (external components) and outputs (phenotypes). A) All the inputs, except drugs, were active. Inflammation phenotype has the highest positive association with protein E. The 'ISG expression antiviral response phenotype' has the highest association (negative) with Nsp3. B) All the inputs were active. The 'Inflammation phenotype' has the highest positive association with viral protein E and negative association with the MNS drug. An increase in viral proteins leads to higher inflammation. In contrast, the increase in MNS will reduce inflammation. PCC is a partial correlation coefficient measuring the strength of association between external inputs and output variables.

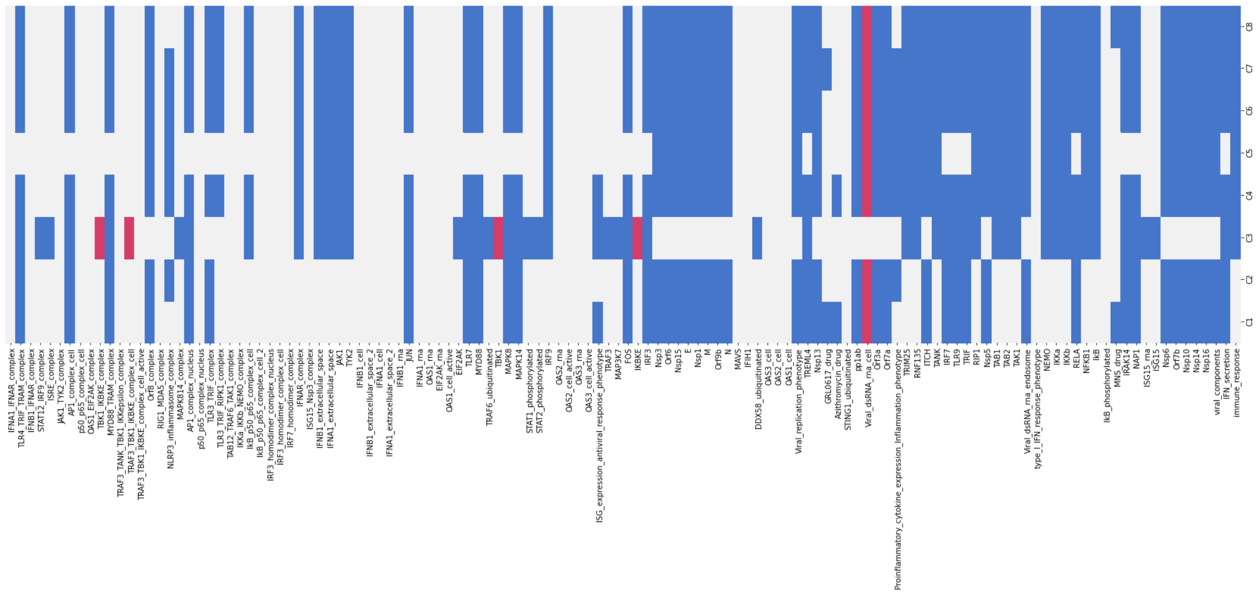

***Supplementary Figure 10.*** The result of input propagation can be visualized in the heatmap where lines represent components of the system and columns represent the 8 selected input conditions. A white cell denotes that the corresponding component is fixed at value 0 in this input condition. Likewise, a blue cell denotes that it is fixed at 1. Red cells denote components which are not fixed by input propagation.

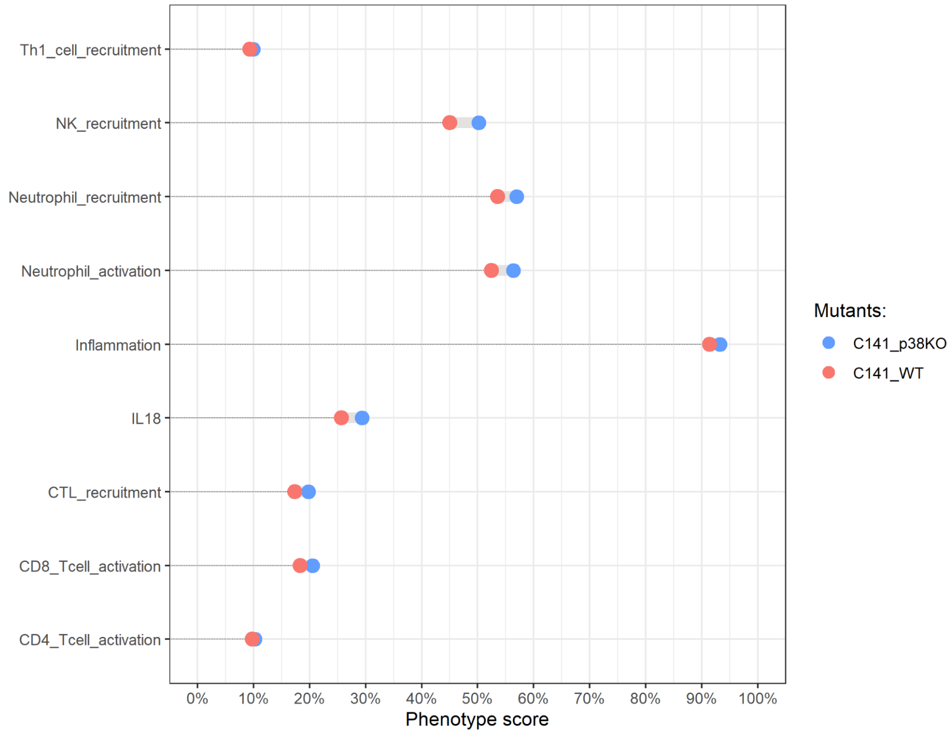


***Supplementary Figure 11:*** Close-up comparison of the effect of p38 knockout on the phenotype scores of the macrophages model personalised for patient C141. Phenotype scores of the recruitment of immune cells were gathered for the wild-type C141 (red) and the p38 knockout C141 MaBoSS simulation (blue).

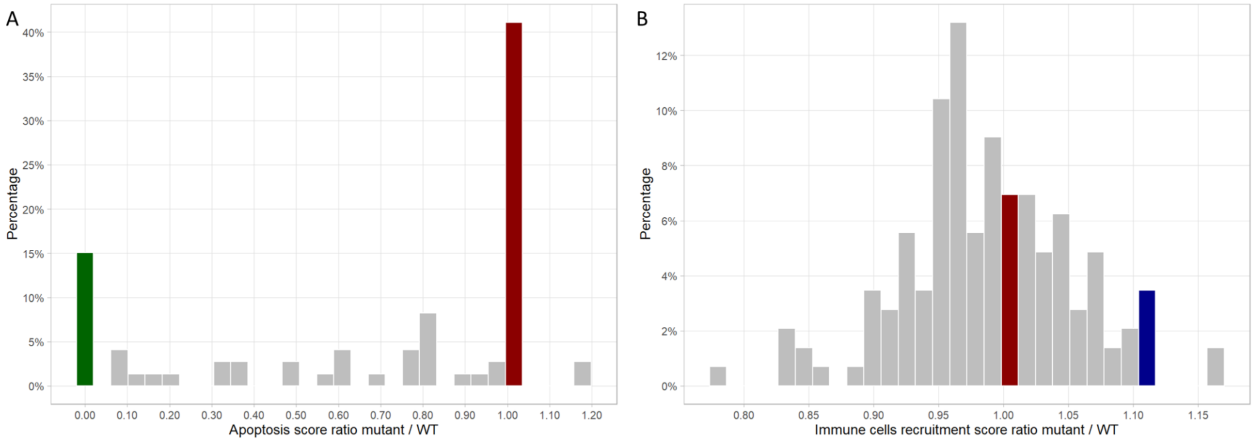


***Supplementary Figure 12:*** High-throughput study of all the KO mutants of the Boolean models considered in the multiscale simulations. The resulting scores for the relevant phenotypes apoptosis of epithelial cells (A) and the recruitment of immune cells by macrophages (B) were compared to the wild-type values (red bar). The green bar represents the score of FADD knockout, and the blue bar represents the score of p38 knockout.

***Supplementary Table 5: provided as a separate excel file***

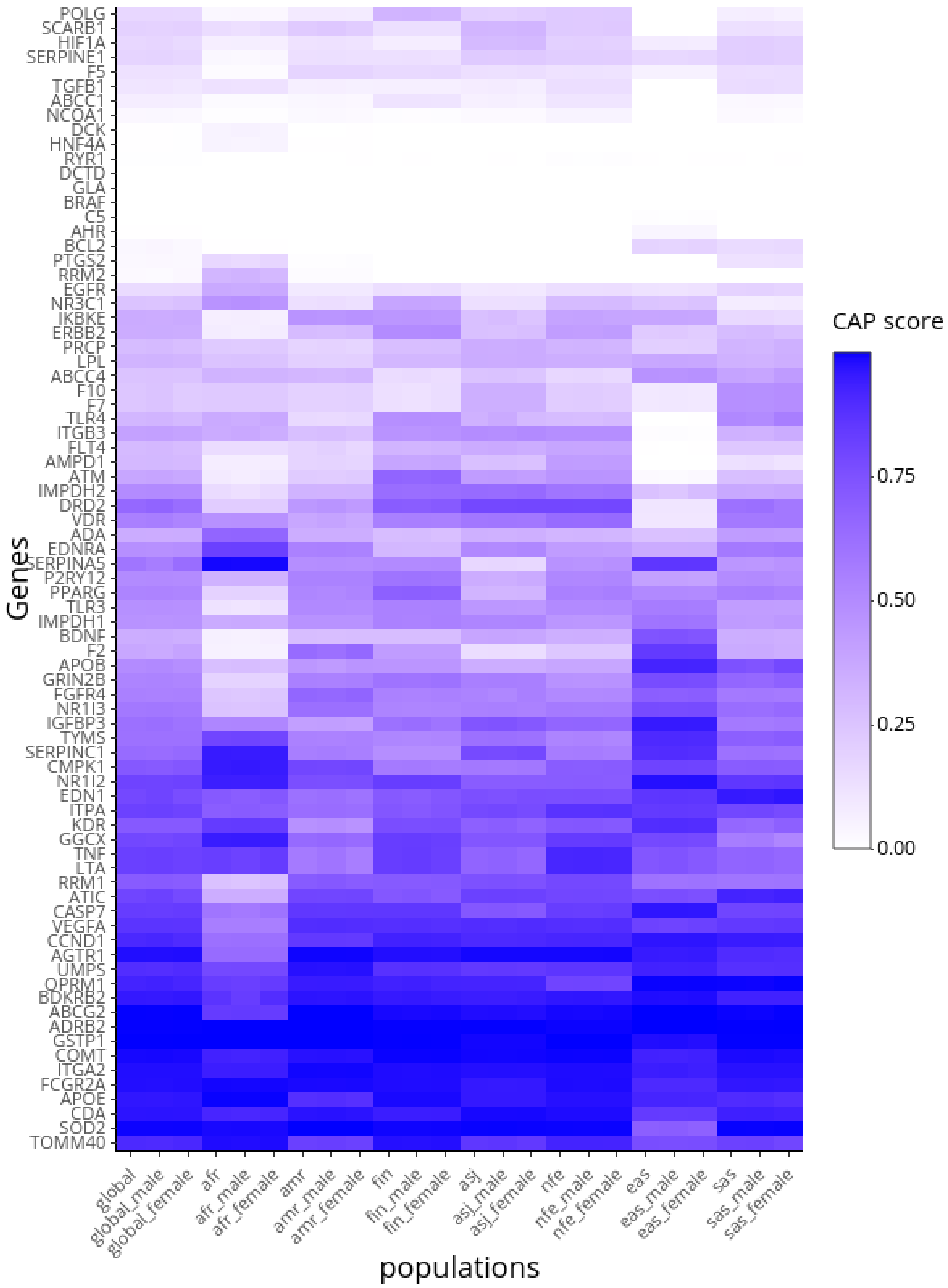


***Supplementary Figure 13***: CAP score across different populations and stratified by sex. The heatmap displays the CAP score for each gene of the 79 genes computed for each different population and sex. The populations tested are afr: African/African American, eas: East Asian, asj: Ashkenazi Jewish, nfe: non-Finnish European, fin: Finnish European, sas: South Asian, amr: Latino/Admixed American.

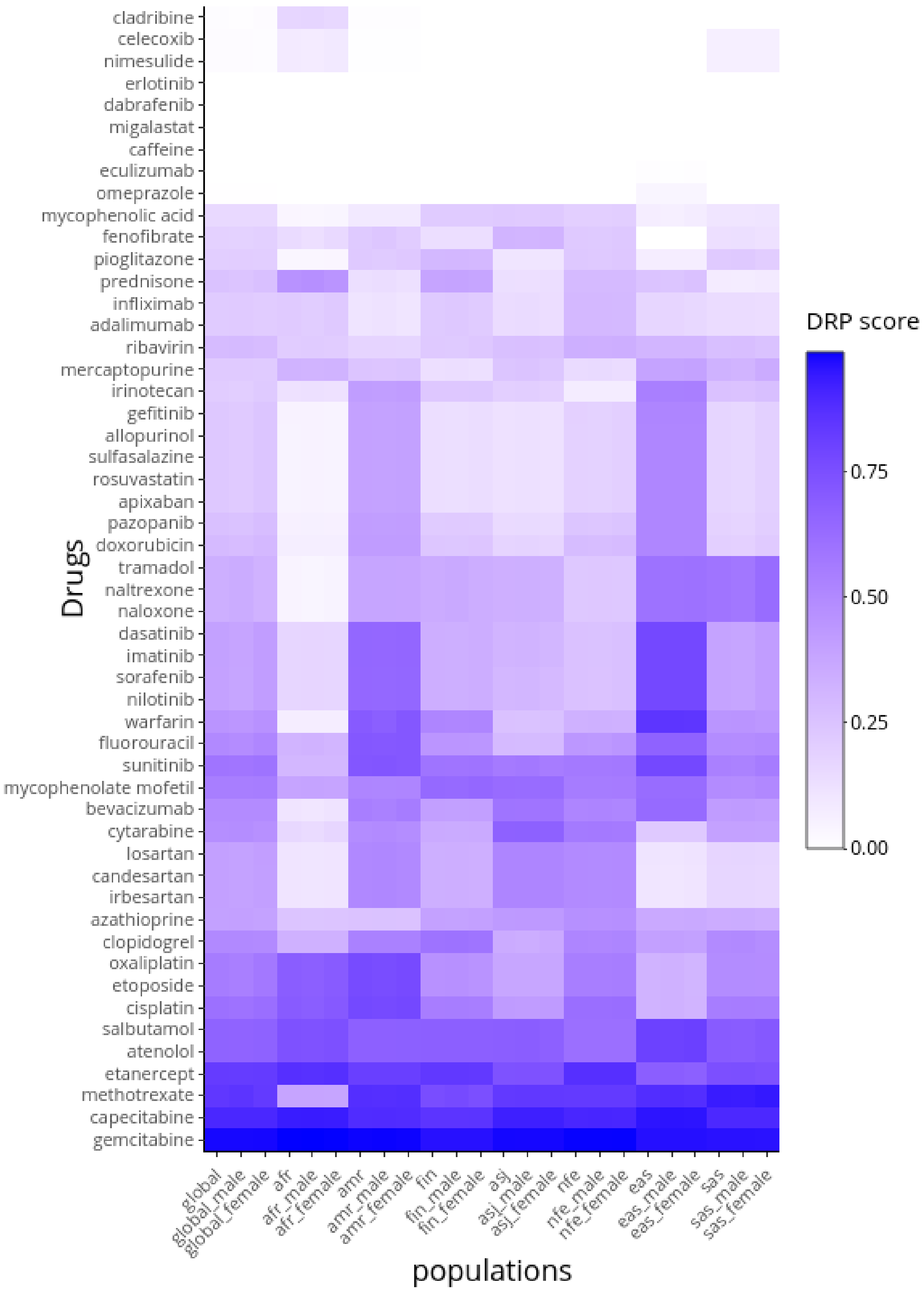


***Supplementary Figure 14***. DRP score for the set of 52 drugs was computed for each population and sex. The populations tested are afr: African/African American, eas: East Asian, asj: Ashkenazi Jewish, nfe: non-Finnish European, fin: Finnish European, sas: South Asian, amr: Latino/Admixed American.
https://docs.google.com/spreadsheets/d/1NAO3x72pZjWgACns44pJWCe9O8_8Wdamg7xMhxC3Atw/edit#gid=0

***Supplementary Table 6:*** Top ranking molecules across all occurrences.

| Ranking | Aggregated centrality value | Map species |
|---|---|---|
| 1 | 3.088928834 | UNIPROT:P0DTD1<br>**Replicase polyprotein 1ab SARS COV2** |
| 2 | 3.060313914 | UNIPROT:P0DTC9<br>**Nucleoprotein NCAP_SARS2** |
| 3 | 3.045092464 | UNIPROT:Q9BYF1<br>**Angiotensin-converting enzyme 2** |
| 4 | 2.974406697 | UNIPROT:P0DTC3<br>**ORF3a protein SARS COV2** |
| 5 | 2.960697776 | UNIPROT:P59632<br>**ORF3a protein SARS COV** |
| 6 | 2.926930595 | UNIPROT:P59637<br>**Envelope small membrane protein SARS COV** |
| 7 | 2.921492419 | UNIPROT:P59595<br>**Nucleoprotein SARS COV** |
| 8 | 2.91891734 | **Nsp6** |
| 9 | 2.917060121 | UNIPROT:P0DTC4<br>**Envelope small membrane protein SARS COV2** |
| 10 | 2.894981602 | UNIPROT:P0DTC1<br>**Replicase polyprotein 1a SARS COV2** |

***Supplementary Table 7.*** Top ranking species regarding the aggregated centrality values within the Interferon type I pathway.

| Aggregated centrality value | Map species |
|---|---|
| 2.99852173 | UNIPROT:Q14653<br>**IRF3** |
| 2.90197689 | **Nsp3** |
| 2.833324697 | UNIPROT:Q04206;UNIPROT:P19838 |

| | |
|---|---|
| | **RELA:NFKB1** |
| 2.770348427 | **Nsp15** |
| 2.673571447 | UNIPROT:Q7Z434<br>**MAVS** |
| 2.568232753 | **Viral_dsRNA** |
| 2.522809862 | UNIPROT:P01574<br>**IFNB** |
| 2.425625918 | **Orf6** |
| 2.393485439 | UNIPROT:Q14164;UNIPROT:Q9UHD2;UNIPROT:Q13114;UNIPROT:Q92844<br>**IKBKE:TBK1:TRAF3:TANK** |
| 2.341295307 | UNIPROT:O15455<br>**TLR3** |

***Supplementary Table 8.*** Top ranking species regarding the aggregated centrality values within the Interferon lambda pathway.

| **Aggregated centrality value** | **Map species** |
|---|---|
| 2.647094921 | UNIPROT:Q14653<br>**IRF3** |
| 2.614378989 | **IFN_III** |
| 2.324754461 | UNIPROT:P10914<br>**IRF1** |
| 2.313925511 | UNIPROT:Q8IZI9;UNIPROT:Q8IZJ0;UNIPROT:Q8IU54;UNIPROT:K9M1U5<br>**IFNL3: IFNL2:IFNL1:IFNL4** |
| 2.122800448 | UNIPROT:Q8IU57;UNIPROT:P23458;UNIPROT:P29597<br>**IFNLR1:JAK1:TYK2** |
| 2.027010129 | **dsRNA** |
| 1.923644106 | UNIPROT:O95786<br>**RIG-I** |
| 1.814816851 | UNIPROT:O14920;UNIPROT:O15111;UNIPROT:O95786;UNIPROT:Q9UHD2;UNIPROT:Q7Z434<br>**IKK:TBK1:RIG-I:MAVS** |

| | |
|---|---|
| 1.790330986 | UNIPROT:O95786;UNIPROT:Q7Z434<br>**RIG-I:MAVS** |
| 1.537042334 | UNIPROT:Q7Z434;UNIPROT:O95786<br>**MAVS:RIG-I** |

***Supplementary Table 9.*** Top ranking species regarding the aggregated centrality values within the Apoptosis pathway.

| **Aggregated centrality value** | **Map species** |
|---|---|
| 2.790670304 | UNIPROT:Q14790<br>**CASP8** |
| 2.575114314 | UNIPROT:P55211<br>**CASP9** |
| 2.46327704 | **Apoptosis** |
| 2.33125067 | UNIPROT:P59637<br>**Envelope small membrane protein SARS** |
| 2.208875087 | UNIPROT:P42574<br>**CASP3** |
| 2.208875087 | UNIPROT:P55210<br>**CASP7** |
| 2.179943105 | UNIPROT:Q13158<br>**FADD** |
| 2.146130422 | UNIPROT:P31749<br>**AKT1** |
| 1.796281512 | UNIPROT:P99999<br>**CYCS** |
| 1.775255449 | UNIPROT:P55957<br>**BID** |

***Supplementary Table 10.*** Top ranking species regarding the aggregated centrality values within the Coagulation pathway.

| **Aggregated centrality value** | **Map species** |
|---|---|
| 3.024512997 | **SARS_CoV_2_infection** |
| 2.768981505 | UNIPROT:P00734<br>**Prothrombin/ Thrombin (F2)** |

| | |
|---|---|
| 2.742875284 | UNIPROT:P00747<br>**Plasminogen (PLG)** |
| 2.664962377 | UNIPROT:P13671;UNIPROT:P07357;UNIPROT:P01031;UNIPROT:P10643;UNIPROT:P07360;UNIPROT:P02748;UNIPROT:P07358<br>**C6:C8A:C5:C7:C8G:C9:C8B (C5b-9)** |
| 2.65809911 | UNIPROT:P30556<br>**AGTR1** |
| 2.657606153 | UNIPROT:P0C0L5<br>**C4B** |
| 2.631358595 | **Thrombosis** |
| 2.549145423 | UNIPROT:P00750<br>**PLAT** |
| 2.540683322 | UNIPROT:P04070<br>**PROC** |
| 2.432797266 | UNIPROT:P05121<br>**SERPINE1** |

***Supplementary Table 11.*** Top ranking species regarding the aggregated centrality values within the Renin-Angiotensin pathway.

| **Aggregated centrality value** | **Map species** |
|---|---|
| 2.952636737 | UNIPROT:P30556<br>**AGTR1** |
| 2.876342802 | UNIPROT:Q9BYF1<br>**ACE2** |
| 2.760902828 | UNIPROT:P12821<br>**ACE** |
| 2.56692845 | UNIPROT:P04201<br>**MAS** |
| 2.382326458 | UNIPROT:P50052<br>**AGTR2** |
| 2.296031373 | UNIPROT:P0DTC2;UNIPROT:P59594<br>**SPIKE_SARS2: SPIKE_SARS** |
| 2.237870739 | **angiotensin_A** |

| | |
|---|---|
| 2.217315056 | **angiotensin_1-12** |
| 2.141851032 | **angiotensin_3-7** |
| 2.135741406 | UNIPROT:P08473<br>**EPN** |

***Supplementary Table 12: 18 out of the 54 identified targets that rank in the top 1000 occurrences of the aggregated graph***

| ***Protein / Gene Complex*** | ***Node label*** | ***Rank*** | ***Aggregated centrality value*** |
|---|---|---|---|
| *ATF4* | *UNIPROT:P18848* | *385* | *1.901239738* |
| *ATF6* | *UNIPROT:P18850* | *129* | *2.33493298* |
| *TP53* | *UNIPROT:P04637* | *938* | *1.382271605* |
| *STAT2* | *UNIPROT:P52630* | *92* | *2.4594734* |
| *BACH1* | *UNIPROT:O14867* | *967* | *1.358971275* |
| *TCF12* | *UNIPROT:Q99081* | *512* | *1.84628427* |
| *OAS1* | *UNIPROT:P00973* | *550* | *1.805509535* |
| *OAS2* | *UNIPROT:P29728* | *549* | *1.805765354* |
| *OAS3* | *UNIPROT:Q9Y6K5* | *548* | *1.805765354* |
| *TLR9* | *UNIPROT:Q9NR96* | *636* | *1.713899544* |
| *TREML4* | *UNIPROT:Q6UXN2* | *145* | *2.287025114* |
| *ISG15* | *UNIPROT:P05161* | *561* | *1.787584161* |
| *LNPEP* | *UNIPROT:Q9UIQ6* | *893* | *1.426754601* |
| *RBX1* | *UNIPROT:P62877* | *98* | *2.437577365* |
| *HMOX1* | *UNIPROT:P09601* | *32* | *2.743196317* |
| *ALG5* | *Hops_space_Complex UNIPROT:J9TC74;UNIPROT:Q96S66;UNIPROT:Q9UH99;UNIPROT:Q8N6S5;UNIPROT:P09601;UNIPROT:Q96JC1;UNIPROT:Q9H270;UNIPROT:Q9Y673* | *678* | *1.670172933* |
| *ARL6IP6* | *Hops_space_Complex UNIPROT:J9TC74;UNIPROT:Q96S66;UNIPROT:Q9UH99;UNIPROT:Q8N6S5;UNIPROT:P09601;UNIPROT:Q96JC1;UNIPROT:Q9H270;UNIPROT:Q9Y673* | *678* | *1.670172933* |
| *CASP7* | *UNIPROT:P55210* | *698* | *1.649386366* |

***Supplementary Table 13: 11 out of the 54 identified targets that rank in the top 30% occurrences of the aggregated graph***

| ***Protein / Gene*** | ***Node label*** | ***Rank*** | ***Aggregated centrality value*** |
|---|---|---|---|
| *HMOX1* | *UNIPROT09601* | *32* | *2,74319632* |
| *STAT2* | *UNIPROT52630* | *92* | *2,4594734* |

| | | | |
|---|---|---|---|
| *RBX1* | *UNIPROT62877* | *98* | *2,43757737* |
| *ATF6* | *UNIPROT18850* | *129* | *2,33493298* |
| *TREML4* | *UNIPROT:Q6UXN2* | *145* | *2,28702511* |
| *ATF4* | *UNIPROT18848* | *385* | *1,90123974* |
| *TCF12* | *UNIPROT:Q99081* | *512* | *1,84628427* |
| *OAS3* | *UNIPROT:Q9Y6K5* | *548* | *1,80576535* |
| *OAS2* | *UNIPROT29728* | *549* | *1,80576535* |
| *OAS1* | *UNIPROT00973* | *550* | *1,80550953* |
| *ISG15* | *UNIPROT05161* | *561* | *1,78758416* |

***Supplementary Table 14***: Number of drug candidates targeting transcription factors detected; "Common Transcription Factor" represents transcription factors detected for both NHBE and A549 cell types listed in Table 1; "External Clinical Trials" represents drugs already in external clinical trials for COVID19 designated in DrugBank.

| | External Clinical Trials | Not External Clinical Trials |
|---|---|---|
| Common Transcription Factor | 47 | 160 |
| Not Common Transcription Factor | 36 | 103 |

```
Cypher query example: process phosphorylation
MATCH path1 = (product_sv:StateVariable) <- [:HAS_STATE_VARIABLE] -
(product:Macromolecule) <- [:PRODUCES] - (process:Process) -
[:CONSUMES] -> (reactant:Macromolecule) - [:HAS_STATE_VARIABLE] ->
(reactant_sv:StateVariable)
WHERE reactant.label = product.label AND reactant_sv.value IS NULL AND
product_sv.value = "P" AND (product_sv.variable IS NULL AND
reactant_sv.variable IS NULL AND product_sv.order = reactant_sv.order OR
product_sv.variable = reactant_sv.variable)
OPTIONAL MATCH path2 = (catalyzer:Macromolecule) - [:CATALYZES] ->
(process)
OPTIONAL MATCH path3 = (catalyzer) - [:HAS_STATE_VARIABLE] ->
(catalyzer_sv:StateVariable)
RETURN path1, path2, path3
```

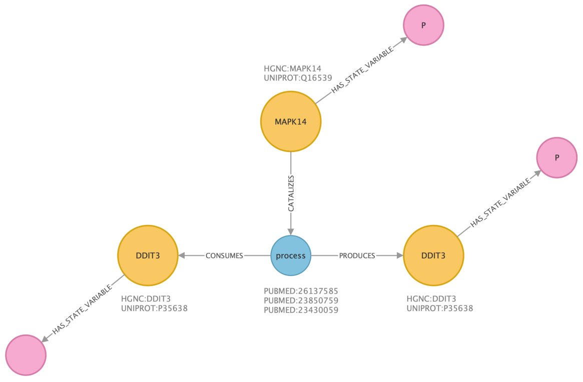


***Supplementary Figure 15.* C19DM-Neo4j data model: an example of a protein phosphorylation process.** For illustration purposes, we show a single process from the Endoplasmic Reticulum Stress map: DDIT3 phosphorylation activated by MAPK14. Nodes represent macromolecules (yellow), processes (blue) and state variables (red). Edges represent relationships: CONSUMES (reactant-process relationship), PRODUCES (process-product), CATALYZES (process-catalyzer),

HAS_STATE_VARIABLE (macromolecule-state_variable). Properties of nodes are shown as text near nodes. Entries of the HGNC database and PubMed IDs for supporting references. The C19DM-Neo4j database and the Cypher query language offer flexible querying of the COVID-19 Disease Map resource. More examples and tutorials are available at https://fairdomhub.org/projects/219.
