## Supplementary Table 5 for "A versatile and interoperable computational framework for the analysis and modeling of COVID-19 disease mechanisms"

| Protein / gene | INDRA drugs | Clinical trials/ TF analysis | AILANI |
|---|---|---|---|
| **MAPK11** | MAPK11sb 203580chebi:CHEBI:90705INDRA (drugs)<br>MAPK11doramapimodchebi:CHEBI:40953INDRA (drugs)<br>MAPK11vx-745chebi:CHEBI:90528INDRA (drugs)<br>MAPK11sb-202190chebi:CHEBI:79090INDRA (drugs)<br>MAPK11[11c]-sorafenibchembl.compound:CHEMBL1760433INDRA (drugs)<br>MAPK11pcid:5353940pubchem.compound:5353940INDRA (drugs)<br>MAPK11nitratechebi:CHEBI:17632INDRA (drugs)<br>MAPK11(+)-abscisic acidchebi:CHEBI:2365INDRA (drugs)<br>MAPK11reactive oxygen specieschebi:CHEBI:26523INDRA (drugs) | | MAPK11mirna:hsa-miR-122-5pAILANI<br>MAPK11mirna:hsa-miR-124-3pAILANI<br>MAPK11mirna:hsa-let-7a-5pAILANI |
| **SMAD1** | SMAD1imatinib methanesulfonatechebi:CHEBI:31690INDRA (drugs)<br>SMAD1dorsomorphinchebi:CHEBI:78510INDRA (drugs)<br>SMAD1ethanolchebi:CHEBI:16236INDRA (drugs)<br>SMAD1lipopolysaccharidechebi:CHEBI:16412INDRA (drugs)<br>SMAD1melatoninchebi:CHEBI:16796INDRA (drugs) | | SMAD1mirna:hsa-miR-155-5pAILANI<br>SMAD1mirna:hsa-miR-26a-5pAILANI<br>SMAD1mirna:hsa-miR-30a-5pAILANI<br>SMAD1mirna:hsa-miR-30b-5pAILANI<br>SMAD1mirna:hsa-miR-30c-5pAILANI<br>SMAD1mirna:hsa-miR-30d-5pAILANI<br>SMAD1mirna:hsa-miR-16-5pAILANI<br>SMAD1mirna:hsa-miR-205-5pAILANI<br>SMAD1mirna:hsa-miR-675-3pAILANI<br>SMAD1mirna:hsa-miR-3117-3pAILANI<br>SMAD1mirna:hsa-miR-216b-5pAILANI<br>SMAD1mirna:hsa-miR-3591-3pAILANI<br>SMAD1mirna:hsa-miR-21-3pAILANI<br>SMAD1mirna:hsa-miR-186-5pAILANI |
| **TICAM1** | TICAM1chebi:CHEBI:15756hexadecanoic acidINDRA (drugs)<br>TICAM1chebi:CHEBI:16412lipopolysaccharideINDRA (drugs)<br>TICAM1chebi:CHEBI:27881resveratrolINDRA (drugs)<br>TICAM1chebi:CHEBI:33351seaborgium atomINDRA (drugs)<br>TICAM1chebi:CHEBI:3962curcuminINDRA (drugs) | | TICAM1mirna:hsa-miR-193b-3pAILANI<br>TICAM1mirna:hsa-miR-221-3pAILANI |
| **TBK1** | TBK1chebi:CHEBI:113532thymoquinoneINDRA (drugs)<br>TBK1chebi:CHEBI:124930n-[3-[[5-cyclopropyl-2-[3-(4-morpholinylmethyl)anilino]-4-pyrimidinyl]amino]propyl]cyclobutanecarboxamideINDRA (drugs)<br>TBK1chebi:CHEBI:16113cholesterolINDRA (drugs)<br>TBK1chebi:CHEBI:17234glucoseINDRA (drugs)<br>TBK1chebi:CHEBI:24261glucocorticoidINDRA (drugs)<br>TBK1chebi:CHEBI:27881resveratrolINDRA (drugs)<br>TBK1chebi:CHEBI:31205amlexanoxINDRA (drugs)<br>TBK1chebi:CHEBI:38940sunitinibINDRA (drugs)<br>TBK1chebi:CHEBI:91439n-[3-[[5-iodo-4-[3-[[oxo(thiophen-2-yl)methyl]amino]propylamino]-2-pyrimidinyl]amino]phenyl]-1-pyrrolidinecarboxamideINDRA (drugs)<br>TBK1chembl.compound:CHEMBL1567sunitinib malateINDRA (drugs) | | TBK1mirna:hsa-let-7f-2-3pAILANI<br>TBK1mirna:hsa-miR-1185-1-3pAILANI<br>TBK1mirna:hsa-miR-1185-2-3pAILANI<br>TBK1mirna:hsa-miR-186-5pAILANI<br>TBK1mirna:hsa-miR-200b-3pAILANI<br>TBK1mirna:hsa-miR-200c-3pAILANI<br>TBK1mirna:hsa-miR-219a-2-3pAILANI<br>TBK1mirna:hsa-miR-219b-3pAILANI<br>TBK1mirna:hsa-miR-221-3pAILANI<br>TBK1mirna:hsa-miR-3133AILANI<br>TBK1mirna:hsa-miR-3671AILANI<br>TBK1mirna:hsa-miR-3674AILANI<br>TBK1mirna:hsa-miR-429AILANI<br>TBK1mirna:hsa-miR-452-5pAILANI<br>TBK1mirna:hsa-miR-4680-3pAILANI<br>TBK1mirna:hsa-miR-4724-5pAILANI<br>TBK1mirna:hsa-miR-6792-5pAILANI |
| **IKBKE** | IKBKE chebi:CHEBI:124930 n-[3-[[5-cyclopropyl-2-[3-(4-morpholinylmethyl)anilino]-4-pyrimidinyl]amino]propyl]cyclobutanecarboxamide INDRA (drugs)<br>IKBKE chebi:CHEBI:31205 amlexanox INDRA (drugs)<br>IKBKE chebi:CHEBI:32692 4-(n-nitrosomethylamino)-1-(3-pyridyl)butan-1-one INDRA (drugs)<br>IKBKE chebi:CHEBI:49468 silver(1+) INDRA (drugs)<br>IKBKE chebi:CHEBI:49786 nickel(2+) INDRA (drugs)<br>IKBKE chebi:CHEBI:73888 gly-asn INDRA (drugs)<br>IKBKE chebi:CHEBI:91439 n-[3-[[5-iodo-4-[3-[[oxo(thiophen-2-yl)methyl]amino]propylamino]-2-pyrimidinyl]amino]phenyl]-1-pyrrolidinecarboxamide INDRA (drugs)<br>IKBKE chembl.compound:CHEMBL1567 sunitinib malate INDRA (drugs)<br>IKBKE pubchem.compound:3086686 sunitinib INDRA (drugs) | | IKBKE mirna:hsa-miR-124-3p AILANI<br>IKBKE mirna:hsa-miR-155-5p AILANI<br>IKBKE mirna:hsa-miR-296-5p AILANI |
| **IRF3** | IRF3 chebi:CHEBI:16412 lipopolysaccharide INDRA (drugs)<br>IRF3 chebi:CHEBI:3962 curcumin INDRA (drugs)<br>IRF3 chebi:CHEBI:67208 double-stranded rna INDRA (drugs)<br>IRF3 pubchem.compound:86583374 poly i:c, polyinosinic-polycytidylic acid INDRA (drugs) | IRF3 chloramphenicol drugbank:DB00446 TF analysis<br>IRF3 zinc drugbank:DB01593 TF analysis<br>IRF3 threonine drugbank:DB00156 TF analysis<br>IRF3 tyrosine drugbank:DB00135 TF analysis | |

| Protein / gene | INDRA drugs | Clinical trials/ TF analysis | AILANI |
|---|---|---|---|
| **ATF4** | ATF4 sorafenib chebi:CHEBI:50924 INDRA (drugs)<br>ATF4 lipopolysaccharide chebi:CHEBI:16412 INDRA (drugs)<br>ATF4 glucose chebi:CHEBI:17234 INDRA (drugs)<br>ATF4 tomatidine chebi:CHEBI:9629 INDRA (drugs)<br>ATF4 ursolic acid chebi:CHEBI:9908 INDRA (drugs)<br>ATF4 sirolimus chebi:CHEBI:9168 INDRA (drugs) | | ATF4 pseudoephedrine drugbank:DB00852 AILANI<br>ATF4 mirna:hsa-miR-214-3p AILANI<br>ATF4 mirna:hsa-miR-196b-5p AILANI<br>ATF4 mirna:hsa-miR-342-3p AILANI |
| **ATF6** | ATF61,10-phenanthrolinechebi:CHEBI:44975INDRA (drugs)<br>ATF6arg-leuchebi:CHEBI:73815INDRA (drugs)<br>ATF6melatoninchebi:CHEBI:16796INDRA (drugs)<br>ATF6ethanolchebi:CHEBI:16236INDRA (drugs)<br>ATF6fluoridechebi:CHEBI:17051INDRA (drugs)<br>ATF6glucosechebi:CHEBI:17234INDRA (drugs) | | ATF6pseudoephedrinedrugbank:DB00852AILANI<br>ATF6mirna:hsa-miR-503-5pAILANI<br>ATF6mirna:hsa-miR-424-5pAILANI<br>ATF6mirna:hsa-miR-335-5pAILANI<br>ATF6mirna:hsa-miR-191-5pAILANI<br>ATF6mirna:hsa-miR-199a-5pAILANI<br>ATF6mirna:hsa-miR-552-5pAILANI<br>ATF6mirna:hsa-miR-6780a-5pAILANI<br>ATF6mirna:hsa-miR-6779-5pAILANI<br>ATF6mirna:hsa-miR-3689cAILANI<br>ATF6mirna:hsa-miR-3689b-3pAILANI<br>ATF6mirna:hsa-miR-3689a-3pAILANI<br>ATF6mirna:hsa-miR-30b-3pAILANI<br>ATF6mirna:hsa-miR-1273h-5pAILANI<br>ATF6mirna:hsa-miR-6799-5pAILANI<br>ATF6mirna:hsa-miR-6883-5pAILANI<br>ATF6mirna:hsa-miR-6785-5pAILANI<br>ATF6mirna:hsa-miR-4728-5pAILANI<br>ATF6mirna:hsa-miR-149-3pAILANI<br>ATF6mirna:hsa-miR-7106-5pAILANI<br>ATF6mirna:hsa-miR-6871-5pAILANI<br>ATF6mirna:hsa-miR-208b-5pAILANI<br>ATF6mirna:hsa-miR-208a-5pAILANI<br>ATF6mirna:hsa-miR-7153-3pAILANI |
| **MBTPS1** | MBTPS13,4-dichloroisocoumarinmesh:C046235INDRA (drugs)<br>MBTPS1antagonistchebi:CHEBI:48706INDRA (drugs)<br>MBTPS1fingolimod hydrochloridechebi:CHEBI:63112INDRA (drugs)<br>MBTPS1sb 203580chebi:CHEBI:90705INDRA (drugs)<br>MBTPS1sb202190pubchem.compound:5353940INDRA (drugs)<br>MBTPS1acetylsalicylic acidchebi:CHEBI:15365INDRA (drugs) | | MBTPS1mirna:hsa-miR-769-3pAILANI<br>MBTPS1mirna:hsa-miR-193b-3pAILANI<br>MBTPS1mirna:hsa-let-7c-5pAILANI |

| Protein / gene | INDRA drugs | Clinical trials/ TF analysis | AILANI |
| --- | --- | --- | --- |
| **TP53** | TP53resveratrolchebi:CHEBI:27881INDRA (drugs)<br>TP53doxorubicinchebi:CHEBI:28748INDRA (drugs)<br>TP53sirolimuschebi:CHEBI:9168INDRA (drugs)<br>TP53cadmium atomchebi:CHEBI:22977INDRA (drugs)<br>TP53trichostatin achebi:CHEBI:46024INDRA (drugs)<br>TP53o-(3-o-d-galactosyl-n-acetyl-beta-d-galactosaminyl)-l-serinechebi:CHEBI:16981 INDRA (drugs)<br>TP53nutlin-3chebi:CHEBI:46742INDRA (drugs)<br>TP53glucosechebi:CHEBI:17234INDRA (drugs)<br>TP53cisplatinchebi:CHEBI:27899INDRA (drugs)<br>TP53nacchebi:CHEBI:7421INDRA (drugs) | TP53 amifostine drugbank:DB01143 TF analysis<br>TP53 aspirin drugbank:DB00945 TF analysis<br>TP53 bleomycin drugbank:DB00290 TF analysis<br>TP53 bortezomib drugbank:DB00188 TF analysis<br>TP53 caffeine drugbank:DB00201 TF analysis<br>TP53 carboplatin drugbank:DB00958 TF analysis<br>TP53 chlorambucil drugbank:DB00291 TF analysis<br>TP53 cisplatin drugbank:DB00515 TF analysis<br>TP53 curcumin drugbank:DB11672 TF analysis<br>TP53 cyclophosphamide drugbank:DB00531 TF analysis<br>TP53 cytarabine drugbank:DB00987 TF analysis<br>TP53 decitabine drugbank:DB01262 TF analysis<br>TP53 dicumarol drugbank:DB00266 TF analysis<br>TP53 docetaxel drugbank:DB01248 TF analysis<br>TP53 doxorubicin drugbank:DB00997 TF analysis<br>TP53 epirubicin drugbank:DB00445 TF analysis<br>TP53 etoposide drugbank:DB00773 TF analysis<br>TP53 fludarabine drugbank:DB01073 TF analysis<br>TP53 fluorouracil drugbank:DB00544 TF analysis<br>TP53 gemcitabine drugbank:DB00441 TF analysis<br>TP53 hydroxyurea drugbank:DB01005 TF analysis<br>TP53 irinotecan drugbank:DB00762 TF analysis<br>TP53 lomustine drugbank:DB01206 TF analysis<br>TP53 melphalan drugbank:DB01042 TF analysis<br>TP53 methylprednisolone drugbank:DB00959 TF analysis<br>TP53 methylprednisolone hemisuccinate drugbank:DB14644 TF analysis<br>TP53 mitomycin drugbank:DB00305 TF analysis<br>TP53 oxaliplatin drugbank:DB00526 TF analysis<br>TP53 paclitaxel drugbank:DB01229 TF analysis<br>TP53 parthenolide drugbank:DB13063 TF analysis<br>TP53 prednisolone drugbank:DB00860 TF analysis<br>TP53 prednisolone acetate drugbank:DB15566 TF analysis<br>TP53 prednisolone phosphate drugbank:DB14631 TF analysis<br>TP53 prednisone drugbank:DB00635 TF analysis<br>TP53 progesterone drugbank:DB00396 TF analysis<br>TP53 rituximab drugbank:DB00073 TF analysis<br>TP53 sulindac drugbank:DB00605 TF analysis<br>TP53 tamoxifen drugbank:DB00675 TF analysis<br>TP53 temozolomide drugbank:DB00853 TF analysis<br>TP53 topotecan drugbank:DB01030 TF analysis<br>TP53 venetoclax drugbank:DB11581 TF analysis<br>TP53 vinblastine drugbank:DB00570 TF analysis<br>TP53 vincristine drugbank:DB00541 TF analysis<br>TP53 zinc drugbank:DB01593 TF analysis<br>TP53 zinc acetate drugbank:DB14487 TF analysis<br>TP53 zinc chloride drugbank:DB14533 TF analysis<br>TP53 zinc sulfate, unspecified form drugbank:DB14548 TF analysis | TP531-(9-ethyl-9h-carbazol-3-yl)-n-methylmethanaminedrugbank: DB08363AILANI TP53mirna:hsa-miR-125b-5pAILANI<br>TP53mirna:hsa-miR-125a-5pAILANI<br>TP53mirna:hsa-miR-25-3pAILANI<br>TP53mirna:hsa-miR-30d-5pAILANI<br>TP53mirna:hsa-miR-1285-3pAILANI<br>TP53mirna:hsa-miR-612AILANI<br>TP53mirna:hsa-miR-15a-5pAILANI<br>TP53mirna:hsa-miR-16-5pAILANI<br>TP53mirna:hsa-miR-221-3pAILANI<br>TP53mirna:hsa-miR-222-3pAILANI<br>TP53mirna:hsa-miR-214-3pAILANI<br>TP53mirna:hsa-miR-10b-5pAILANI<br>TP53mirna:hsa-miR-34a-5pAILANI<br>TP53mirna:hsa-miR-608AILANI<br>TP53mirna:hsa-miR-605-5pAILANI<br>TP53mirna:hsa-miR-504-5pAILANI<br>TP53mirna:hsa-miR-485-5pAILANI<br>TP53mirna:hsa-miR-27a-3pAILANI<br>TP53mirna:hsa-miR-454-3pAILANI<br>TP53mirna:hsa-miR-324-5pAILANI<br>TP53mirna:hsa-miR-150-5pAILANI<br>TP53mirna:hsa-miR-92a-3pAILANI<br>TP53mirna:hsa-miR-375AILANI<br>TP53mirna:hsa-miR-200a-3pAILANI<br>TP53mirna:hsa-miR-491-5pAILANI<br>TP53mirna:hsa-miR-30a-5pAILANI<br>TP53mirna:hsa-miR-30b-5pAILANI<br>TP53mirna:hsa-miR-30c-5pAILANI<br>TP53mirna:hsa-miR-30e-5pAILANI<br>TP53mirna:hsa-miR-19b-3pAILANI<br>TP53mirna:hsa-miR-92a-2-5pAILANI<br>TP53mirna:hsa-miR-92a-1-5pAILANI<br>TP53mirna:hsa-miR-20a-5pAILANI<br>TP53mirna:hsa-miR-18a-5pAILANI<br>TP53mirna:hsa-miR-106b-5pAILANI<br>TP53mirna:hsa-miR-106a-5pAILANI<br>TP53mirna:hsa-miR-17-5pAILANI<br>TP53mirna:hsa-miR-6803-5pAILANI<br>TP53mirna:hsa-miR-6751-5pAILANI<br>TP53mirna:hsa-miR-7150AILANI<br>TP53mirna:hsa-miR-6778-5pAILANI<br>TP53mirna:hsa-miR-1233-5pAILANI<br>TP53mirna:hsa-miR-6766-5pAILANI<br>TP53mirna:hsa-miR-6756-5pAILANI<br>TP53mirna:hsa-miR-7110-5pAILANI<br>TP53mirna:hsa-miR-4651AILANI<br>TP53mirna:hsa-miR-6842-5pAILANI<br>TP53mirna:hsa-miR-6752-5pAILANI<br>TP53mirna:hsa-miR-6835-5pAILANI<br>TP53mirna:hsa-miR-6825-5pAILANI<br>TP53mirna:hsa-miR-6883-5pAILANI<br>TP53mirna:hsa-miR-6785-5pAILANI<br>TP53mirna:hsa-miR-4728-5pAILANI<br>TP53mirna:hsa-miR-149-3pAILANI<br>TP53mirna:hsa-miR-6731-5pAILANI<br>TP53mirna:hsa-miR-8085AILANI<br>TP53mirna:hsa-miR-4736AILANI<br>TP53mirna:hsa-miR-4763-3pAILANI<br>TP53mirna:hsa-miR-1207-5pAILANI<br>TP53mirna:hsa-miR-6797-5pAILANI<br>TP53mirna:hsa-miR-1249-5pAILANI<br>TP53mirna:hsa-miR-4644AILANI<br>TP53mirna:hsa-miR-4306AILANI<br>TP53mirna:hsa-miR-185-5pAILANI<br>TP53mirna:hsa-miR-6722-3pAILANI<br>TP53mirna:hsa-miR-1909-3pAILANI<br>TP53mirna:hsa-miR-6133AILANI<br>TP53mirna:hsa-miR-6130AILANI<br>TP53mirna:hsa-miR-6129AILANI<br>TP53mirna:hsa-miR-6127AILANI<br>TP53mirna:hsa-miR-4510AILANI<br>TP53mirna:hsa-miR-4419aAILANI<br>TP53mirna:hsa-miR-1273fAILANI<br>TP53mirna:hsa-miR-6760-5pAILANI<br>TP53mirna:hsa-miR-5197-5pAILANI<br>TP53mirna:hsa-miR-2110AILANI<br>TP53mirna:hsa-miR-4531AILANI |

| Protein / gene | INDRA drugs | Clinical trials/ TF analysis | AILANI |
|---|---|---|---|
| STAT1 | | STAT1 biotin drugbank:DB00121 TF analysis<br>STAT1 calcitriol drugbank:DB00136 TF analysis<br>STAT1 cholesterol drugbank:DB04540 TF analysis<br>STAT1 cisplatin drugbank:DB00515 TF analysis<br>STAT1 curcumin drugbank:DB11672 TF analysis<br>STAT1 cyclosporin a drugbank:DB00091 TF analysis<br>STAT1 cysteine drugbank:DB00151 TF analysis<br>STAT1 deferoxamine drugbank:DB00746 TF analysis<br>STAT1 dexamethasone drugbank:DB01234 TF analysis<br>STAT1 doxorubicin drugbank:DB00997 TF analysis<br>STAT1 ethanol drugbank:DB00898 TF analysis<br>STAT1 etoposide drugbank:DB00773 TF analysis<br>STAT1 fludarabine drugbank:DB01073 TF analysis<br>STAT1 imatinib drugbank:DB00619 TF analysis<br>STAT1 imiquimod drugbank:DB00724 TF analysis<br>STAT1 indomethacin drugbank:DB00328 TF analysis<br>STAT1 methimazole drugbank:DB00763 TF analysis<br>STAT1 niclosamide drugbank:DB06803 TF analysis<br>STAT1 nitric oxide drugbank:DB00435 TF analysis<br>STAT1 ornithine drugbank:DB00129 TF analysis<br>STAT1 oxygen drugbank:DB09140 TF analysis<br>STAT1 phenylalanine drugbank:DB00120 TF analysis<br>STAT1 prazosin drugbank:DB00457 TF analysis<br>STAT1 progesterone drugbank:DB00396 TF analysis<br>STAT1 ribavirin drugbank:DB00811 TF analysis<br>STAT1 rosiglitazone drugbank:DB00412 TF analysis<br>STAT1 threonine drugbank:DB00156 TF analysis<br>STAT1 tyrosine drugbank:DB00135 TF analysis<br>STAT1 valsartan drugbank:DB00177 TF analysis<br>STAT1 vitamin d drugbank:DB11094 TF analysis<br>STAT1 zinc drugbank:DB01593 TF analysis | |

| Protein / gene | INDRA drugs | Clinical trials/ TF analysis | AILANI |
|---|---|---|---|
| **FOS** | FOScocainechebi:CHEBI:27958INDRA (drugs)<br>FOS(-)-epigallocatechin 3-gallatechebi:CHEBI:4806INDRA (drugs)<br>FOSisotretinoinchebi:CHEBI:6067INDRA (drugs)<br>FOSn-acetyl-l-cysteinechebi:CHEBI:28939INDRA (drugs)<br>FOSmorphine(1+)chebi:CHEBI:58097INDRA (drugs)<br>FOSheparinchebi:CHEBI:28304INDRA (drugs)<br>FOSethanolchebi:CHEBI:16236INDRA (drugs)<br>FOScurcuminchebi:CHEBI:3962INDRA (drugs)<br>FOSheparinpubchem.compound:22833565INDRA (drugs) | FOS amphetamine drugbank:DB00182 TF analysis<br>FOS apomorphine drugbank:DB00714 TF analysis<br>FOS calcitriol drugbank:DB00136 TF analysis<br>FOS capsaicin drugbank:DB06774 TF analysis<br>FOS cetrorelix drugbank:DB00050 TF analysis<br>FOS chloramphenicol drugbank:DB00446 TF analysis<br>FOS clozapine drugbank:DB00363 TF analysis<br>FOS cocaine drugbank:DB00907 TF analysis<br>FOS curcumin drugbank:DB11672 TF analysis<br>FOS dexamethasone drugbank:DB01234 TF analysis<br>FOS dopamine drugbank:DB00988 TF analysis<br>FOS estradiol benzoate drugbank:DB13953 TF analysis<br>FOS haloperidol drugbank:DB00502 TF analysis<br>FOS inulin drugbank:DB00638 TF analysis<br>FOS levothyroxine drugbank:DB00451 TF analysis<br>FOS losartan drugbank:DB00678 TF analysis<br>FOS loxapine drugbank:DB00408 TF analysis<br>FOS manidipine drugbank:DB09238 TF analysis<br>FOS morphine drugbank:DB00295 TF analysis<br>FOS nadroparin drugbank:DB08813 TF analysis<br>FOS naloxone drugbank:DB01183 TF analysis<br>FOS olanzapine drugbank:DB00334 TF analysis<br>FOS ornithine drugbank:DB00129 TF analysis<br>FOS pilocarpine drugbank:DB01085 TF analysis<br>FOS pseudoephedrine drugbank:DB00852 TF analysis<br>FOS remoxipride drugbank:DB00409 TF analysis<br>FOS tamoxifen drugbank:DB00675 TF analysis<br>FOS threonine drugbank:DB00156 TF analysis<br>FOS troglitazone drugbank:DB00197 TF analysis<br>FOS tyrosine drugbank:DB00135 TF analysis | FOS pseudoephedrine drugbank:DB00852 AILANI<br>FOSmirna:hsa-miR-101-3pAILANI<br>FOSmirna:hsa-miR-221-3pAILANI<br>FOSmirna:hsa-miR-222-3pAILANI<br>FOSmirna:hsa-miR-181a-5pAILANI<br>FOSmirna:hsa-miR-139-5pAILANI<br>FOSmirna:hsa-miR-29b-3pAILANI<br>FOSmirna:hsa-miR-335-5pAILANI<br>FOSmirna:hsa-miR-215-5pAILANI<br>FOSmirna:hsa-miR-192-5pAILANI<br>FOSmirna:hsa-miR-490-5pAILANI<br>FOSmirna:hsa-miR-155-5pAILANI<br>FOSmirna:hsa-miR-181b-5pAILANI<br>FOSmirna:hsa-miR-770-5pAILANI<br>FOSmirna:hsa-miR-543AILANI<br>FOSmirna:hsa-miR-493-5pAILANI<br>FOSmirna:hsa-miR-338-3pAILANI<br>FOSmirna:hsa-miR-4530AILANI<br>FOSmirna:hsa-miR-323b-3pAILANI<br>FOSmirna:hsa-miR-4733-3pAILANI<br>FOSmirna:hsa-miR-3065-3pAILANI<br>FOSmirna:hsa-miR-5586-5pAILANI<br>FOSmirna:hsa-miR-29a-3pAILANI<br>FOSmirna:hsa-miR-29c-3pAILANI<br>FOSmirna:hsa-miR-6885-3pAILANI<br>FOSmirna:hsa-miR-8081AILANI<br>FOSmirna:hsa-miR-5089-3pAILANI<br>FOSmirna:hsa-miR-627-5pAILANI<br>FOSmirna:hsa-miR-937-5pAILANI<br>FOSmirna:hsa-miR-5581-3pAILANI<br>FOSmirna:hsa-miR-19b-2-5pAILANI<br>FOSmirna:hsa-miR-19b-1-5pAILANI<br>FOSmirna:hsa-miR-19a-5pAILANI<br>FOSmirna:hsa-miR-6077AILANI<br>FOSmirna:hsa-miR-548vAILANI<br>FOSmirna:hsa-miR-187-5pAILANI<br>FOSmirna:hsa-miR-6872-3pAILANI<br>FOSmirna:hsa-miR-6854-3pAILANI<br>FOSmirna:hsa-miR-8083AILANI<br>FOSmirna:hsa-miR-6816-3pAILANI<br>FOSmirna:hsa-miR-8080AILANI<br>FOSmirna:hsa-miR-619-5pAILANI<br>FOSmirna:hsa-miR-7151-3pAILANI<br>FOSmirna:hsa-miR-5095AILANI<br>FOSmirna:hsa-miR-6508-3pAILANI<br>FOSmirna:hsa-miR-4726-5pAILANI<br>FOSmirna:hsa-miR-4640-5pAILANI<br>FOSmirna:hsa-miR-3622b-5pAILANI<br>FOSmirna:hsa-miR-7107-5pAILANI<br>FOSmirna:hsa-miR-1234-3pAILANI<br>FOSmirna:hsa-miR-1292-3pAILANI<br>FOSmirna:hsa-miR-4438AILANI<br>FOSmirna:hsa-miR-6504-3pAILANI |

| Protein / gene | INDRA drugs | Clinical trials/ TF analysis | AILANI |
|---|---|---|---|
| **JUN** | JUNcurcuminchebi:CHEBI:3962INDRA (drugs)<br>JUNquercetinchebi:CHEBI:16243INDRA (drugs)<br>JUNminocyclinechebi:CHEBI:50694INDRA (drugs)<br>JUNacroleinchebi:CHEBI:15368INDRA (drugs)<br>JUNluteolinchebi:CHEBI:15864INDRA (drugs)<br>JUNanthra[1,9-cd]pyrazol-6(2h)-onechebi:CHEBI:90695INDRA (drugs)<br>JUN(-)-epigallocatechin 3-gallatechebi:CHEBI:4806INDRA (drugs)<br>JUN3',5'-cyclic ampchebi:CHEBI:17489INDRA (drugs)<br>JUNlipopolysaccharidechebi:CHEBI:16412INDRA (drugs)<br>JUNresveratrolchebi:CHEBI:27881INDRA (drugs) | JUN adapalene drugbank:DB00210 TF analysis<br>JUN arsenic trioxide drugbank:DB01169 TF analysis<br>JUN bexarotene drugbank:DB00307 TF analysis<br>JUN bortezomib drugbank:DB00188 TF analysis<br>JUN calcitriol drugbank:DB00136 TF analysis<br>JUN cerivastatin drugbank:DB00439 TF analysis<br>JUN chenodeoxycholic acid drugbank:DB06777 TF analysis<br>JUN chloramphenicol drugbank:DB00446 TF analysis<br>JUN curcumin drugbank:DB11672 TF analysis<br>JUN cyclosporin a drugbank:DB00091 TF analysis<br>JUN cysteine drugbank:DB00151 TF analysis<br>JUN cytarabine drugbank:DB00987 TF analysis<br>JUN dexamethasone drugbank:DB01234 TF analysis<br>JUN dicumarol drugbank:DB00266 TF analysis<br>JUN etoposide drugbank:DB00773 TF analysis<br>JUN irbesartan drugbank:DB01029 TF analysis<br>JUN mifepristone drugbank:DB00834 TF analysis<br>JUN oxygen drugbank:DB09140 TF analysis<br>JUN pioglitazone drugbank:DB01132 TF analysis<br>JUN plicamycin drugbank:DB06810 TF analysis<br>JUN pseudoephedrine drugbank:DB00852 TF analysis<br>JUN raloxifene drugbank:DB00481 TF analysis<br>JUN rosiglitazone drugbank:DB00412 TF analysis<br>JUN tamoxifen drugbank:DB00675 TF analysis<br>JUN threonine drugbank:DB00156 TF analysis<br>JUN troglitazone drugbank:DB00197 TF analysis<br>JUN tyrosine drugbank:DB00135 TF analysis<br>JUN vinblastine drugbank:DB00570 TF analysis<br>JUN vitamin d drugbank:DB11094 TF analysis | JUN vinblastine drugbank:DB00570 AILANI<br>JUN arsenic trioxide drugbank:DB01169 AILANI<br>JUN irbesartan drugbank:DB01029 AILANI<br>JUN pseudoephedrine drugbank:DB00852 AILANI<br>JUNmirna:hsa-miR-16-5pAILANI<br>JUNmirna:hsa-miR-15a-5pAILANI<br>JUNmirna:hsa-miR-30a-5pAILANI<br>JUNmirna:hsa-miR-155-5pAILANI<br>JUNmirna:hsa-miR-29c-3pAILANI<br>JUNmirna:hsa-miR-101-3pAILANI<br>JUNmirna:hsa-miR-26b-5pAILANI<br>JUNmirna:hsa-miR-342-3pAILANI<br>JUNmirna:hsa-miR-149-5pAILANI<br>JUNmirna:hsa-miR-93-5pAILANI<br>JUNmirna:hsa-miR-203a-3pAILANI<br>JUNmirna:hsa-miR-139-5pAILANI<br>JUNmirna:hsa-miR-149-3pAILANI<br>JUNmirna:hsa-miR-4728-5pAILANI<br>JUNmirna:hsa-miR-6785-5pAILANI<br>JUNmirna:hsa-miR-6883-5pAILANI<br>JUNmirna:hsa-miR-1321AILANI<br>JUNmirna:hsa-miR-4739AILANI<br>JUNmirna:hsa-miR-4756-5pAILANI<br>JUNmirna:hsa-miR-3691-3pAILANI<br>JUNmirna:hsa-miR-3178AILANI<br>JUNmirna:hsa-miR-4783-5pAILANI<br>JUNmirna:hsa-miR-4480AILANI<br>JUNmirna:hsa-miR-655-3pAILANI<br>JUNmirna:hsa-miR-374c-5pAILANI<br>JUNmirna:hsa-miR-4724-5pAILANI<br>JUNmirna:hsa-miR-8084AILANI<br>JUNmirna:hsa-miR-429AILANI<br>JUNmirna:hsa-miR-200b-3pAILANI<br>JUNmirna:hsa-miR-200c-3pAILANI<br>JUNmirna:hsa-miR-2681-5pAILANI<br>JUNmirna:hsa-miR-3653-3pAILANI<br>JUNmirna:hsa-miR-633AILANI<br>JUNmirna:hsa-miR-4720-3pAILANI<br>JUNmirna:hsa-miR-4804-3pAILANI<br>JUNmirna:hsa-miR-4328AILANI |
| **RELA** | RELA lipopolysaccharide chebi:CHEBI:16412 INDRA (drugs)<br>RELA curcumin chebi:CHEBI:3962 INDRA (drugs)<br>RELA bortezomib chebi:CHEBI:52717 INDRA (drugs)<br>RELA resveratrol chebi:CHEBI:27881 INDRA (drugs)<br>RELA fenofibrate chebi:CHEBI:5001 INDRA (drugs)<br>RELA nitric oxide chebi:CHEBI:16480 INDRA (drugs)<br>RELA (-)-epigallocatechin 3-gallate chebi:CHEBI:4806 INDRA (drugs)<br>RELA pyruvic acid chebi:CHEBI:32816 INDRA (drugs)<br>RELA parthenolide chebi:CHEBI:7939 INDRA (drugs)<br>RELA cyclosporin a chebi:CHEBI:4031 INDRA (drugs) | RELA daunorubicin drugbank:DB00694 TF analysis<br>RELA deoxycholic acid drugbank:DB03619 TF analysis<br>RELA dimethyl fumarate drugbank:DB08908 TF analysis<br>RELA gefitinib drugbank:DB00317 TF analysis<br>RELA oxygen drugbank:DB09140 TF analysis<br>RELA testosterone drugbank:DB00624 TF analysis | RELAmirna:hsa-miR-373-3pAILANI<br>RELAmirna:hsa-miR-124-3pAILANI<br>RELAmirna:hsa-miR-7-5pAILANI<br>RELAmirna:hsa-miR-96-5pAILANI<br>RELAmirna:hsa-miR-324-5pAILANI<br>RELAmirna:hsa-miR-30e-5pAILANI<br>RELAmirna:hsa-miR-320aAILANI<br>RELAmirna:hsa-miR-1236-3pAILANI<br>RELAmirna:hsa-miR-1261AILANI<br>RELAmirna:hsa-miR-3161AILANI<br>RELAmirna:hsa-miR-599AILANI<br>RELAmirna:hsa-miR-6077AILANI<br>RELAmirna:hsa-miR-3134AILANI<br>RELAmirna:hsa-miR-8082AILANI<br>RELAmirna:hsa-miR-4534AILANI<br>RELAmirna:hsa-miR-1303AILANI<br>RELAmirna:hsa-miR-6720-5pAILANI<br>RELAmirna:hsa-miR-6866-5pAILANI<br>RELAmirna:hsa-miR-6512-3pAILANI<br>RELAmirna:hsa-miR-877-5pAILANI<br>RELAmirna:hsa-miR-6849-3pAILANI<br>RELAmirna:hsa-miR-6859-5pAILANI<br>RELAmirna:hsa-miR-3916AILANI<br>RELAmirna:hsa-miR-3125AILANI |

| Protein / gene | INDRA drugs | Clinical trials/ TF analysis | AILANI |
|---|---|---|---|
| **NFKB1** | NFKB1curcuminchebi:CHEBI:3962INDRA (drugs)<br>NFKB1bortezomibchebi:CHEBI:52717INDRA (drugs)<br>NFKB1n-acetyl-l-cysteinechebi:CHEBI:28939INDRA (drugs)<br>NFKB1vorinostatchebi:CHEBI:45716INDRA (drugs)<br>NFKB12-(2,6-dioxopiperidin-3-yl)-1h-isoindole-1,3(2h)-dionechebi:CHEBI:74947 INDRA (drugs)<br>NFKB1lipopolysaccharidechebi:CHEBI:16412INDRA (drugs)<br>NFKB1salicylic acidchebi:CHEBI:16914INDRA (drugs)<br>NFKB1sodium(1+)chebi:CHEBI:29101INDRA (drugs)<br>NFKB1nitric oxidechebi:CHEBI:16480INDRA (drugs)<br>NFKB1acetylsalicylic acidchebi:CHEBI:15365INDRA (drugs) | NFKB1 thalidomide drugbank:DB01041 Clinical Trials DB, AILANI, TF analysis<br>NFKB1 bortezomib drugbank:DB00188 TF analysis<br>NFKB1 donepezil drugbank:DB00843 TF analysis<br>NFKB1 etoposide drugbank:DB00773 TF analysis<br>NFKB1 fish oil drugbank:DB13961 TF analysis<br>NFKB1 glycyrrhizin drugbank:DB13751 TF analysis<br>NFKB1 nitric oxide drugbank:DB00435 TF analysis<br>NFKB1 pseudoephedrine drugbank:DB00852 TF analysis<br>NFKB1 sulfasalazine drugbank:DB00795 TF analysis<br>NFKB1 triflusal drugbank:DB08814 TF analysis | NFKB1mirna:hsa-miR-9-5pAILANI<br>NFKB1mirna:hsa-miR-146a-5pAILANI<br>NFKB1mirna:hsa-miR-146b-5pAILANI<br>NFKB1mirna:hsa-miR-15a-5pAILANI<br>NFKB1mirna:hsa-let-7a-5pAILANI<br>NFKB1mirna:hsa-miR-16-5pAILANI<br>NFKB1mirna:hsa-miR-21-5pAILANI<br>NFKB1mirna:mmu-miR-210-3pAILANI<br>NFKB1mirna:hsa-miR-155-5pAILANI<br>NFKB1mirna:hsa-miR-26b-5pAILANI<br>NFKB1mirna:hsa-miR-92a-3pAILANI<br>NFKB1mirna:hsa-miR-199a-5pAILANI<br>NFKB1mirna:hsa-miR-139-5pAILANI |
| **STAT2** | STAT2ns-5chebi:CHEBI:81035INDRA (drugs)<br>STAT2cidofovir anhydrouschebi:CHEBI:3696INDRA (drugs)<br>STAT2arg-valchebi:CHEBI:73823INDRA (drugs)<br>STAT2acroleinchebi:CHEBI:15368INDRA (drugs)<br>STAT2ethanolchebi:CHEBI:16236INDRA (drugs) | STAT2 niclosamide drugbank:DB06803 TF analysis<br>STAT2 phenylalanine drugbank:DB00120 TF analysis<br>STAT2 tyrosine drugbank:DB00135 TF analysis | STAT2mirna:hsa-miR-335-5pAILANI<br>STAT2mirna:hsa-miR-125a-5pAILANI<br>STAT2mirna:hsa-let-7c-5pAILANI<br>STAT2mirna:hsa-miR-548sAILANI<br>STAT2mirna:hsa-miR-3926AILANI<br>STAT2mirna:hsa-miR-3190-5pAILANI<br>STAT2mirna:hsa-miR-372-5pAILANI<br>STAT2mirna:hsa-miR-371a-5pAILANI<br>STAT2mirna:hsa-miR-616-5pAILANI<br>STAT2mirna:hsa-miR-373-5pAILANI<br>STAT2mirna:hsa-miR-371b-5pAILANI<br>STAT2mirna:hsa-miR-8073AILANI<br>STAT2mirna:hsa-miR-221-5pAILANI<br>STAT2mirna:hsa-miR-3613-3pAILANI<br>STAT2mirna:hsa-miR-4668-3pAILANI<br>STAT2mirna:hsa-miR-548c-3pAILANI<br>STAT2mirna:hsa-miR-1273eAILANI<br>STAT2mirna:hsa-miR-4266AILANI<br>STAT2mirna:hsa-miR-6894-5pAILANI<br>STAT2mirna:hsa-miR-7160-5pAILANI<br>STAT2mirna:hsa-miR-202-3pAILANI<br>STAT2mirna:hsa-miR-98-5pAILANI<br>STAT2mirna:hsa-miR-4500AILANI<br>STAT2mirna:hsa-miR-4458AILANI<br>STAT2mirna:hsa-let-7i-5pAILANI<br>STAT2mirna:hsa-let-7g-5pAILANI<br>STAT2mirna:hsa-let-7f-5pAILANI<br>STAT2mirna:hsa-let-7e-5pAILANI<br>STAT2mirna:hsa-let-7d-5pAILANI<br>STAT2mirna:hsa-let-7b-5pAILANI<br>STAT2mirna:hsa-let-7a-5pAILANI<br>STAT2mirna:hsa-miR-6884-5pAILANI<br>STAT2mirna:hsa-miR-485-5pAILANI<br>STAT2mirna:hsa-miR-7162-3pAILANI<br>STAT2mirna:hsa-miR-6840-3pAILANI<br>STAT2mirna:hsa-miR-665AILANI<br>STAT2mirna:hsa-miR-4459AILANI<br>STAT2mirna:hsa-miR-4478AILANI<br>STAT2mirna:hsa-miR-4419bAILANI<br>STAT2mirna:hsa-miR-3929AILANI<br>STAT2mirna:hsa-miR-4779AILANI<br>STAT2mirna:hsa-miR-4695-5pAILANI<br>STAT2mirna:hsa-miR-4482-5pAILANI<br>STAT2mirna:hsa-miR-3130-5pAILANI<br>STAT2mirna:hsa-miR-1912AILANI<br>STAT2mirna:hsa-miR-1295b-5pAILANI |

| Protein / gene | INDRA drugs | Clinical trials/ TF analysis | AILANI |
|---|---|---|---|
| **FOSL1** | | | FOSL1mirna:hsa-miR-1207-3pAILANI<br>FOSL1mirna:hsa-miR-1275AILANI<br>FOSL1mirna:hsa-miR-138-5pAILANI<br>FOSL1mirna:hsa-miR-3150b-3pAILANI<br>FOSL1mirna:hsa-miR-3179AILANI<br>FOSL1mirna:hsa-miR-3202AILANI<br>FOSL1mirna:hsa-miR-33a-5pAILANI<br>FOSL1mirna:hsa-miR-34a-5pAILANI<br>FOSL1mirna:hsa-miR-34c-5pAILANI<br>FOSL1mirna:hsa-miR-4271AILANI<br>FOSL1mirna:hsa-miR-4447AILANI<br>FOSL1mirna:hsa-miR-4472AILANI<br>FOSL1mirna:hsa-miR-4525AILANI<br>FOSL1mirna:hsa-miR-4665-5pAILANI<br>FOSL1mirna:hsa-miR-4723-5pAILANI<br>FOSL1mirna:hsa-miR-4725-3pAILANI<br>FOSL1mirna:hsa-miR-4747-5pAILANI<br>FOSL1mirna:hsa-miR-4784AILANI<br>FOSL1mirna:hsa-miR-5010-5pAILANI<br>FOSL1mirna:hsa-miR-5196-5pAILANI<br>FOSL1mirna:hsa-miR-5698AILANI<br>FOSL1mirna:hsa-miR-625-5pAILANI<br>FOSL1mirna:hsa-miR-6780b-5pAILANI<br>FOSL1mirna:hsa-miR-6786-5pAILANI<br>FOSL1mirna:hsa-miR-6825-5pAILANI<br>FOSL1mirna:hsa-miR-6870-5pAILANI<br>FOSL1mirna:hsa-miR-7106-5pAILANI<br>FOSL1mirna:hsa-miR-7109-5pAILANI<br>FOSL1mirna:hsa-miR-7111-5pAILANI<br>FOSL1mirna:hsa-miR-765AILANI<br>FOSL1mirna:hsa-miR-92a-2-5pAILANI |
| **IRF9** | IRF9enzyme inhibitorchebi:CHEBI:23924INDRA (drugs)<br>IRF9cortisolchebi:CHEBI:17650INDRA (drugs)<br>IRF9cyclosporin achebi:CHEBI:4031INDRA (drugs)<br>IRF9staurosporinechebi:CHEBI:15738INDRA (drugs)<br>IRF9trichostatin achebi:CHEBI:46024INDRA (drugs) | IRF9 adenine drugbank:DB00173 TF analysis<br>IRF9 tyrosine drugbank:DB00135 TF analysis | |
| **IRF3** | IRF3lipopolysaccharidechebi:CHEBI:16412INDRA (drugs)<br>IRF3double-stranded rnachebi:CHEBI:67208INDRA (drugs)<br>IRF3curcuminchebi:CHEBI:3962INDRA (drugs) | | |

| Protein / gene | INDRA drugs | Clinical trials/ TF analysis | AILANI |
|---|---|---|---|
| **BACH1** | BACH1heminchebi:CHEBI:50385INDRA (drugs)<br>BACH1cadmium atomchebi:CHEBI:22977INDRA (drugs)<br>BACH1arsenite(3-)chebi:CHEBI:29866INDRA (drugs)<br>BACH1leptomycin bchebi:CHEBI:52646INDRA (drugs)<br>BACH1n-benzyloxycarbonyl-l-leucyl-l-leucyl-l-leucinalchebi:CHEBI:75142INDRA (drugs)<br>BACH1dihydroartemisininchebi:CHEBI:207229INDRA (drugs) | BACH1 cysteine drugbank:DB00151 TF analysis<br>BACH1 deferoxamine drugbank:DB00746 TF analysis<br>BACH1 iron drugbank:DB01592 TF analysis<br>BACH1 isoniazid drugbank:DB00951 TF analysis<br>BACH1 oxygen drugbank:DB09140 TF analysis | BACH1mirna:hsa-miR-155-5pAILANI<br>BACH1mirna:kshv-miR-K12-11-3pAILANI<br>BACH1mirna:hsa-miR-196a-5pAILANI<br>BACH1mirna:hsa-miR-769-5pAILANI<br>BACH1mirna:hsa-miR-125b-5pAILANI<br>BACH1mirna:hsa-miR-20a-5pAILANI<br>BACH1mirna:hsa-miR-185-5pAILANI<br>BACH1mirna:hsa-miR-190a-3pAILANI<br>BACH1mirna:hsa-miR-4306AILANI<br>BACH1mirna:hsa-miR-3924AILANI<br>BACH1mirna:hsa-miR-4644AILANI<br>BACH1mirna:hsa-miR-5011-5pAILANI<br>BACH1mirna:hsa-miR-8076AILANI<br>BACH1mirna:hsa-miR-302a-3pAILANI<br>BACH1mirna:hsa-miR-302b-3pAILANI<br>BACH1mirna:hsa-miR-302c-3pAILANI<br>BACH1mirna:hsa-miR-302d-3pAILANI<br>BACH1mirna:hsa-miR-372-3pAILANI<br>BACH1mirna:hsa-miR-373-3pAILANI<br>BACH1mirna:hsa-miR-520eAILANI<br>BACH1mirna:hsa-miR-520f-3pAILANI<br>BACH1mirna:hsa-miR-520a-3pAILANI<br>BACH1mirna:hsa-miR-520bAILANI<br>BACH1mirna:hsa-miR-520c-3pAILANI<br>BACH1mirna:hsa-miR-520d-3pAILANI<br>BACH1mirna:hsa-miR-539-5pAILANI<br>BACH1mirna:hsa-miR-302eAILANI<br>BACH1mirna:hsa-miR-4451AILANI<br>BACH1mirna:hsa-miR-6832-5pAILANI<br>BACH1mirna:hsa-let-7a-5pAILANI<br>BACH1mirna:hsa-let-7b-5pAILANI<br>BACH1mirna:hsa-let-7c-5pAILANI<br>BACH1mirna:hsa-let-7d-5pAILANI<br>BACH1mirna:hsa-let-7e-5pAILANI<br>BACH1mirna:hsa-let-7f-5pAILANI<br>BACH1mirna:hsa-miR-98-5pAILANI<br>BACH1mirna:hsa-let-7g-5pAILANI<br>BACH1mirna:hsa-let-7i-5pAILANI<br>BACH1mirna:hsa-miR-142-3pAILANI<br>BACH1mirna:hsa-miR-196b-5pAILANI<br>BACH1mirna:hsa-miR-643AILANI<br>BACH1mirna:hsa-miR-4280AILANI<br>BACH1mirna:hsa-miR-3668AILANI<br>BACH1mirna:hsa-miR-3935AILANI<br>BACH1mirna:hsa-miR-4458AILANI<br>BACH1mirna:hsa-miR-4500AILANI<br>BACH1mirna:hsa-miR-6504-3pAILANI<br>BACH1mirna:hsa-miR-6728-5pAILANI<br>BACH1mirna:hsa-miR-6871-5pAILANI<br>BACH1mirna:hsa-miR-6516-5pAILANI<br>BACH1mirna:hsa-miR-513a-3pAILANI<br>BACH1mirna:hsa-miR-559AILANI<br>BACH1mirna:hsa-miR-548b-5pAILANI<br>BACH1mirna:hsa-miR-590-3pAILANI<br>BACH1mirna:hsa-miR-548a-5pAILANI<br>BACH1mirna:hsa-miR-548c-5pAILANI<br>BACH1mirna:hsa-miR-548d-5pAILANI<br>BACH1mirna:hsa-miR-548j-5pAILANI<br>BACH1mirna:hsa-miR-548h-5pAILANI<br>BACH1mirna:hsa-miR-548iAILANI<br>BACH1mirna:hsa-miR-664a-5pAILANI<br>BACH1mirna:hsa-miR-513c-3pAILANI<br>BACH1mirna:hsa-miR-548wAILANI<br>BACH1mirna:hsa-miR-3606-3pAILANI<br>BACH1mirna:hsa-miR-548yAILANI<br>BACH1mirna:hsa-miR-548o-5pAILANI<br>BACH1mirna:hsa-miR-548abAILANI<br>BACH1mirna:hsa-miR-548ad-5pAILANI<br>BACH1mirna:hsa-miR-548ae-5pAILANI<br>BACH1mirna:hsa-miR-548akAILANI<br>BACH1mirna:hsa-miR-548am-5pAILANI<br>BACH1mirna:hsa-miR-4794AILANI<br>BACH1mirna:hsa-miR-548ap-5pAILANI<br>BACH1mirna:hsa-miR-548aq-5pAILANI<br>BACH1mirna:hsa-miR-548ar-5pAILANI<br>BACH1mirna:hsa-miR-548as-5pAILANI<br>BACH1mirna:hsa-miR-5580-5pAILANI<br>BACH1mirna:hsa-miR-548au-5pAILANI |

| Protein / gene | INDRA drugs | Clinical trials/ TF analysis | AILANI |
|---|---|---|---|
| **TBP** | TBP2-(4-methyl-1,4-diazepan-1-yl)-n-[(5-methyl-2-pyrazinyl)methyl]-5-oxo-[1,3] benzothiazolo[3,2-a][1,8]naphthyridine-6-carboxamidechebi:CHEBI:95090INDRA (drugs)<br>TBPglutathionechebi:CHEBI:16856INDRA (drugs)<br>TBPcalcium(2+)chebi:CHEBI:29108INDRA (drugs)<br>TBPhistonefplx:HistoneINDRA (drugs)<br>TBPacetyl-coachebi:CHEBI:15351INDRA (drugs)<br>TBPatpchebi:CHEBI:15422INDRA (drugs) | TBP cisplatin drugbank:DB00515 TF analysis<br>TBP dexamethasone drugbank:DB01234 TF analysis<br>TBP formaldehyde drugbank:DB03843 TF analysis<br>TBP glycerol drugbank:DB09462 TF analysis<br>TBP iron drugbank:DB01592 TF analysis<br>TBP levothyroxine drugbank:DB00451 TF analysis<br>TBP oxygen drugbank:DB09140 TF analysis<br>TBP potassium drugbank:DB14500 TF analysis<br>TBP progesterone drugbank:DB00396 TF analysis<br>TBP tocopherol drugbank:DB11251 TF analysis<br>TBP zinc drugbank:DB01593 TF analysis<br>TBP adenine drugbank:DB00173 TF analysis<br>TBP cysteine drugbank:DB00151 TF analysis<br>TBP phenylalanine drugbank:DB00120 TF analysis<br>TBP vitamin d drugbank:DB11094 TF analysis | |
| **TCF12** | TCF12uridine 5'-monophosphatechebi:CHEBI:16695INDRA (drugs) | | TCF12mirna:hsa-miR-204-5pAILANI<br>TCF12mirna:hsa-miR-211-5pAILANI<br>TCF12mirna:hsa-miR-375AILANI<br>TCF12mirna:hsa-miR-155-5pAILANI<br>TCF12mirna:hsa-miR-26b-5pAILANI<br>TCF12mirna:hsa-miR-1224-3pAILANI<br>TCF12mirna:hsa-miR-548vAILANI<br>TCF12mirna:hsa-miR-33a-3pAILANI<br>TCF12mirna:hsa-miR-139-5pAILANI<br>TCF12mirna:hsa-miR-8485AILANI |
| **IFIH1** | IFIH1flavonoidchebi:CHEBI:47916INDRA (drugs)<br>IFIH1poly i:cpubchem.compound:86583374INDRA (drugs)<br>IFIH1lactatechebi:CHEBI:24996INDRA (drugs)<br>IFIH1cisplatinchebi:CHEBI:27899INDRA (drugs)<br>IFIH1l-prolylglycinechebi:CHEBI:61695INDRA (drugs) | | IFIH1mirna:hsa-miR-26b-5pAILANI<br>IFIH1mirna:hsa-miR-424-5pAILANI<br>IFIH1mirna:hsa-miR-15b-5pAILANI<br>IFIH1mirna:hsa-miR-29b-3pAILANI |
| **OAS1** | OAS1double-stranded rnachebi:CHEBI:67208INDRA (drugs)<br>OAS1zinc atomchebi:CHEBI:27363INDRA (drugs)<br>OAS1heparinpubchem.compound:22833565INDRA (drugs) | no clinical trial | OAS1 cysteine-s-acetamide drugbank:DB02987<br>INDRA (drugs), AILANI OAS1mirna:hsa-miR-335-5pAILANI |
| **OAS2** | OAS2zinc atomchebi:CHEBI:27363INDRA (drugs)<br>OAS2etanerceptchebi:CHEBI:4875INDRA (drugs)<br>OAS2sirolimuschebi:CHEBI:9168INDRA (drugs) | no clinical trial | OAS2mirna:hsa-miR-335-5pAILANI<br>OAS2mirna:hsa-miR-132-3pAILANI<br>OAS2mirna:hsa-miR-7-5pAILANI<br>OAS2mirna:hsa-miR-620AILANI<br>OAS2mirna:hsa-miR-4531AILANI<br>OAS2mirna:hsa-miR-1270AILANI<br>OAS2mirna:hsa-miR-573AILANI<br>OAS2mirna:hsa-miR-3616-5pAILANI<br>OAS2mirna:hsa-miR-5192AILANI<br>OAS2mirna:hsa-miR-4644AILANI<br>OAS2mirna:hsa-miR-4306AILANI<br>OAS2mirna:hsa-miR-185-5pAILANI<br>OAS2mirna:hsa-miR-6823-3pAILANI<br>OAS2mirna:hsa-miR-2114-3pAILANI<br>OAS2mirna:hsa-miR-4635AILANI<br>OAS2mirna:hsa-miR-4670-3pAILANI<br>OAS2mirna:hsa-miR-6751-3pAILANI<br>OAS2mirna:hsa-miR-1199-5pAILANI<br>OAS2mirna:hsa-miR-148a-5pAILANI |
| **OAS3** | OAS3etanerceptchebi:CHEBI:4875INDRA (drugs) | no clinical trial | OAS3mirna:hsa-miR-502-5pAILANI<br>OAS3mirna:hsa-miR-1273fAILANI<br>OAS3mirna:hsa-miR-4708-5pAILANI<br>OAS3mirna:hsa-miR-6088AILANI<br>OAS3mirna:hsa-miR-4770AILANI<br>OAS3mirna:hsa-miR-143-3pAILANI |
| **IRF7** | IRF7lansoprazolechebi:CHEBI:6375INDRA (drugs)<br>IRF7nickel(2+)chebi:CHEBI:49786INDRA (drugs)<br>IRF7aktippubchem.compound:5469318INDRA (drugs)<br>IRF7all-trans-retinoic acidchebi:CHEBI:15367INDRA (drugs)<br>IRF7theophyllinechebi:CHEBI:28177INDRA (drugs) | | IRF7mirna:hsa-miR-146a-5pAILANI |

| **Protein / gene** | **INDRA drugs** | **Clinical trials/ TF analysis** | **AILANI** |
|---|---|---|---|
| **IFNB1 RNA** | IFNB1beta-d-glucosechebi:CHEBI:15903INDRA (drugs)<br>IFNB1nimesulidechebi:CHEBI:44445INDRA (drugs)<br>IFNB1ifosfamidechebi:CHEBI:5864INDRA (drugs)<br>IFNB1lipopolysaccharidechebi:CHEBI:16412INDRA (drugs)<br>IFNB1poly i:cpubchem.compound:86583374INDRA (drugs)<br>IFNB1chloroquinechebi:CHEBI:3638INDRA (drugs)<br>IFNB1arg-valchebi:CHEBI:73823INDRA (drugs)<br>IFNB1simvastatinchebi:CHEBI:9150INDRA (drugs) | | IFNB1beta-d-glucosedrugbank:DB02379AILANI<br>IFNB1mirna:hsa-let-7b-5pAILANI<br>IFNB1mirna:hsa-miR-26a-5pAILANI<br>IFNB1mirna:hsa-miR-145-5pAILANI<br>IFNB1mirna:hsa-miR-34a-5pAILANI<br>IFNB1mirna:hsa-miR-607AILANI<br>IFNB1mirna:hsa-miR-6080AILANI<br>IFNB1mirna:hsa-miR-4698AILANI<br>IFNB1mirna:hsa-miR-8063AILANI<br>IFNB1mirna:hsa-miR-513c-3pAILANI<br>IFNB1mirna:hsa-miR-513a-3pAILANI<br>IFNB1mirna:hsa-miR-3606-3pAILANI<br>IFNB1mirna:hsa-miR-4282AILANI<br>IFNB1mirna:hsa-miR-548c-3pAILANI |
| **TLR7** | TLR7hydroxychloroquinedrugbank:DB01611Clinical Trials DB, AILANI<br>TLR7isatoribinedrugbank:DB04860INDRA (drugs), AILANI | | TLR7imiquimoddrugbank:DB00724AILANITLR7mirna:hsa-miR-758-3pAILANI<br>TLR7mirna:hsa-miR-586AILANI<br>TLR7mirna:hsa-miR-8087AILANI<br>TLR7mirna:hsa-miR-1228-3pAILANI<br>TLR7mirna:hsa-miR-5089-5pAILANI<br>TLR7mirna:hsa-miR-5589-5pAILANI<br>TLR7mirna:hsa-miR-4731-5pAILANI<br>TLR7mirna:hsa-miR-6778-3pAILANI<br>TLR7mirna:hsa-miR-6506-5pAILANI<br>TLR7mirna:hsa-miR-619-5pAILANI<br>TLR7mirna:hsa-miR-150-5pAILANI<br>TLR7mirna:hsa-miR-7107-5pAILANI<br>TLR7mirna:hsa-miR-1234-3pAILANI<br>TLR7mirna:hsa-miR-6747-3pAILANI<br>TLR7mirna:hsa-miR-186-3pAILANI<br>TLR7mirna:hsa-miR-520hAILANI<br>TLR7mirna:hsa-miR-520g-3pAILANI<br>TLR7mirna:hsa-miR-512-3pAILANI<br>TLR7mirna:hsa-miR-520eAILANI<br>TLR7mirna:hsa-miR-520d-3pAILANI<br>TLR7mirna:hsa-miR-520c-3pAILANI<br>TLR7mirna:hsa-miR-520bAILANI<br>TLR7mirna:hsa-miR-520a-3pAILANI<br>TLR7mirna:hsa-miR-373-3pAILANI<br>TLR7mirna:hsa-miR-372-3pAILANI<br>TLR7mirna:hsa-miR-302eAILANI<br>TLR7mirna:hsa-miR-302d-3pAILANI<br>TLR7mirna:hsa-miR-302c-3pAILANI<br>TLR7mirna:hsa-miR-302b-3pAILANI<br>TLR7mirna:hsa-miR-302a-3pAILANI<br>TLR7mirna:hsa-miR-93-5pAILANI<br>TLR7mirna:hsa-miR-526b-3pAILANI<br>TLR7mirna:hsa-miR-519d-3pAILANI<br>TLR7mirna:hsa-miR-20b-5pAILANI<br>TLR7mirna:hsa-miR-20a-5pAILANI<br>TLR7mirna:hsa-miR-17-5pAILANI<br>TLR7mirna:hsa-miR-106b-5pAILANI<br>TLR7mirna:hsa-miR-106a-5pAILANI |
| **TLR9** | TLR9 chloroquine chebi:CHEBI:3638 INDRA (drugs)<br>TLR9 hydroxychloroquine chebi:CHEBI:5801 INDRA (drugs)<br>TLR9 agatolimod chembl.compound:CHEMBL2103792 INDRA (drugs)<br>TLR9 cpg 10101 drugbank:DB05530 INDRA (drugs)<br>TLR9 golotimod drugbank:DB05475 INDRA (drugs)<br>TLR9 lipopolysaccharide chebi:CHEBI:16412 INDRA (drugs)<br>TLR9 3',5'-cyclic amp chebi:CHEBI:17489 INDRA (drugs)<br>TLR9 gold(1+) chebi:CHEBI:49482 INDRA (drugs)<br>TLR9 methamphetamine chebi:CHEBI:6809 INDRA (drugs)<br>TLR9 pyridoxamine chebi:CHEBI:16410 INDRA (drugs)* | TLR9 chloroquine drugbank:DB00608 Clinical Trials DB, AILANI<br>TLR9 hydroxychloroquine drugbank:DB01611 Clinical Trials DB, AILANI | |
| **TREML4** | none | none | none |

| **Protein / gene** | **INDRA drugs** | **Clinical trials/ TF analysis** | **AILANI** |
|---|---|---|---|
| **TBK1** | TBK1 n-[3-[[5-iodo-4-[3-[[oxo(thiophen-2-yl)methyl]amino]propylamino]-2-pyrimidin<br>TBK1 amlexanox chebi:CHEBI:31205 INDRA (drugs)<br>TBK1 n-[3-[[5-cyclopropyl-2-[3-(4-morpholinylmethyl)anilino]-4-pyrimidinyl]amino]pr<br>TBK1 sunitinib chebi:CHEBI:38940 INDRA (drugs)<br>TBK1 sunitinib malate chembl.compound:CHEMBL1567 INDRA (drugs)<br>TBK1 resveratrol chebi:CHEBI:27881 INDRA (drugs)<br>TBK1 thymoquinone chebi:CHEBI:113532 INDRA (drugs)<br>TBK1 cholesterol chebi:CHEBI:16113 INDRA (drugs)<br>TBK1 glucose chebi:CHEBI:17234 INDRA (drugs)<br>TBK1 glucocorticoid chebi:CHEBI:24261 INDRA (drugs) | none | none |
| **AGTR1/2** | AGTR1losartanchebi:CHEBI:6541INDRA (drugs)<br>AGTR1telmisartanchebi:CHEBI:9434INDRA (drugs)<br>AGTR1irbesartanchebi:CHEBI:5959INDRA (drugs)<br>AGTR1valsartanchebi:CHEBI:9927INDRA (drugs)<br>AGTR1candesartanchebi:CHEBI:3347INDRA (drugs)<br>AGTR117alpha-ethynylestradiolchebi:CHEBI:4903INDRA (drugs)<br>AGTR1estrogenchebi:CHEBI:50114INDRA (drugs)<br>AGTR1nitric oxidechebi:CHEBI:16480INDRA (drugs)<br>AGTR1glucosechebi:CHEBI:17234INDRA (drugs)<br>AGTR11,4-dithiothreitolchebi:CHEBI:18320INDRA (drugs) | angiotensin ii drugbank:DB11842 Clinical Trials DB | AGTR1 tasosartan drugbank:DB01349 AILANI<br>AGTR1 saprisartan drugbank:DB01347 AILANI<br>AGTR1 forasartan drugbank:DB01342 AILANI<br>AGTR1 eprosartan drugbank:DB00876 AILANI<br>AGTR1 irbesartan drugbank:DB01029 AILANI<br>AGTR1 azilsartan medoxomil drugbank:DB08822<br>AILANI<br>AGTR1 olmesartan drugbank:DB00275 AILANI<br>AGTR1 mirna:hsa-miR-155-5p AILANI<br>AGTR1 mirna:hsa-miR-124-3p AILANI<br>AGTR1 mirna:hsa-miR-26b-5p AILANI |
| **STAT1** | STAT1 (-)-epigallocatechin 3-gallate chebi:CHEBI:4806 INDRA (drugs)<br>STAT1 simvastatin chebi:CHEBI:9150 INDRA (drugs)<br>STAT1 herbimycin chebi:CHEBI:5674 INDRA (drugs)<br>STAT1 colchicine chebi:CHEBI:23359 INDRA (drugs)<br>STAT1 zinc atom chebi:CHEBI:27363 INDRA (drugs)<br>STAT1 lipopolysaccharide chebi:CHEBI:16412 INDRA (drugs)<br>STAT1 tofacitinib chebi:CHEBI:71200 INDRA (drugs)<br>STAT1 daidzein chebi:CHEBI:28197 INDRA (drugs)<br>STAT1 quercetin chebi:CHEBI:16243 INDRA (drugs)<br>STAT1 butyric acid chebi:CHEBI:30772 INDRA (drugs) | STAT1 biotin drugbank:DB00121 TF analysis<br>STAT1 calcitriol drugbank:DB00136 TF analysis<br>STAT1 cholesterol drugbank:DB04540 TF analysis<br>STAT1 cisplatin drugbank:DB00515 TF analysis<br>STAT1 curcumin drugbank:DB11672 TF analysis<br>STAT1 cyclosporin a drugbank:DB00091 TF analysis<br>STAT1 cysteine drugbank:DB00151 TF analysis<br>STAT1 deferoxamine drugbank:DB00746 TF analysis<br>STAT1 dexamethasone drugbank:DB01234 TF analysis<br>STAT1 doxorubicin drugbank:DB00997 TF analysis<br>STAT1 ethanol drugbank:DB00898 TF analysis<br>STAT1 etoposide drugbank:DB00773 TF analysis<br>STAT1 fludarabine drugbank:DB01073 TF analysis<br>STAT1 imatinib drugbank:DB00619 TF analysis<br>STAT1 imiquimod drugbank:DB00724 TF analysis<br>STAT1 indomethacin drugbank:DB00328 TF analysis<br>STAT1 methimazole drugbank:DB00763 TF analysis<br>STAT1 niclosamide drugbank:DB06803 TF analysis<br>STAT1 nitric oxide drugbank:DB00435 TF analysis<br>STAT1 ornithine drugbank:DB00129 TF analysis<br>STAT1 oxygen drugbank:DB09140 TF analysis<br>STAT1 phenylalanine drugbank:DB00120 TF analysis<br>STAT1 prazosin drugbank:DB00457 TF analysis<br>STAT1 progesterone drugbank:DB00396 TF analysis<br>STAT1 ribavirin drugbank:DB00811 TF analysis<br>STAT1 rosiglitazone drugbank:DB00412 TF analysis<br>STAT1 threonine drugbank:DB00156 TF analysis<br>STAT1 tyrosine drugbank:DB00135 TF analysis<br>STAT1 valsartan drugbank:DB00177 TF analysis<br>STAT1 vitamin d drugbank:DB11094 TF analysis<br>STAT1 zinc drugbank:DB01593 TF analysis | STAT1 mirna:hsa-miR-145-5p AILANI<br>STAT1 mirna:hsa-miR-146a-5p AILANI<br>STAT1 mirna:hsa-miR-34a-5p AILANI<br>STAT1 mirna:hsa-miR-615-3p AILANI<br>STAT1 mirna:hsa-miR-653-5p AILANI<br>STAT1 mirna:hsa-miR-203b-3p AILANI<br>STAT1 mirna:hsa-miR-7158-3p AILANI<br>STAT1 mirna:hsa-miR-4693-5p AILANI<br>STAT1 mirna:hsa-miR-5009-3p AILANI<br>STAT1 mirna:hsa-miR-501-5p AILANI<br>STAT1 mirna:hsa-miR-1183 AILANI<br>STAT1 mirna:hsa-miR-500a-5p AILANI<br>STAT1 mirna:hsa-miR-605-5p AILANI |
| **IRF3** | IRF3Helicaseuniprot:PRO_0000449630INDRA (EMMAA)<br>IRF3Uridylate-specific endoribonucleaseuniprot:PRO_0000449632INDRA (EMMAA)<br>IRF3lipopolysaccharidechebi:CHEBI:16412INDRA (drugs)<br>IRF3double-stranded rnachebi:CHEBI:67208INDRA (drugs)<br>IRF3curcuminchebi:CHEBI:3962INDRA (drugs) | | |
| **BAX** | BAXresveratrolchebi:CHEBI:27881INDRA (drugs)<br>BAXsirolimuschebi:CHEBI:9168INDRA (drugs)<br>BAXbortezomibchebi:CHEBI:52717INDRA (drugs)<br>BAXenzyme inhibitorchebi:CHEBI:23924INDRA (drugs)<br>BAXtroglitazonechebi:CHEBI:9753INDRA (drugs)<br>BAXnacchebi:CHEBI:7421INDRA (drugs)<br>BAXcyclosporin achebi:CHEBI:4031INDRA (drugs)<br>BAXglucosechebi:CHEBI:17234INDRA (drugs)<br>BAXmelatoninchebi:CHEBI:16796INDRA (drugs)<br>BAXreactive oxygen specieschebi:CHEBI:26523INDRA (drugs) | | BAXmirna:hsa-miR-504-5pAILANI<br>BAXmirna:hsa-miR-148b-3pAILANI<br>BAXmirna:hsa-miR-122-5pAILANI<br>BAXmirna:hsa-miR-1-3pAILANI<br>BAXmirna:hsa-miR-30a-5pAILANI<br>BAXmirna:hsa-miR-365a-3pAILANI<br>BAXmirna:hsa-miR-128-3pAILANI<br>BAXmirna:hsa-miR-214-3pAILANI<br>BAXmirna:hsa-miR-766-3pAILANI<br>BAXmirna:hsa-miR-298AILANI |

| Protein / gene | INDRA drugs | Clinical trials/ TF analysis | AILANI |
|---|---|---|---|
| **FADD** | FADDenzyme inhibitorchebi:CHEBI:23924INDRA (drugs)<br>FADDml-7chebi:CHEBI:78761INDRA (drugs)<br>FADDml 9mesh:C056218INDRA (drugs)<br>FADDpotassium atomchebi:CHEBI:26216INDRA (drugs)<br>FADDcocainechebi:CHEBI:27958INDRA (drugs)<br>FADDtridecanechebi:CHEBI:35998INDRA (drugs)<br>FADDmtor inhibitorchebi:CHEBI:68481INDRA (drugs)<br>FADDnacchebi:CHEBI:7421INDRA (drugs) | | FADDmirna:hsa-miR-146a-5pAILANI<br>FADDmirna:hsa-miR-155-5pAILANI<br>FADDmirna:hsa-miR-1-3pAILANI<br>FADDmirna:hsa-miR-7-5pAILANI<br>FADDmirna:hsa-miR-15b-5pAILANI<br>FADDmirna:hsa-miR-128-3pAILANI |
| **AKT1** | AKT1ly294002chebi:CHEBI:65329INDRA (drugs)<br>AKT1staurosporinechebi:CHEBI:15738INDRA (drugs)<br>AKT1mk-2206chebi:CHEBI:67271INDRA (drugs)<br>AKT1curcuminchebi:CHEBI:3962INDRA (drugs)<br>AKT1resveratrolchebi:CHEBI:27881INDRA (drugs)<br>AKT1sirolimuschebi:CHEBI:9168INDRA (drugs)<br>AKT1ceramidechebi:CHEBI:17761INDRA (drugs)<br>AKT1hexadecanoic acidchebi:CHEBI:15756INDRA (drugs)<br>AKT13',5'-cyclic ampchebi:CHEBI:17489INDRA (drugs)<br>AKT1tanespimycinchebi:CHEBI:64153INDRA (drugs) | | AKT1 adenosine triphosphate drugbank:DB00171 AILANI<br>AKT1 inositol 1,3,4,5-tetrakisphosphate drugbank:DB01863 AILANI<br>AKT1 arsenic trioxide drugbank:DB01169 AILANI<br>AKT1 n-[2-(5-methyl-4h-1,2,4-triazol-3-yl)phenyl]-7h-pyrrolo[2,3-d]pyrimidin-4-amine drugbank:DB07584 AILANI<br>AKT1 5-(5-chloro-7h-pyrrolo[2,3-d]pyrimidin-4-yl)-4,5,6,7-tetrahydro-1h-imidazo[4,5-c]pyridine drugbank:DB07585 AILANI AKT1mirna:hsa-miR-199a-3pAILANI<br>AKT1mirna:hsa-miR-125b-5pAILANI<br>AKT1mirna:hsa-miR-185-5pAILANI<br>AKT1mirna:hsa-miR-149-3pAILANI<br>AKT1mirna:hsa-miR-451aAILANI<br>AKT1mirna:hsa-miR-302a-3pAILANI<br>AKT1mirna:hsa-miR-302b-3pAILANI<br>AKT1mirna:hsa-miR-302c-3pAILANI<br>AKT1mirna:hsa-miR-302d-3pAILANI<br>AKT1mirna:hsa-miR-143-3pAILANI<br>AKT1mirna:hsa-miR-26b-5pAILANI<br>AKT1mirna:hsa-miR-193b-3pAILANI<br>AKT1mirna:hsa-miR-100-5pAILANI<br>AKT1mirna:hsa-miR-19a-3pAILANI<br>AKT1mirna:hsa-miR-133bAILANI<br>AKT1mirna:hsa-miR-708-5pAILANI<br>AKT1mirna:hsa-miR-105-5pAILANI<br>AKT1mirna:hsa-miR-27a-5pAILANI<br>AKT1mirna:hsa-miR-192-3pAILANI<br>AKT1mirna:hsa-miR-155-5pAILANI<br>AKT1mirna:hsa-miR-496AILANI<br>AKT1mirna:hsa-miR-4742-3pAILANI<br>AKT1mirna:hsa-miR-654-3pAILANI<br>AKT1mirna:hsa-miR-4757-3pAILANI<br>AKT1mirna:hsa-miR-365b-3pAILANI<br>AKT1mirna:hsa-miR-365a-3pAILANI<br>AKT1mirna:hsa-miR-3191-5pAILANI<br>AKT1mirna:hsa-miR-6873-3pAILANI<br>AKT1mirna:hsa-miR-6503-3pAILANI<br>AKT1mirna:hsa-miR-422aAILANI<br>AKT1mirna:hsa-miR-378iAILANI<br>AKT1mirna:hsa-miR-378hAILANI<br>AKT1mirna:hsa-miR-378fAILANI<br>AKT1mirna:hsa-miR-378eAILANI<br>AKT1mirna:hsa-miR-378dAILANI<br>AKT1mirna:hsa-miR-378cAILANI<br>AKT1mirna:hsa-miR-378bAILANI<br>AKT1mirna:hsa-miR-378a-3pAILANI<br>AKT1mirna:hsa-miR-4638-3pAILANI<br>AKT1mirna:hsa-miR-6824-3pAILANI<br>AKT1mirna:hsa-miR-6764-3pAILANI<br>AKT1mirna:hsa-miR-330-5pAILANI<br>AKT1mirna:hsa-miR-326AILANI<br>AKT1mirna:hsa-miR-6817-3pAILANI<br>AKT1mirna:hsa-miR-518c-5pAILANI<br>AKT1mirna:hsa-miR-7110-3pAILANI<br>AKT1mirna:hsa-miR-6866-3pAILANI<br>AKT1mirna:hsa-miR-188-5pAILANI<br>AKT1mirna:hsa-miR-625-3pAILANI |

| Protein / gene | INDRA drugs | Clinical trials/ TF analysis | AILANI |
|---|---|---|---|
| STAT3 | STAT3resveratrolchebi:CHEBI:27881INDRA (drugs)<br>STAT3sorafenibchebi:CHEBI:50924INDRA (drugs)<br>STAT3s3i-201chebi:CHEBI:91224INDRA (drugs)<br>STAT3(-)-epigallocatechin 3-gallatechebi:CHEBI:4806INDRA (drugs)<br>STAT3cisplatinchebi:CHEBI:27899INDRA (drugs)<br>STAT3curcuminchebi:CHEBI:3962INDRA (drugs)<br>STAT3statticchebi:CHEBI:86989INDRA (drugs)<br>STAT3metforminchebi:CHEBI:6801INDRA (drugs)<br>STAT3sunitinibchebi:CHEBI:38940INDRA (drugs)<br>STAT3dihydroartemisininchebi:CHEBI:207229INDRA (drugs) | | STAT3mirna:hsa-miR-20b-5pAILANI<br>STAT3mirna:hsa-miR-125b-5pAILANI<br>STAT3mirna:hsa-miR-337-3pAILANI<br>STAT3mirna:hsa-miR-155-5pAILANI<br>STAT3mirna:hsa-miR-93-5pAILANI<br>STAT3mirna:hsa-miR-21-5pAILANI<br>STAT3mirna:hsa-miR-92a-3pAILANI<br>STAT3mirna:hsa-miR-20a-5pAILANI<br>STAT3mirna:hsa-let-7e-5pAILANI<br>STAT3mirna:hsa-miR-124-3pAILANI<br>STAT3mirna:hsa-miR-130b-3pAILANI<br>STAT3mirna:hsa-miR-106a-5pAILANI<br>STAT3mirna:hsa-miR-106b-5pAILANI<br>STAT3mirna:hsa-miR-874-3pAILANI<br>STAT3mirna:hsa-miR-4516AILANI<br>STAT3mirna:hsa-miR-17-5pAILANI<br>STAT3mirna:hsa-miR-181a-5pAILANI<br>STAT3mirna:hsa-miR-1234-3pAILANI<br>STAT3mirna:hsa-miR-1181AILANI<br>STAT3mirna:hsa-miR-6878-5pAILANI<br>STAT3mirna:hsa-miR-5681aAILANI<br>STAT3mirna:hsa-miR-6837-5pAILANI<br>STAT3mirna:hsa-miR-4685-5pAILANI<br>STAT3mirna:hsa-miR-7113-5pAILANI<br>STAT3mirna:hsa-miR-6754-5pAILANI<br>STAT3mirna:hsa-miR-4270AILANI<br>STAT3mirna:hsa-miR-4441AILANI<br>STAT3mirna:hsa-miR-922AILANI<br>STAT3mirna:hsa-miR-6760-5pAILANI<br>STAT3mirna:hsa-miR-1288-5pAILANI<br>STAT3mirna:hsa-miR-6871-3pAILANI<br>STAT3mirna:hsa-miR-6809-3pAILANI<br>STAT3mirna:hsa-miR-6833-3pAILANI<br>STAT3mirna:hsa-miR-4768-5pAILANI<br>STAT3mirna:hsa-miR-1178-3pAILANI<br>STAT3mirna:hsa-miR-7158-5pAILANI<br>STAT3mirna:hsa-miR-4684-5pAILANI<br>STAT3mirna:hsa-miR-138-2-3pAILANI<br>STAT3mirna:hsa-miR-371b-3pAILANI<br>STAT3mirna:hsa-miR-548nAILANI<br>STAT3mirna:hsa-miR-6875-3pAILANI<br>STAT3mirna:hsa-miR-5581-3pAILANI<br>STAT3mirna:hsa-miR-4778-3pAILANI<br>STAT3mirna:hsa-miR-7110-3pAILANI<br>STAT3mirna:hsa-miR-6845-3pAILANI<br>STAT3mirna:hsa-miR-877-3pAILANI<br>STAT3mirna:hsa-miR-618AILANI<br>STAT3mirna:hsa-miR-6807-5pAILANI<br>STAT3mirna:hsa-miR-4720-3pAILANI<br>STAT3mirna:hsa-miR-6762-3pAILANI<br>STAT3mirna:hsa-miR-4804-3pAILANI<br>STAT3mirna:hsa-miR-6872-3pAILANI<br>STAT3mirna:hsa-miR-7151-3pAILANI<br>STAT3mirna:hsa-miR-5095AILANI<br>STAT3mirna:hsa-miR-365b-5pAILANI<br>STAT3mirna:hsa-miR-365a-5pAILANI<br>STAT3mirna:hsa-miR-8052AILANI<br>STAT3mirna:hsa-miR-3199AILANI<br>STAT3mirna:hsa-miR-6789-3pAILANI<br>STAT3mirna:hsa-miR-500b-3pAILANI<br>STAT3mirna:hsa-miR-4438AILANI<br>STAT3mirna:hsa-miR-519d-3pAILANI |
| ISG15 | ISG15chebi:CHEBI:15367all-trans-retinoic acidINDRA (drugs)<br>ISG15chebi:CHEBI:16234hydroxideINDRA (drugs)<br>ISG15chebi:CHEBI:16480nitric oxideINDRA (drugs)<br>ISG15chebi:CHEBI:26536retinoic acidINDRA (drugs)<br>ISG15chebi:CHEBI:3165bradykininINDRA (drugs)<br>ISG15chebi:CHEBI:80630irinotecanINDRA (drugs) | | ISG15mirna:hsa-miR-1-3pAILANI<br>ISG15mirna:hsa-miR-146a-5pAILANI |

| Protein / gene | INDRA drugs | Clinical trials/ TF analysis | AILANI |
|---|---|---|---|
| **JUND** | JUNDchebi:CHEBI:15414s-adenosyl-l-methionineINDRA (drugs)<br>JUNDchebi:CHEBI:15864luteolinINDRA (drugs)<br>JUNDchebi:CHEBI:27881resveratrolINDRA (drugs)<br>JUNDchebi:CHEBI:27958cocaineINDRA (drugs)<br>JUNDchebi:CHEBI:37537phorbol 13-acetate 12-myristateINDRA (drugs)<br>JUNDchebi:CHEBI:5181fucoidanINDRA (drugs) JUNDpubchem.compound:4611oxamflatinINDRA (drugs)<br>JUNDpubchem.compound:5353940sb202190INDRA (drugs) | | JUNDmirna:hsa-miR-186-5pAILANI<br>JUNDmirna:hsa-miR-1908-5pAILANI<br>JUNDmirna:hsa-miR-3173-3pAILANI<br>JUNDmirna:hsa-miR-3185AILANI<br>JUNDmirna:hsa-miR-335-5pAILANI<br>JUNDmirna:hsa-miR-4271AILANI<br>JUNDmirna:hsa-miR-4695-5pAILANI<br>JUNDmirna:hsa-miR-4707-5pAILANI<br>JUNDmirna:hsa-miR-4725-3pAILANI<br>JUNDmirna:hsa-miR-4755-3pAILANI<br>JUNDmirna:hsa-miR-4778-3pAILANI<br>JUNDmirna:hsa-miR-625-5pAILANI<br>JUNDmirna:hsa-miR-663aAILANI<br>JUNDmirna:hsa-miR-6780b-5pAILANI<br>JUNDmirna:hsa-miR-6787-5pAILANI<br>JUNDmirna:hsa-miR-6875-3pAILANI<br>JUNDmirna:hsa-miR-6891-5pAILANI |

| Protein / gene | INDRA drugs | Clinical trials/ TF analysis | AILANI |
|---|---|---|---|
| **LNPEP** | LNPEP chebi:CHEBI:16236 ethanol INDRA (drugs)<br>LNPEP chebi:CHEBI:16794 scopolamine INDRA (drugs)<br>LNPEP chebi:CHEBI:16914 salicylic acid INDRA (drugs)<br>LNPEP chebi:CHEBI:26523 reactive oxygen species INDRA (drugs)<br>LNPEP chebi:CHEBI:36773 camphor INDRA (drugs)<br>LNPEP chebi:CHEBI:95044 (2s)-2-[[(2r)-2-[(1s)-1-hydroxy-2-(hydroxyamino)-2-oxoethyl]-4-methyl-1-oxopentyl]amino]-2-phenylacetic acid cyclopentyl ester | | LNPEPmirna:hsa-miR-1-3pAILANI<br>LNPEPmirna:hsa-miR-1197AILANI<br>LNPEPmirna:hsa-miR-1237-3pAILANI<br>LNPEPmirna:hsa-miR-1250-3pAILANI<br>LNPEPmirna:hsa-miR-1253AILANI<br>LNPEPmirna:hsa-miR-128-3pAILANI<br>LNPEPmirna:hsa-miR-133a-5pAILANI<br>LNPEPmirna:hsa-miR-138-5pAILANI<br>LNPEPmirna:hsa-miR-1468-3pAILANI<br>LNPEPmirna:hsa-miR-148a-3pAILANI<br>LNPEPmirna:hsa-miR-148b-3pAILANI<br>LNPEPmirna:hsa-miR-152-3pAILANI<br>LNPEPmirna:hsa-miR-153-5pAILANI<br>LNPEPmirna:hsa-miR-18a-3pAILANI<br>LNPEPmirna:hsa-miR-216a-3pAILANI<br>LNPEPmirna:hsa-miR-218-5pAILANI<br>LNPEPmirna:hsa-miR-30e-3pAILANI<br>LNPEPmirna:hsa-miR-3124-3pAILANI<br>LNPEPmirna:hsa-miR-329-3pAILANI<br>LNPEPmirna:hsa-miR-3529-5pAILANI<br>LNPEPmirna:hsa-miR-361-5pAILANI<br>LNPEPmirna:hsa-miR-362-3pAILANI<br>LNPEPmirna:hsa-miR-3681-3pAILANI<br>LNPEPmirna:hsa-miR-3689a-5pAILANI<br>LNPEPmirna:hsa-miR-3689b-5pAILANI<br>LNPEPmirna:hsa-miR-3689eAILANI<br>LNPEPmirna:hsa-miR-3689fAILANI<br>LNPEPmirna:hsa-miR-379-5pAILANI<br>LNPEPmirna:hsa-miR-3941AILANI<br>LNPEPmirna:hsa-miR-4302AILANI<br>LNPEPmirna:hsa-miR-4451AILANI<br>LNPEPmirna:hsa-miR-4469AILANI<br>LNPEPmirna:hsa-miR-4524a-3pAILANI<br>LNPEPmirna:hsa-miR-4650-3pAILANI<br>LNPEPmirna:hsa-miR-4689AILANI<br>LNPEPmirna:hsa-miR-4697-3pAILANI<br>LNPEPmirna:hsa-miR-4704-3pAILANI<br>LNPEPmirna:hsa-miR-4720-3pAILANI<br>LNPEPmirna:hsa-miR-4797-5pAILANI<br>LNPEPmirna:hsa-miR-4804-3pAILANI<br>LNPEPmirna:hsa-miR-485-3pAILANI<br>LNPEPmirna:hsa-miR-5007-3pAILANI<br>LNPEPmirna:hsa-miR-512-3pAILANI<br>LNPEPmirna:hsa-miR-5196-3pAILANI<br>LNPEPmirna:hsa-miR-539-3pAILANI<br>LNPEPmirna:hsa-miR-548aj-5pAILANI<br>LNPEPmirna:hsa-miR-548awAILANI<br>LNPEPmirna:hsa-miR-548f-5pAILANI<br>LNPEPmirna:hsa-miR-548g-5pAILANI<br>LNPEPmirna:hsa-miR-548x-5pAILANI<br>LNPEPmirna:hsa-miR-5692aAILANI<br>LNPEPmirna:hsa-miR-5705AILANI<br>LNPEPmirna:hsa-miR-603AILANI<br>LNPEPmirna:hsa-miR-6131AILANI<br>LNPEPmirna:hsa-miR-636AILANI<br>LNPEPmirna:hsa-miR-6749-3pAILANI<br>LNPEPmirna:hsa-miR-6770-5pAILANI<br>LNPEPmirna:hsa-miR-6783-5pAILANI<br>LNPEPmirna:hsa-miR-6844AILANI<br>LNPEPmirna:hsa-miR-6858-5pAILANI<br>LNPEPmirna:hsa-miR-6868-3pAILANI<br>LNPEPmirna:hsa-miR-6888-5pAILANI<br>LNPEPmirna:hsa-miR-744-5pAILANI<br>LNPEPmirna:hsa-miR-8080AILANI<br>LNPEPmirna:hsa-miR-8485AILANI<br>LNPEPmirna:hsa-miR-877-5pAILANI |

| Protein / gene | INDRA drugs | Clinical trials/ TF analysis | AILANI |
|---|---|---|---|
| | | | AHRdrugbank:DB00379mexiletineAILANI |
| | | | AHRdrugbank:DB00393nimodipineAILANI |
| | | | AHRdrugbank:DB00499flutamideAILANI |
| | | | AHRdrugbank:DB01076atorvastatinAILANI |
| | | | AHRdrugbank:DB01097leflunomideAILANI |
| | | | AHRdrugbank:DB01404ginsengAILANI |
| | | | AHRmirna:hsa-let-7a-5pAILANI |
| | | | AHRmirna:hsa-let-7b-5pAILANI |
| | | | AHRmirna:hsa-let-7c-5pAILANI |
| | | | AHRmirna:hsa-let-7d-5pAILANI |
| | | | AHRmirna:hsa-let-7e-5pAILANI |
| | | | AHRmirna:hsa-let-7f-5pAILANI |
| | | | AHRmirna:hsa-let-7g-5pAILANI |
| | | | AHRmirna:hsa-let-7i-5pAILANI |
| | | | AHRmirna:hsa-miR-1197AILANI |
| | | | AHRmirna:hsa-miR-1227-3pAILANI |
| | | | AHRmirna:hsa-miR-1233-5pAILANI |
| | | | AHRmirna:hsa-miR-124-3pAILANI |
| | | | AHRmirna:hsa-miR-1258AILANI |
| | | | AHRmirna:hsa-miR-1273eAILANI |
| | | | AHRmirna:hsa-miR-1273h-5pAILANI |
| | | | AHRmirna:hsa-miR-130b-3pAILANI |
| | | | AHRmirna:hsa-miR-149-3pAILANI |
| | | | AHRmirna:hsa-miR-16-1-3pAILANI |
| | | | AHRmirna:hsa-miR-181a-5pAILANI |
| | | | AHRmirna:hsa-miR-2467-5pAILANI |
| | | | AHRmirna:hsa-miR-26a-5pAILANI |
| | | | AHRmirna:hsa-miR-30b-3pAILANI |
| | | | AHRmirna:hsa-miR-30c-1-3pAILANI |
| | | | AHRmirna:hsa-miR-30c-2-3pAILANI |
| | | | AHRmirna:hsa-miR-3117-3pAILANI |
| | | | AHRmirna:hsa-miR-3122AILANI |
| | | | AHRmirna:hsa-miR-3152-5pAILANI |
| | | | AHRmirna:hsa-miR-3689a-3pAILANI |
| | | | AHRmirna:hsa-miR-3689b-3pAILANI |
| | | | AHRmirna:hsa-miR-3689cAILANI |
| | | | AHRmirna:hsa-miR-377-5pAILANI |
| | | | AHRmirna:hsa-miR-383-3pAILANI |
| | | | AHRmirna:hsa-miR-383-5pAILANI |
| | | | AHRmirna:hsa-miR-3913-5pAILANI |
| | | | AHRmirna:hsa-miR-3927-3pAILANI |
| | | | AHRmirna:hsa-miR-4284AILANI |
| | | | AHRmirna:hsa-miR-4451AILANI |
| | | | AHRmirna:hsa-miR-4458AILANI |
| | | | AHRmirna:hsa-miR-4459AILANI |
| | | | AHRmirna:hsa-miR-4500AILANI |
| | | | AHRmirna:hsa-miR-450a-1-3pAILANI |
| | | | AHRmirna:hsa-miR-455-3pAILANI |
| | | | AHRmirna:hsa-miR-4645-3pAILANI |
| | | | AHRmirna:hsa-miR-4699-5pAILANI |
| | | | AHRmirna:hsa-miR-4728-5pAILANI |
| | | | AHRmirna:hsa-miR-4761-5pAILANI |
| | | | AHRmirna:hsa-miR-4774-3pAILANI |
| | | | AHRmirna:hsa-miR-4775AILANI |
| | | | AHRmirna:hsa-miR-545-5pAILANI |
| | | | AHRmirna:hsa-miR-550b-2-5pAILANI |
| | | | AHRmirna:hsa-miR-6086AILANI |
| | | | AHRmirna:hsa-miR-625-5pAILANI |
| | | | AHRmirna:hsa-miR-6499-3pAILANI |
| | | | AHRmirna:hsa-miR-6513-5pAILANI |
| | | | AHRmirna:hsa-miR-6516-5pAILANI |
| | | | AHRmirna:hsa-miR-665AILANI |
| | | | AHRmirna:hsa-miR-6778-5pAILANI |
| | | | AHRmirna:hsa-miR-6779-5pAILANI |
| | | | AHRmirna:hsa-miR-6780a-5pAILANI |
| | | | AHRmirna:hsa-miR-6785-5pAILANI |
| | AHR chebi:CHEBI:16236 ethanol INDRA (drugs) | | AHRmirna:hsa-miR-6788-5pAILANI |
| | AHR chebi:CHEBI:16412 lipopolysaccharide INDRA (drugs) | | AHRmirna:hsa-miR-6799-5pAILANI |
| | AHR chebi:CHEBI:27881 resveratrol INDRA (drugs) | | AHRmirna:hsa-miR-6807-5pAILANI |
| | AHR chebi:CHEBI:28119 2,3,7,8-tetrachlorodibenzodioxine INDRA (drugs) | | AHRmirna:hsa-miR-6831-5pAILANI |
| | AHR chebi:CHEBI:41879 dexamethasone INDRA (drugs) | | AHRmirna:hsa-miR-6840-3pAILANI |
| | AHR chebi:CHEBI:4806 (-)-epigallocatechin 3-gallate INDRA (drugs) | | AHRmirna:hsa-miR-6883-5pAILANI |
| | AHR chebi:CHEBI:6804 methacholine INDRA (drugs) | | AHRmirna:hsa-miR-7106-5pAILANI |
| | AHR chebi:CHEBI:76995 alpha-naphthoflavone INDRA (drugs) | | AHRmirna:hsa-miR-7977AILANI |
| | AHR chebi:CHEBI:77260 5-methoxy-2-\{[(4-methoxy-3,5-dimethylpyridin-2-yl)methyl]sulfinyl\}-1h-benzimidazole INDRA (drugs) | | AHRmirna:hsa-miR-887-5pAILANI |
| **AHR** | AHR chebi:CHEBI:9168 sirolimus INDRA (drugs) | | AHRmirna:hsa-miR-98-5pAILANI |

| Protein / gene | INDRA drugs | Clinical trials/ TF analysis | AILANI |
|---|---|---|---|
| **MAS1** | MAS1chebi:CHEBI:15355acetylcholineINDRA (drugs)<br>MAS1chebi:CHEBI:16134ammoniaINDRA (drugs)<br>MAS1chebi:CHEBI:16680s-adenosyl-l-homocysteineINDRA (drugs)<br>MAS1chebi:CHEBI:18421superoxideINDRA (drugs)<br>MAS1chebi:CHEBI:280825alpha-cholestane-3beta,5,6beta-triolINDRA (drugs) | | |
| **RBX1** | RBX1pubchem.compound:10985033rccINDRA (drugs) RBX1 chebi:CHEBI:30778 gallic acid INDRA (drugs)<br>RBX1 chebi:CHEBI:31823 mercury dichloride INDRA (drugs)<br>RBX1 chebi:CHEBI:63791 lenalidomide INDRA (drugs)<br>RBX1 chebi:CHEBI:72690 pomalidomide INDRA (drugs)<br>RBX1 chebi:CHEBI:74947 2-(2,6-dioxopiperidin-3-yl)-1h-isoindole-1,3(2h)-dione INDRA (drugs) | | RBX1mirna:hsa-miR-188-3pAILANI<br>RBX1mirna:hsa-miR-194-5pAILANI<br>RBX1mirna:hsa-miR-2276-5pAILANI<br>RBX1mirna:hsa-miR-3136-5pAILANI<br>RBX1mirna:hsa-miR-4439AILANI<br>RBX1mirna:hsa-miR-4513AILANI<br>RBX1mirna:hsa-miR-5187-3pAILANI<br>RBX1mirna:hsa-miR-642a-5pAILANI<br>RBX1mirna:hsa-miR-6855-3pAILANI<br>RBX1mirna:hsa-miR-6857-3pAILANI<br>RBX1mirna:hsa-miR-6875-3pAILANI<br>RBX1mirna:hsa-miR-6892-3pAILANI<br>RBX1mirna:hsa-miR-92a-3pAILANI |
| **KEAP1** | KEAP1chebi:CHEBI:15948lycopeneINDRA (drugs)<br>KEAP1chebi:CHEBI:16243quercetinINDRA (drugs)<br>KEAP1chebi:CHEBI:26523reactive oxygen speciesINDRA (drugs)<br>KEAP1chebi:CHEBI:27881resveratrolINDRA (drugs)<br>KEAP1chebi:CHEBI:31697indolin-2-oneINDRA (drugs)<br>KEAP1chebi:CHEBI:3259cccpINDRA (drugs)<br>KEAP1chebi:CHEBI:76004dimethyl fumarateINDRA (drugs)<br>KEAP1chebi:CHEBI:777332-(3,4-dimethoxyphenyl)-5-\{[2-(3,4-dimethoxyphenyl)ethyl](methyl)amino\}-2-(propan-2-yl)pentanenitrileINDRA (drugs)<br>KEAP1chebi:CHEBI:8198trans-piceidINDRA (drugs)<br>KEAP1drugbank:DB05983bardoxolone methylINDRA (drugs) | | KEAP1drugbank:DB08908dimethyl fumarateAILANI<br>KEAP1mirna:hsa-miR-141-3pAILANI<br>KEAP1mirna:hsa-miR-200a-3pAILANI<br>KEAP1mirna:hsa-miR-26b-5pAILANI<br>KEAP1mirna:hsa-miR-7-5pAILANI<br>KEAP1mirna:hsa-miR-92b-3pAILANI |

| Protein / gene | INDRA drugs | Clinical trials/ TF analysis | AILANI |
| --- | --- | --- | --- |
| CUL3 | CUL3 chebi:CHEBI:145535 pevonedistat INDRA (drugs)<br>CUL3 chebi:CHEBI:16136 hydrogen sulfide INDRA (drugs)<br>CUL3 chebi:CHEBI:50845 doxycycline INDRA (drugs)<br>CUL3 chebi:CHEBI:63637 vemurafenib INDRA (drugs) | | CUL3mirna:hsa-let-7b-5pAILANI<br>CUL3mirna:hsa-miR-10a-5pAILANI<br>CUL3mirna:hsa-miR-1225-3pAILANI<br>CUL3mirna:hsa-miR-1233-3pAILANI<br>CUL3mirna:hsa-miR-1265AILANI<br>CUL3mirna:hsa-miR-1266-3pAILANI<br>CUL3mirna:hsa-miR-130a-3pAILANI<br>CUL3mirna:hsa-miR-130b-3pAILANI<br>CUL3mirna:hsa-miR-15a-5pAILANI<br>CUL3mirna:hsa-miR-15b-5pAILANI<br>CUL3mirna:hsa-miR-16-5pAILANI<br>CUL3mirna:hsa-miR-18a-3pAILANI<br>CUL3mirna:hsa-miR-192-5pAILANI<br>CUL3mirna:hsa-miR-195-5pAILANI<br>CUL3mirna:hsa-miR-203a-3pAILANI<br>CUL3mirna:hsa-miR-218-5pAILANI<br>CUL3mirna:hsa-miR-301a-3pAILANI<br>CUL3mirna:hsa-miR-301b-3pAILANI<br>CUL3mirna:hsa-miR-302a-3pAILANI<br>CUL3mirna:hsa-miR-302b-3pAILANI<br>CUL3mirna:hsa-miR-302c-3pAILANI<br>CUL3mirna:hsa-miR-302d-3pAILANI<br>CUL3mirna:hsa-miR-302eAILANI<br>CUL3mirna:hsa-miR-3120-3pAILANI<br>CUL3mirna:hsa-miR-3177-5pAILANI<br>CUL3mirna:hsa-miR-339-5pAILANI<br>CUL3mirna:hsa-miR-3529-5pAILANI<br>CUL3mirna:hsa-miR-3613-5pAILANI<br>CUL3mirna:hsa-miR-3666AILANI<br>CUL3mirna:hsa-miR-372-3pAILANI<br>CUL3mirna:hsa-miR-373-3pAILANI<br>CUL3mirna:hsa-miR-379-5pAILANI<br>CUL3mirna:hsa-miR-424-5pAILANI<br>CUL3mirna:hsa-miR-4295AILANI<br>CUL3mirna:hsa-miR-4451AILANI<br>CUL3mirna:hsa-miR-4477aAILANI<br>CUL3mirna:hsa-miR-4524a-5pAILANI<br>CUL3mirna:hsa-miR-4524b-5pAILANI<br>CUL3mirna:hsa-miR-4536-5pAILANI<br>CUL3mirna:hsa-miR-454-3pAILANI<br>CUL3mirna:hsa-miR-4635AILANI<br>CUL3mirna:hsa-miR-4650-3pAILANI<br>CUL3mirna:hsa-miR-4668-3pAILANI<br>CUL3mirna:hsa-miR-4705AILANI<br>CUL3mirna:hsa-miR-4764-5pAILANI<br>CUL3mirna:hsa-miR-497-5pAILANI<br>CUL3mirna:hsa-miR-503-5pAILANI<br>CUL3mirna:hsa-miR-511-3pAILANI<br>CUL3mirna:hsa-miR-5197-3pAILANI<br>CUL3mirna:hsa-miR-519a-3pAILANI<br>CUL3mirna:hsa-miR-519b-3pAILANI<br>CUL3mirna:hsa-miR-519c-3pAILANI<br>CUL3mirna:hsa-miR-520a-3pAILANI<br>CUL3mirna:hsa-miR-520bAILANI<br>CUL3mirna:hsa-miR-520c-3pAILANI<br>CUL3mirna:hsa-miR-520d-3pAILANI<br>CUL3mirna:hsa-miR-520eAILANI<br>CUL3mirna:hsa-miR-591AILANI<br>CUL3mirna:hsa-miR-646AILANI<br>CUL3mirna:hsa-miR-670-5pAILANI<br>CUL3mirna:hsa-miR-6734-3pAILANI<br>CUL3mirna:hsa-miR-6807-5pAILANI<br>CUL3mirna:hsa-miR-6838-5pAILANI<br>CUL3mirna:hsa-miR-99a-5pAILANI |

| Protein / gene | INDRA drugs | Clinical trials/ TF analysis | AILANI |
|---|---|---|---|
| HMOX1 | HMOX1 chebi:CHEBI:15430 protoporphyrin INDRA (drugs)<br>HMOX1 chebi:CHEBI:16412 lipopolysaccharide INDRA (drugs)<br>HMOX1 chebi:CHEBI:17245 carbon monoxide INDRA (drugs)<br>HMOX1 chebi:CHEBI:18248 iron atom INDRA (drugs)<br>HMOX1 chebi:CHEBI:26523 reactive oxygen species INDRA (drugs)<br>HMOX1 chebi:CHEBI:27007 tin atom INDRA (drugs)<br>HMOX1 chebi:CHEBI:28783 zinc protoporphyrin INDRA (drugs)<br>HMOX1 chebi:CHEBI:7421 nac INDRA (drugs) HMOX1 mesh:C032628 tin protoporphyrin ix INDRA (drugs) LNPEPpubchem.compound:53 | | HMOX1 drugbank:DB00157 nadh AILANI<br>HMOX1 drugbank:DB01942 formic acid AILANI<br>HMOX1 drugbank:DB02073 biliverdine ix alpha AILANI<br>HMOX1 drugbank:DB02468 12-phenylheme AILANI<br>HMOX1 drugbank:DB03906 2-phenylheme AILANI<br>HMOX1 drugbank:DB04912 stannsoporfin AILANI<br>HMOX1 drugbank:DB06914 1-({2-[2-(4-chlorophenyl)ethyl]-1,3-dioxolan-2-yl}methyl)-1h-imidazole AILANI<br>HMOX1 drugbank:DB07342 1-(adamantan-1-yl)-2-(1h-imidazol-1-yl)ethanone AILANI HMOX1 mirna:hsa-miR-1-3p AILANI<br>HMOX1 mirna:hsa-miR-122-5p AILANI<br>HMOX1 mirna:hsa-miR-124-3p AILANI<br>HMOX1 mirna:hsa-miR-128-3p AILANI<br>HMOX1 mirna:hsa-miR-1291 AILANI<br>HMOX1 mirna:hsa-miR-148b-3p AILANI<br>HMOX1 mirna:hsa-miR-16-5p AILANI<br>HMOX1 mirna:hsa-miR-196a-5p AILANI<br>HMOX1 mirna:hsa-miR-216a-3p AILANI<br>HMOX1 mirna:hsa-miR-24-3p AILANI<br>HMOX1 mirna:hsa-miR-26b-5p AILANI<br>HMOX1 mirna:hsa-miR-3126-5p AILANI<br>HMOX1 mirna:hsa-miR-3135b AILANI<br>HMOX1 mirna:hsa-miR-3180-5p AILANI<br>HMOX1 mirna:hsa-miR-335-5p AILANI<br>HMOX1 mirna:hsa-miR-3612 AILANI<br>HMOX1 mirna:hsa-miR-3652 AILANI<br>HMOX1 mirna:hsa-miR-3667-3p AILANI<br>HMOX1 mirna:hsa-miR-3681-3p AILANI<br>HMOX1 mirna:hsa-miR-4430 AILANI<br>HMOX1 mirna:hsa-miR-4443 AILANI<br>HMOX1 mirna:hsa-miR-4446-3p AILANI<br>HMOX1 mirna:hsa-miR-4448 AILANI<br>HMOX1 mirna:hsa-miR-4492 AILANI<br>HMOX1 mirna:hsa-miR-4498 AILANI<br>HMOX1 mirna:hsa-miR-4691-5p AILANI<br>HMOX1 mirna:hsa-miR-4721 AILANI<br>HMOX1 mirna:hsa-miR-5001-5p AILANI<br>HMOX1 mirna:hsa-miR-5008-5p AILANI<br>HMOX1 mirna:hsa-miR-5193 AILANI<br>HMOX1 mirna:hsa-miR-520a-5p AILANI<br>HMOX1 mirna:hsa-miR-525-5p AILANI<br>HMOX1 mirna:hsa-miR-5590-5p AILANI<br>HMOX1 mirna:hsa-miR-650 AILANI<br>HMOX1 mirna:hsa-miR-660-3p AILANI<br>HMOX1 mirna:hsa-miR-6734-3p AILANI<br>HMOX1 mirna:hsa-miR-6749-3p AILANI<br>HMOX1 mirna:hsa-miR-6758-5p AILANI<br>HMOX1 mirna:hsa-miR-6775-3p AILANI<br>HMOX1 mirna:hsa-miR-6792-3p AILANI<br>HMOX1 mirna:hsa-miR-6856-5p AILANI<br>HMOX1 mirna:hsa-miR-6873-5p AILANI<br>HMOX1 mirna:hsa-miR-6875-5p AILANI<br>HMOX1 mirna:hsa-miR-762 AILANI<br>HMOX1 mirna:hsa-miR-7845-5p AILANI<br>HMOX1 mirna:hsa-miR-7976 AILANI |
| ALG5 | | | ALG5mirna:hsa-miR-24-3pAILANI |

| Protein / gene | INDRA drugs | Clinical trials/ TF analysis | AILANI |
|---|---|---|---|
| ARL6IP6 | | | ARL6IP6mirna:hsa-let-7a-5pAILANI<br>ARL6IP6mirna:hsa-miR-1260aAILANI<br>ARL6IP6mirna:hsa-miR-1260bAILANI<br>ARL6IP6mirna:hsa-miR-132-3pAILANI<br>ARL6IP6mirna:hsa-miR-142-3pAILANI<br>ARL6IP6mirna:hsa-miR-181a-5pAILANI<br>ARL6IP6mirna:hsa-miR-188-3pAILANI<br>ARL6IP6mirna:hsa-miR-205-3pAILANI<br>ARL6IP6mirna:hsa-miR-3140-5pAILANI<br>ARL6IP6mirna:hsa-miR-3156-3pAILANI<br>ARL6IP6mirna:hsa-miR-32-3pAILANI<br>ARL6IP6mirna:hsa-miR-329-3pAILANI<br>ARL6IP6mirna:hsa-miR-362-3pAILANI<br>ARL6IP6mirna:hsa-miR-374c-3pAILANI<br>ARL6IP6mirna:hsa-miR-3941AILANI<br>ARL6IP6mirna:hsa-miR-3942-3pAILANI<br>ARL6IP6mirna:hsa-miR-4293AILANI<br>ARL6IP6mirna:hsa-miR-4301AILANI<br>ARL6IP6mirna:hsa-miR-4427AILANI<br>ARL6IP6mirna:hsa-miR-4760-3pAILANI<br>ARL6IP6mirna:hsa-miR-4760-5pAILANI<br>ARL6IP6mirna:hsa-miR-5100AILANI<br>ARL6IP6mirna:hsa-miR-5696AILANI<br>ARL6IP6mirna:hsa-miR-579-3pAILANI<br>ARL6IP6mirna:hsa-miR-596AILANI<br>ARL6IP6mirna:hsa-miR-603AILANI<br>ARL6IP6mirna:hsa-miR-664a-3pAILANI<br>ARL6IP6mirna:hsa-miR-664b-3pAILANI<br>ARL6IP6mirna:hsa-miR-8061AILANI<br>ARL6IP6mirna:hsa-miR-8485AILANI<br>ARL6IP6uniprot:P36895AILANI |
| CASP7 | CASP7 chebi:CHEBI:15746 spermine INDRA (drugs)<br>CASP7 chebi:CHEBI:16509 1,4-benzoquinone INDRA (drugs)<br>CASP7 chebi:CHEBI:16796 melatonin INDRA (drugs)<br>CASP7 chebi:CHEBI:18388 apigenin INDRA (drugs)<br>CASP7 chebi:CHEBI:27363 zinc atom INDRA (drugs)<br>CASP7 chebi:CHEBI:27899 cisplatin INDRA (drugs)<br>CASP7 chebi:CHEBI:59385 ac-asp-glu-val-asp-h INDRA (drugs)<br>CASP7 chebi:CHEBI:7421 nac INDRA (drugs)<br>CASP7 drugbank:DB03384 fica INDRA (drugs), AILANI<br>CASP7 drugbank:DB05408 emricasan INDRA (drugs) | | CASP7mirna:hsa-miR-106a-5pAILANI<br>CASP7mirna:hsa-miR-106b-5pAILANI<br>CASP7mirna:hsa-miR-122-5pAILANI<br>CASP7mirna:hsa-miR-132-3pAILANI<br>CASP7mirna:hsa-miR-192-5pAILANI<br>CASP7mirna:hsa-miR-215-5pAILANI<br>CASP7mirna:hsa-miR-26b-5pAILANI<br>CASP7mirna:hsa-miR-484AILANI |
| EGFR | | EGFR zanubrutinib drugbank:DB15035 Clinical Trials DB<br>EGFR abivertinib drugbank:DB15327 Clinical Trials DB | |
